# Personalized biventricular mechanics and sensitivity to model morphology

**DOI:** 10.64898/2025.12.11.693778

**Authors:** Aaron L. Brown, Lei Shi, Matteo Salvador, Fanwei Kong, Daniel B. Ennis, Ian Chen, Vijay Vedula, Alison L. Marsden

## Abstract

We present a computational framework for constructing patient-specific models of cardiac mechanics based on standard clinical data, including electrocardiogram (ECG), cuff blood pressure, and electrocardiography-gated computed tomography angiography (CTA) imaging. The model is coupled to a closed-loop lumped parameter network (LPN) circulatory model and incorporates rule-based fiber architecture, as well as spatially varying epicardial boundary conditions to approximate surrounding tissue support. Model parameters are personalized through a multistep procedure that sequentially tunes circulatory dynamics, passive mechanics, and active contraction. The resulting personalized BiV model closely matches clinical pressure and volume measurements and reasonably agrees with image-based myocardial deformation. To assess the impact of anatomical model choice, we compare the BiV model to two commonly-used simplifications: a truncated BiV (t-BiV) model cut at the basal plane and a left ventricle-only (LV) model. For these models, we also evaluate their sensitivity to plausible variations in boundary conditions and contractile strength. With all other inputs held fixed, the LV model exhibits similar global pressure/volume behavior, despite moderate differences in regional deformation. In contrast, the t-BiV model produces substantial differences in both global function and local myocardial mechanics. These results suggest that while LV-only models may be sufficient for biomechanical studies, truncation at the basal plane strongly impacts model outputs and should be used with caution.

## 1 Introduction

Cardiac computational models have incredible potential for improving our understanding of the heart in both health and disease. Simulations have already been applied in specific clinical contexts, including evaluating pacing strategies [1], optimizing cardiac resynchronization therapy [2, 3], identifying optimal targets for ablation therapy in arrhythmias [4, 5], assessing implantable device performance [6, 7], guiding surgical decision-making in congenital heart disease [8, 9], and enabling virtual drug trials to anticipate electromechanical disturbances [10, 11]. Models also provide a framework for studying disease processes that are difficult to probe directly. In hypertrophic cardiomyopathy, models have clarified the role of fibrosis on arrhythmia risk [12] and quantified the effect of altered tissue characteristics on ventricular dysfunction [13]. Multiphysics models have revealed how atrial fibrosis promotes stroke [14], and explored subcellular and metabolic contributions to heart failure [15]. Meanwhile, developmental models have elucidated the structural roles of trabeculae in embryonic hearts [16] as well the mechanobiologic pathways underpinning trabecular formation [17].

The clinical utility of these models is greatly enhanced when they are personalized to patient-specific data [18]. Personalization typically involves reconstructing a patient’s anatomy from medical imaging, a process that is labor-intensive [19] but has been accelerated by recent advances in machine learning methods [20, 21, 22, 23, 24, 25, 26]. Beyond anatomy, personalization also requires calibrating dozens of model parameters to match clinical measurements, a task complicated by the high computational cost of simulations and the inherent complexity of cardiac models [18]. Due to the diversity of models, available data, and clinical applications, no standardized parameter-tuning method exists, and manual approaches are still commonly used [2, 27, 28]. Recent work has explored a range of strategies, including sensitivity analyses to identify influential parameters [29, 30], which can inform novel optimization methods [31, 32], and acceleration techniques, including reduced-order surrogate modeling [33, 34, 35], neural networks surrogate modeling [30, 36, 37, 38, 39], and GPU-based computation [40]. At present, many image-based models can adequately capture global features like chamber volumes. However, relatively few studies have compared the predicted myocardial motion in these models against time-resolved imaging data, and in those cases, local discrepancies often remain [27, 32, 41, 42]. Bridging this gap remains a key challenge in cardiac mechanics modeling.

As cardiac modeling frameworks grow more sophisticated, a parallel question has emerged: how much anatomical detail is necessary to achieve physiologically meaningful results? Anatomical models used in cardiac simulations vary widely in complexity, ranging from idealized univentricular representations to four-chamber geometries [43]. Whole-heart models, which include all four chambers and sometimes the roots of the aorta and pulmonary arteries, offer the most physiologically complete representation of cardiac anatomy [27, 32, 41, 44, 45, 46, 47, 48]. However, constructing patient-specific whole-heart models remains challenging due to limited image resolution, especially when modeling thin-walled structures such as the right ventricle (RV), atria, atrioventricular regions, outflow tracts, and cardiac valves [49]. Moreover, whole-heart simulations are computationally expensive and more challenging to calibrate [50], and patient-specific whole-heart models remain rare [32, 51].

To reduce complexity, many studies still employ biventricular models that include only the LV and RV [34, 52, 53, 54]. These are often further simplified by truncating the geometry at the basal plane [33, 50, 55, 56, 57, 58], thereby removing the thinner and more complex near-valve regions of the ventricles. The simplest commonly used models focus on the LV alone [59, 60], sometimes similarly truncated [49, 61, 62, 63]. These simplifications are motivated in part by the practical difficulties of reconstructing detailed multi-chamber geometries from coarse imaging data (e.g., MRI or ultrasound) and of defining suitable models (e.g., boundary conditions or myofiber orientations) when simulating such complex structures, and partly by the view that the ventricles, especially the LV, dominate the heart’s mechanical function. However, this assumption may overlook important physiological phenomena and, critically, influence the conclusions drawn from such simulations.

Indeed, the various structures of the heart interact electrically, mechanically, and hemodynamically [64, 65]. So-called “ventricular interdependence” is mediated by the shared ventricular septum, the pericardium, and blood flow through the closed-loop circulatory system [66, 67], with both LV-to-RV and RV-to-LV interactions becoming especially important in disease states [68, 69]. For example, RV dysfunction can reduce LV filling and mimic diastolic dysfunction [70], while LV assist devices (LVADs) can shift RV pressure-volume (PV) loops and alter septal curvature [71]. Similarly, the atria and ventricles are mechanically and hemodynamically coupled across the atrioventricular interface and valves [46, 72, 73]. Ventricular contraction contributes to atrial filling, an effect enhanced by the presence of the pericardium [41], while atrial contraction enhances ventricular preload, although its quantitative importance is debated [74, 75]. Further, even passive atrial tissue can influence ventricular deformation due to inertial effects [76].

As Rodero et al. note, “even subtle changes in cardiac anatomy can have a large impact on cardiac function” [77]; however, few studies have directly examined the effect of excluding certain cardiac structures from a model. One example is the work of Palit et al. [78], who compared diastolic mechanics in LV and BiV models constructed from the same data. They found that the RV modestly enhances LV diastolic filling. Although RV pressure on the interventricular septum tends to hinder LV filling, pressure on the RV free wall promotes it, and the latter effect is stronger. Nevertheless, a more thorough understanding of how anatomical model choices influence simulation outcomes over the entire cardiac cycle is needed to guide modelers in selecting anatomies that balance model efficiency with physiological fidelity. This study aims to fill that gap.

Further, the selection of boundary conditions (BCs) plays an equally critical role in shaping cardiac function [79] and is inherently linked to the choice of anatomical model. For example, in models that truncate the ventricles at the basal plane, the modeler must impose a mechanical BC on this artificial boundary that replicates, to some extent, the influence of the omitted structures. Inappropriate or overly simplistic basal BCs can lead to unphysiological constraints on ventricular motion, such as atrioventricular plane displacement, which is a key contributor to stroke volume [41]. While early approaches adopted a fixed basal plane [80, 81, 82] or restricted motion along individual axes [83, 84], later studies introduced more realistic BCs, such as constraining only average in-plane motion [50], applying Robin-type spring-dashpot conditions [33, 52, 85], or using energy-consistent formulations that account for the effect of blood pressure on the omitted structures [86, 87].

Epicardial BCs are similarly important. Physiologically, the heart mechanically interacts with the pericardium and surrounding thoracic anatomy, which constrain epicardial normal motion while allowing tangential sliding. Some studies have modeled this interaction directly using contact mechanics and separate pericardial meshes [41, 88], but more commonly, researchers apply a mixed Robin-type boundary condition that penalizes normal displacement while permitting tangential motion [27]. These BCs have been shown to improve alignment with image-derived motion data and support more realistic atrioventricular coupling and filling dynamics, especially when spatially varying pericardial support is modeled [42, 46, 47, 89, 90]. However, current models still struggle to reproduce detailed myocardial deformation patterns observed in image data, a limitation that could be addressed by more accurate epicardial boundary conditions. Towards this end, Jilberto & Nordsletten [91] recently proposed modeling the localized forces exerted by the diaphragm and ribs on the RV, and found greater accuracy in estimating the reference configuration of the heart. As with anatomical detail, the impact of boundary conditions is often underappreciated but is critical for producing physiologically relevant simulations.

In this paper, we present a patient-specific biventricular (BiV) mechanics model constructed from patient CTA images. We outline a multistep personalization procedure, adapted from the one developed by Shi et al. for the left atrium [92], and demonstrate that the model accurately reproduces the patient’s cuff blood pressure measurements and phase-resolved chamber volumes, with reasonable agreement in myocardial deformation. To assess the impact of anatomical complexity on simulation results, we compare the BiV model to two common alternatives: a truncated BiV (t-BiV) model excluding tissue above the basal plane, and an LV-only model excluding the RV. Importantly, all parameters are kept fixed, isolating the effect of geometry; for the simplified models, we explore reasonable variations in boundary conditions and contractile strength. These comparisons reveal how anatomical choices influence the fidelity and interpretability of cardiac mechanics model outputs, within a personalized and physiologically relevant setting.

The remainder of the paper is organized as follows. Section 2 describes our patient-specific modeling pipeline, including patient data collected for this study (Section 2.1), construction of the patient-specific geometric models (Section 2.2), myocardial mechanics formulation and simulation details (Section 2.3), multistep personalization procedure (Section 2.4), and simulation details for the anatomically simplified models (Section 2.5). Results are presented in Section 3 and discussed in Section 4. In particular, we demonstrate the effectiveness of our multistep personalization method and reveal the effect of geometry on simulation results, in terms of hemodynamic pressures and volumes, as well as local myocardial deformation. We summarize our work and principal findings in Section 5.

## 2 Methods

A graphical summary of this study is given in Figure 1. The first part of this work involves constructing a patient-specific model of BiV mechanics. We begin by collecting patient data, which includes phase-resolved gated computed tomography angiography (CTA) images, an ECG waveform, and cuff blood pressure measurements. Then, we create a multiscale finite element (FE) model, complete with rule-based fiber directions [93, 94] and physiological boundary conditions [42], and coupled to a closed-loop 0D circulation model [85, 87]. A multistep personalization strategy is then used to tune the circulation parameters, passive material properties, and active contraction parameters [92]. We evaluate the personalized BiV model by its ability to replicate myocardial motion observed in the CTA data, match key clinical metrics, and produce physiological patterns of tissue stress and strain. In the second part of this work, we explore the effect of geometry on model outputs. Considering the BiV model as a baseline, we simulate truncated BiV (t-BiV) and LV-only (LV) geometries, keeping all other model inputs identical and allowing reasonable variations in boundary conditions and contractile strength.

**Figure 1.**
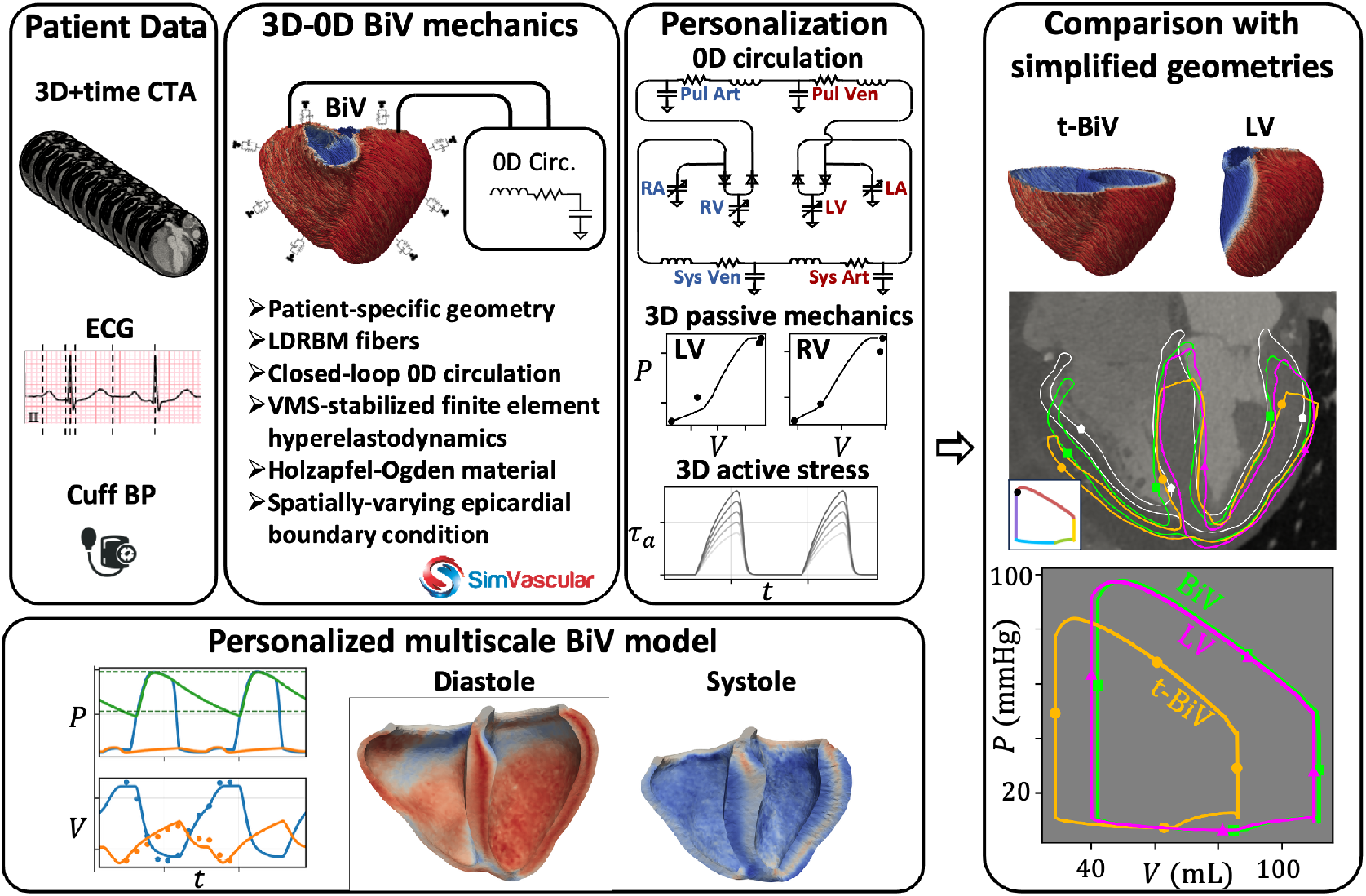
Graphical abstract. We first develop a personalized model of biventricular (BiV) mechanics. This involves three major steps: 1) collecting patient data, including gated computed tomography angiography (CTA) images, electrocardiogram (ECG), and cuff blood pressure (BP); 2) constructing a multiscale model of BiV mechanics; and 3) personalizing the model to recapitulate the collected patient data. The personalized model is then evaluated in terms of agreement with image data and clinical metrics, as well as tissue stress and fiber strains. In the second part of this study, we compare the personalized BiV model with two simplified geometries: a truncated BiV model (t-BiV) and a left ventricle-only (LV) model, keeping all other model inputs identical. For these models, we also evaluate their sensitivity to plausible variations in boundary conditions and contractile strength. LDRBM: Laplace-Dirichlet Rule-Based method; VMS: variational multiscale; LA: left atrium; RA: right atrium; LV: left ventricle; RV: right ventricle; Sys: systemic, Pul: pulmonary, Art: arterial, Ven: venous.

### 2.1 Patient data

Clinical data is obtained from a 50-year-old male patient with mild coronary artery disease. ECG-gated CTA for this patient yields 3D volumetric images at 10 phases of the cardiac cycle, sampled at every 10% of the RR interval (Figure 2). The CTA volumes contain 512 *×* 512 *×* 213 voxels, and each voxel’s resolution is 0.39 *×* 0.39 *×* 0.7 mm^3^. Using MeshDeformNet, a deep-learning-based whole-heart mesh reconstruction approach [22], the images are automatically segmented to track the motion of the major cardiac structures over the cardiac cycle. Volumes for the LV, RV, and right atrium (RA) are calculated from these moving meshes, while for the left atrium (LA), a manual segmentation is used to calculate its cavity volume, since MeshDeformNet does not suitably capture the left atrial appendage (Table 1). Additional clinical measurements, including cuff blood pressures and an ECG waveform, are also obtained for this individual (Table 2). All data collection procedures are approved by an Institutional Review Board (IRB) and conducted in compliance with HIPAA regulations. More details about the data collection protocol can be found in Chen et al. [99]. In addition to measured data, we also select reference values for certain circulatory pressures, which we found necessary to adequately constrain the model parameterization (Table 3), discussed in Section 2.4.1.

**Table 1.** Calculated chamber volumes (mL) for each cardiac chamber at 10 phases of the cardiac cycle. Volumes for the left atrium (LA) are calculated manually to properly account for the LA appendage, while volumes for the other chambers are calculated from an automatic segmentation of the CTA data (Figure 2). Minimum and maximum volumes for each chamber are bolded.

|  | RR0% | RR10% | RR20% | RR30% | RR40% | RR50% | RR60% | RR70% | RR80% | RR90% |
| --- | --- | --- | --- | --- | --- | --- | --- | --- | --- | --- |
| LA | **38.2** | 47.5 | 54.4 | 62.1 | 68.9 | **72.7** | 63.3 | 60.0 | 58.8 | 43.9 |
| RA | 61.1 | 70.6 | 83.6 | 93.3 | 101.1 | **107.9** | 101.8 | 93.7 | 84.1 | **56.0** |
| LV | **115.8** | 99.6 | 62.8 | 39.8 | **38.1** | 43.5 | 68.2 | 83.0 | 92.5 | 115.2 |
| RV | **173.5** | 156.4 | 120.6 | 96.5 | **87.7** | 91.1 | 107.9 | 123.0 | 138.0 | 172.1 |

**Table 2.** Additional clinical measurements obtained for the studied patient. PR interval: time from the start of the P-wave to the start of the QRS complex

| Cuff blood pressure (sys/dias) | Heart rate | PR interval |
| --- | --- | --- |
| 98/53 mmHg | 87 BPM | 182 ms |

**Table 3.** Literature-based reference pressure values are used to constrain model parameterization, ensuring physiological circulatory dynamics. PAP: pulmonary arterial pressure; CVP: central venous pressure (approximately mean right atrial pressure); PAWP: pulmonary arterial wedge pressure (approximately mean left atrial pressure); PVP: peripheral venous pressure (approximately CVP + 2 mmHg [96]).

| PAP (sys/dias) | CVP | PAWP | PVP |
| --- | --- | --- | --- |
| $20 \pm 5 / 11.5 \pm 3.5$ mmHg [97] | $4 \pm 4$ mmHg [98] | $7 \pm 5$ mmHg [98] | $6 \pm 4$ mmHg [96] |

**Figure 2.**
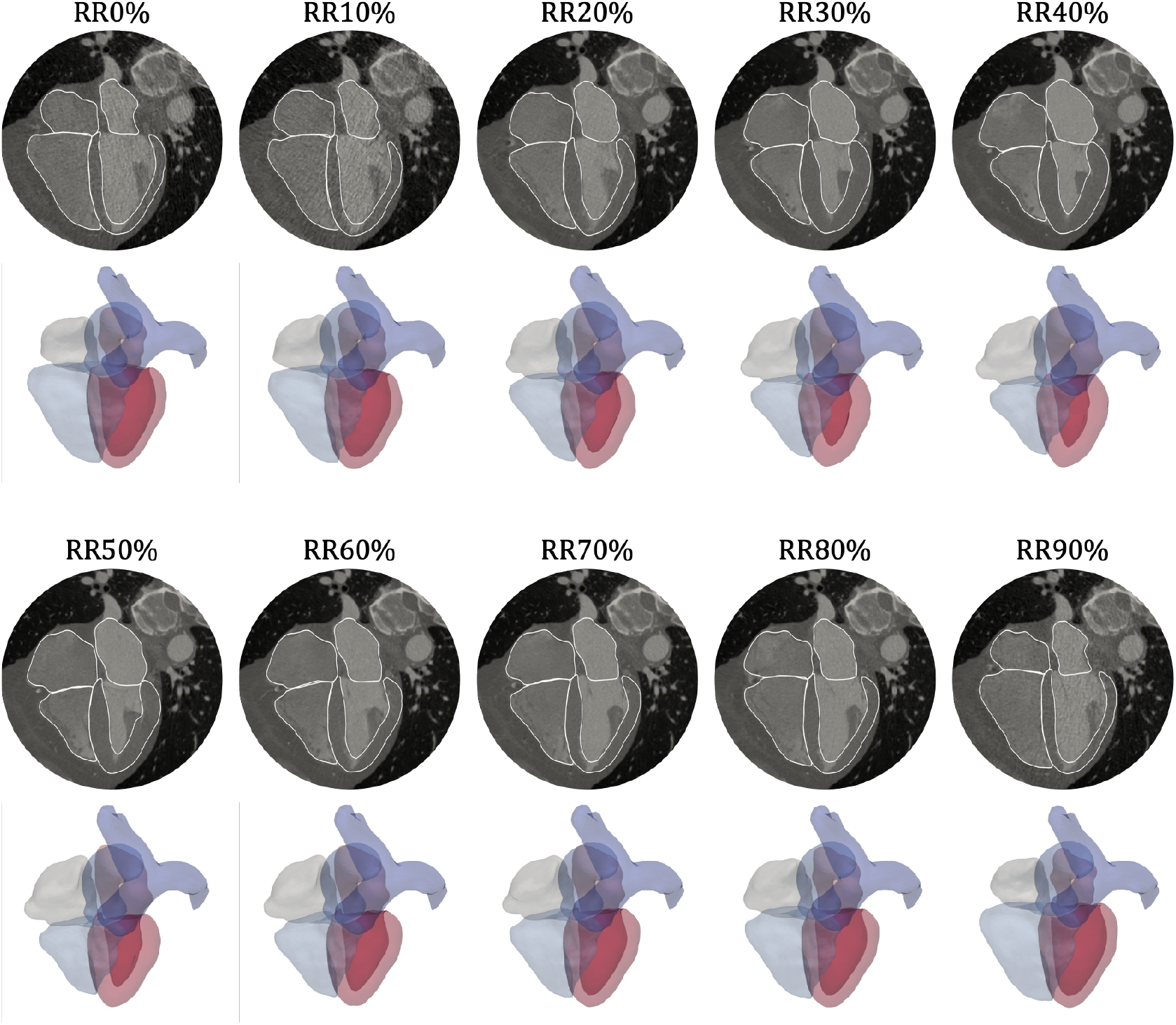
Phase-resolved ECG-gated CTA images were obtained at 10 time points over the cardiac cycle. Images are shown in the 4-chamber slice view, defined by a plane containing the apex, mitral valve center, and tricuspid valve center [95]. These images are automatically segmented using a deep-learning method [22] (white boundary in 4-chamber views and transparent 3D models). The method segments the blood pools of the LA, RA, and RV, as well as the myocardium of the LV. The trunks of the aorta and pulmonary arteries are also segmented. RRx% denotes the percentage of the cardiac cycle duration, starting at the R wave on an ECG. RR0% is approximately end-diastole, and RR40% is approximately end-systole.

### 2.2 Anatomical model construction

Three ventricular myocardial models – a biventricle (BiV), a truncated biventricle (t-BiV), and an isolated left ventricle (LV) – are constructed using a semiautomatic pipeline from the CTA volume at RR70% (diastasis). These models, visualized in Figure 3A-C, are constructed as follows.

**Figure 3.**
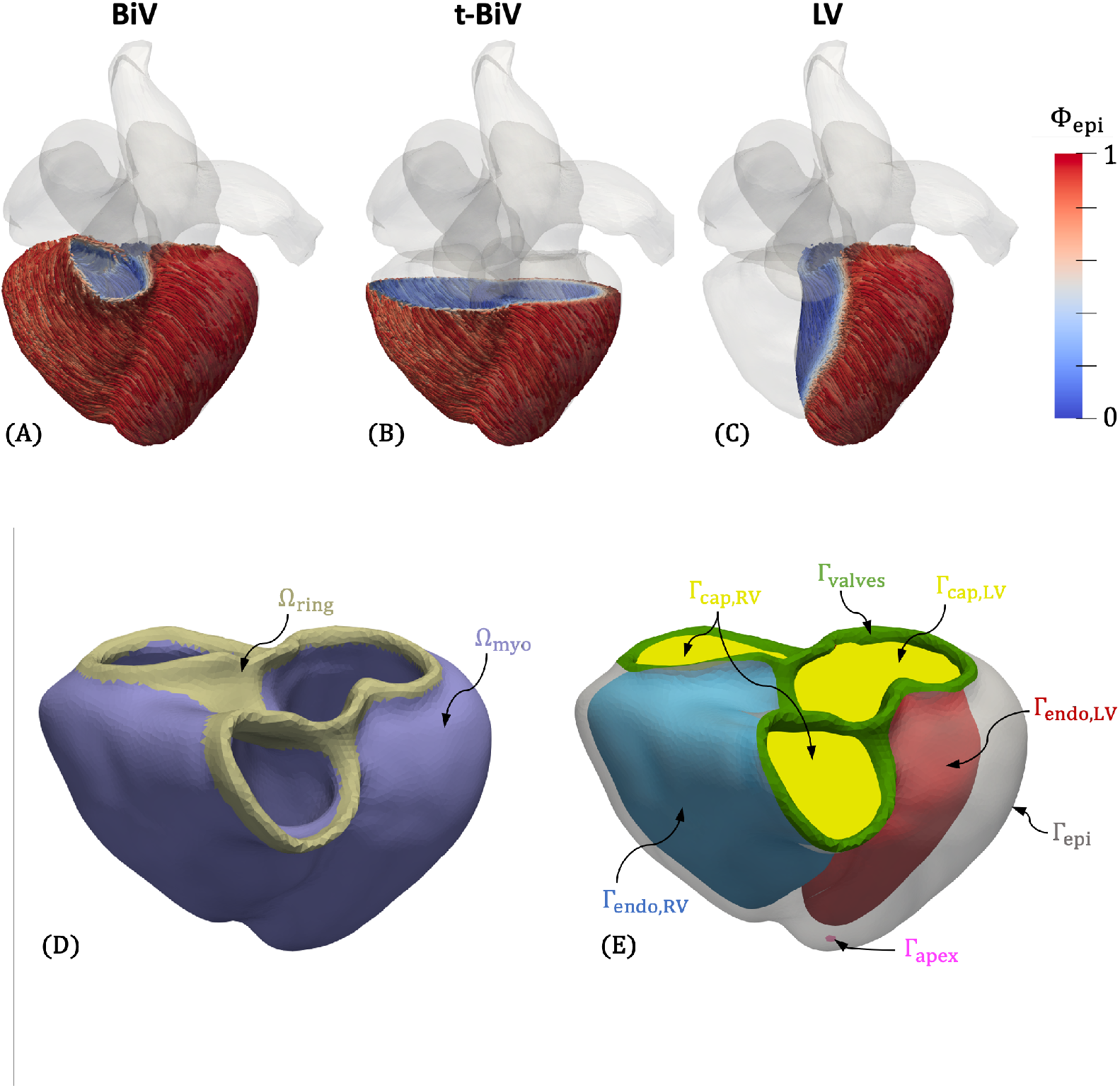
**A-C**: Three anatomical models are considered in this study: biventricle (BiV), biventricle truncated at the basal plane (t-BiV), and left ventricle (LV). The generated longitudinal fiber direction is also shown, colored by the transmural Laplace solution Φ_epi_. **D**: Volume labels for the BiV mesh (shown here for the RR70% configuration). We partition the complete computational domain Ω into a myocardial domain (Ω_myo_) and a valve ring domain (Ω_ring_), where we prescribe different material characteristics. **E**: Surface labels for the BiV mesh. We partition the complete domain boundary Γ = ∂Ω into the epicardium (Γ_epi_), LV endocardium (Γ_endo,LV_), RV endocardium (Γ_endo,RV_), and valve rings (Γ_valve_). We also label the apex (Γ_apex_ ⊂ Γ_epi_) and generate cap surfaces for the LV and RV endocardia (Γ_cap,LV_ and Γ_cap,RV_).

We first construct the BiV model. While MeshDeformNet directly outputs an LV myocardial segmentation, due to the lack of suitable training data, MeshDeformNet does not provide an RV myocardial segmentation (Figure 2). Thus, we create an RV myocardium by dilating the RV lumen segmentation, assuming a uniform RV thickness of 3.5mm [19], which was found to adequately match the image data. Then, we perform smoothing and manual editing in 3D Slicer (https://www.slicer.org/) [100] to improve agreement with the image data. Next, we export the segmentation as a triangulated surface and perform additional smoothing and remeshing in MeshMixer (https://www.meshmixer.com/). We also manually label surfaces (Figure 3E), including the epicardium (Γ_epi_), LV (Γ_endo,LV_) and RV (Γ_endo,RV_) endocardial surfaces, valve rings (Γ_valves_), and epicardial apex (Γ_apex_). The labeled BiV model is then imported into SimVascular (https://simvascular.github.io/) [101], where we create a tetrahedral volume mesh using TetGen [102]. We choose a global maximum edge size of 2 mm, which is comparable to meshes used in other studies [46, 48, 103].

Myofiber directions are generated using the Laplace-Dirichlet rule-based method (LDRBM) of Bayer et al. [93]. Following [94], we consider the valve ring surface (Γ_valves_ in Figure 3E) as the ventricular base. Using the solution fields of several Laplace-Dirichlet problems solved on the domain, the method defines a local orthonormal coordinate system {**f, n, s**} everywhere in the mesh, where **f** is the orientation of fibers, **n** is the normal vector of myocardial sheets, and **s** is a vector orthogonal to both ^1^. The input parameters to the LDRBM are rotation angles on the endocardial and epicardial surfaces for **f** (*α*_endo_, *α*_epi_) and **s** (*β*_endo_, *β*_epi_). In the absence of clinically-measured orientations, we adopt standard literature values for these angles in humans [94] ^2^

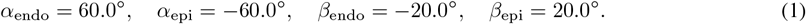

Additionally, the Laplace solution fields are themselves useful parameterizations of the domain, which can inform boundary conditions or enable efficient message passing in graph neural networks [36]. In particular, Φ_epi_, which is 0 on the LV and RV endocardium and 1 on the epicardium, parameterizes the transmural distance, and Φ_ab_, which is 0 at the apex and −1 at the base (or valve rings), parameterizes the apex-to-base or longitudinal direction. Using the same approach, we generate an auxiliary field Φ_l2r_ to distinguish between the LV and RV, assigning a value of 1 on the LV endocardium and 1 on the RV endocardium. Φ_l2r_ is used to help assign a spatially varying epicardial boundary condition (Section 2.3.4), and all three fields are used to partition the BiV model when analyzing simulation results (Section 3.2).

Finally, we perform additional mesh processing necessary for subsequent steps. Using the `vtkFillHolesFilter` from the Visualization Toolkit (VTK) [106], we generate surface meshes that cap the LV and RV endocardial surfaces (Figure 3E), creating water-tight geometries required to compute cavity volumes and fluxes, which are exchanged with the 0D solver in our multiscale coupling scheme [85]. We also label the most basal part of the volume mesh near the valve rings as a separate domain (Ω_ring_ in Figure 3D). This is done by thresholding on Φ_ab_ = 0.995. In this domain, corresponding to the collagen-dense valve annuli [107], we assign a stiff, isotropic material [46, 108], which we found is helpful in preventing excessive deformation of the valve annuli. Finally, to aid prescribing a spatially varying epicardial boundary condition (Section 2.3.4), we compute the long axis of the heart as the vector between the apex Γ_apex_ and the center of the best-fitting plane to the valvular surface Γ_valves_, **v**_long_ = **x**_base_ − **x**_apex_. A normalized long-axis coordinate, Ψ_long_ ^3^, is then generated for each mesh node **x** by projecting its position vector relative to the apex onto the long-axis as,

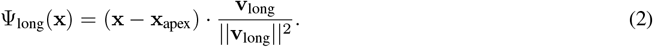

Some points on extreme basal portions of the model, such as pulmonary outlet of the RV, may have Ψ_long_ *>* 1. Thus, we clip these values to be between 0 and 1, as

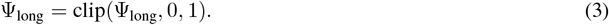

Following model construction from the RR70% images, we propagate the geometry to all other imaged time points (RR0%–RR90%) using the deformations predicted by MeshDeformNet (Figure 2). We denote these image-morphed BiV surface meshes, shown in Figure 4, as Γ_img,RR*i*_, *i* ∈ [0, 10, …, 90]. These are used as target data in an iFEA framework (Section 2.4.2), and as a reference against which we compare the model deformation predicted by our simulations (Section 3).

**Figure 4.**
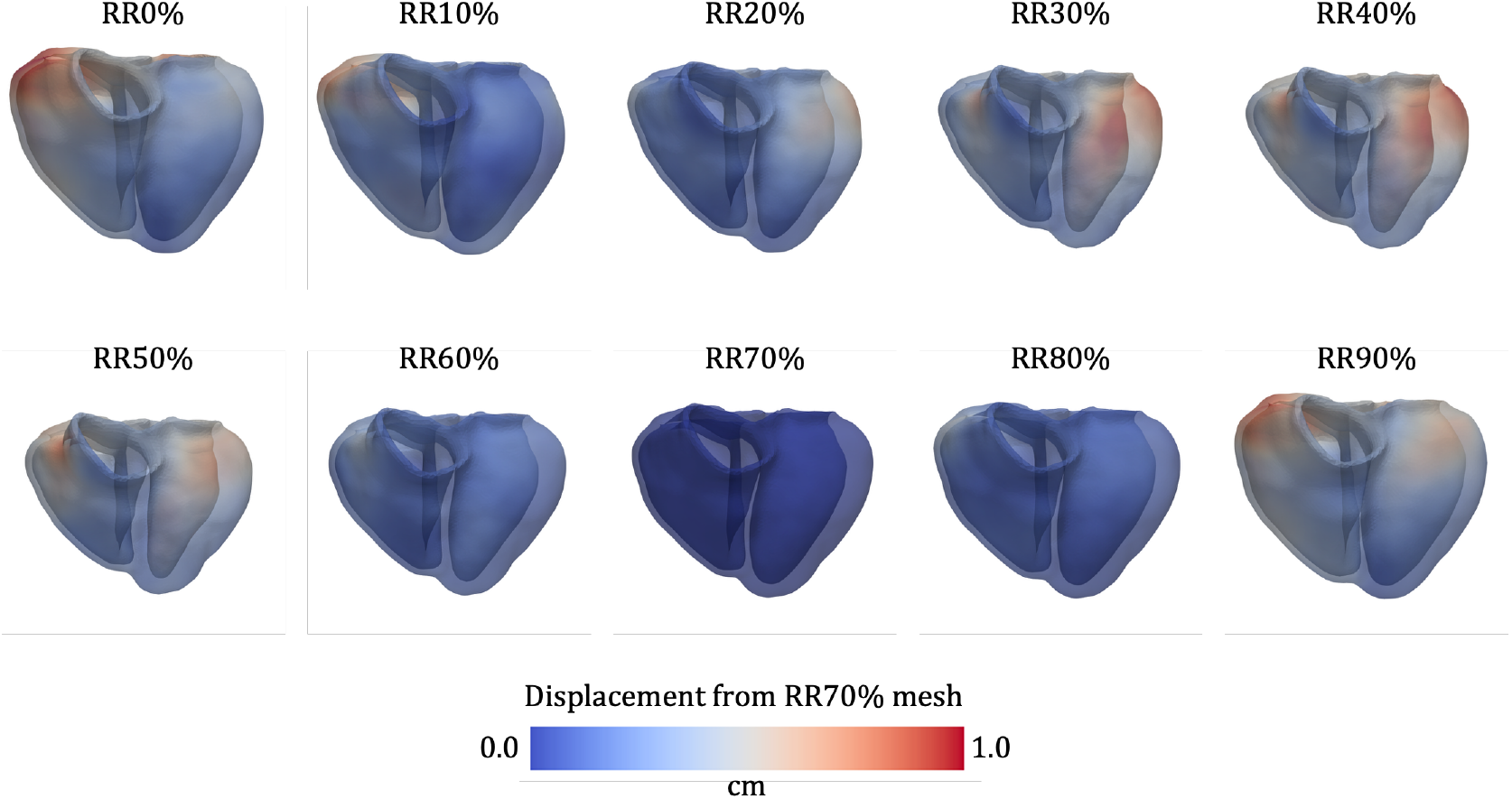
The image-derived motion of BiV surface mesh over the cardiac cycle, denoted Γ_img,RR*i*_, *i* ∈ [0, 10, …, 90]. The deformation is obtained by morphing these models from the RR70% configuration using the motion predicted by MeshDeformNet. The color indicates the magnitude of displacement from the RR70% mesh.

Once the BiV model is fully constructed, the t-BiV and LV models are obtained by manually removing the appropriate portions of the BiV model in MeshMixer. For t-BiV, the truncation is performed using a plane cut approximately perpendicular to the long axis of the heart and just below the protrusion of the pulmonary trunk of the RV (Figure 3B). For the LV-only model, we remove the RV using a combination of plane cuts, erase operations, bridge operations, and smoothing (Figure 3C). BiV surface labels are retained where applicable, and new surface labels are created for the basal plane surface on t-BiV and the septal portion of the epicardium on LV. The simulation setup for the t-BiV and LV models is discussed further in Section 2.5. Both models are then imported to SimVascular and volume meshed using the same method as BiV.

Myofiber directions are generated differently for t-BiV and LV. Instead of solving Laplace-Dirichlet problems on the new geometries, we interpolate the Laplace solution fields from BiV onto t-BiV and LV, then generate myofiber directions using identical parameters. This ensures that at the same anatomical location across the three models, the myocardial mesostructure is identical, facilitating a fair comparison among these models. More details are provided in Section 2.5.

Finally, the additional mesh processing applied to BiV is repeated for t-BiV and LV, with the following differences. For t-BiV, we do not create a valve ring volumetric domain since this region is removed by truncation. Likewise, an RV cap surface is not required for the LV-only model. For both t-BiV and LV, instead of calculating long-axis coordinate (Ψ_long_) on the new geometries, we interpolate Ψ_long_ from BiV onto t-BiV and LV. Again, more details are provided in Section 2.5.

### 2.3 Multiscale model of ventricular mechanics

In this section, we describe our mathematical model of multiscale biventricular mechanics, including the nonlinear solid mechanics formulation, boundary conditions, constitutive model, coupling with 0D circulation, and numerical implementation.

#### 2.3.1 Continuum mechanics formulation

Let Ω denote the computational biventricular domain (Figure 3D). We denote the reference or stress-free configuration as Ω_*X*_ and the current or deformed configuration as Ω_*x*_. The boundary of the domain, Γ = ∂Ω, is partitioned into the epicardium Γ_epi_, LV endocardium Γ_endo,LV_, RV endocardium Γ_endo,RV_, and valve rings Γ_valves_ (Figure 3E). The motion of the myocardium is then given by a time-dependent deformation map, φ : Ω_*X*_ → Ω_*x*_, which maps material points at position **X** in the reference configuration to position **x** in the current configuration, **x** = φ(**X**, *t*). From this deformation, we define the displacement **u** and velocity **v** as

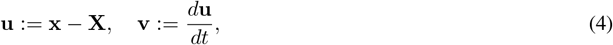

where 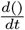 is the total derivative with respect to time. We also define the deformation gradient tensor **F**, along with the Jacobian *J*, right Cauchy-Green tensor **C**, and Green-Lagrange strain tensor **E**,

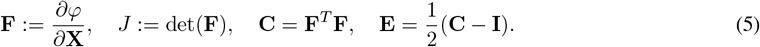

Next, we define the isochoric terms

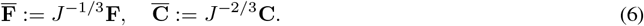

We use a mixed formulation in which the hyperelastic material behavior is described by a Gibbs free energy [110, 111], which is decoupled into isochoric and volumetric components [92],

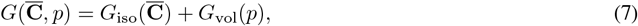

where p is the thermodynamic pressure. *G*_iso_ describes the isochoric material behavior, while *G*_vol_(*p*) is the contribution to the strain energy due to volumetric deformations only.

The governing equations of motion, without body forces, may be written in the current configuration as

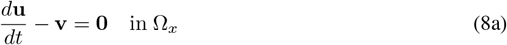

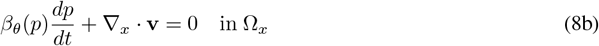

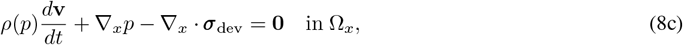

where Eq. (8a) enforces kinematic compatibility between displacement and velocity, Eq. (8b) represents mass continuity and Eq. (8c) represents linear momentum balance. *ρ* is the pressure-dependent material density in the current configuration, *β*_*θ*_ is isothermal compressibility coefficient, and ***σ***_dev_ is the deviatoric part of the Cauchy stress, ***σ*** = ***σ***_dev_ − *p***I**. These are related to the specific Gibbs free energy components (per unit mass) as follows:

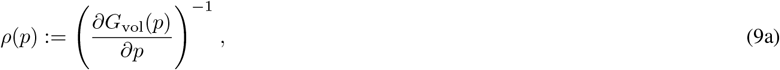

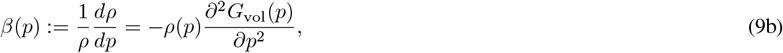

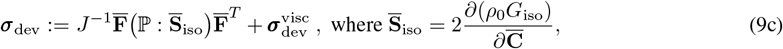

where *ρ*_0_ is the density of the material in the reference configuration and 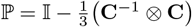 is a projection tensor. Note that in Eq. (9c), in addition to a hyperelastic contribution, we also include a viscous deviatoric stress 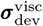, described in Section 2.3.2.

Physiological boundary conditions are applied to the myocardium. For the BiV case, we have

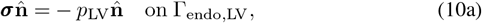

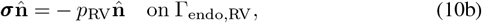

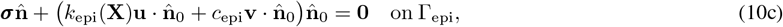

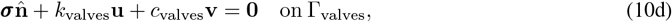

where 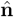 and 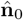 are the unit surface normals in the current and reference configurations, respectively. In Eqs. (10a) and (10b), the blood pressures *p*_*LV*_ and *p*_*RV*_ are obtained via coupling with a 0D circulation model (Section (2.3.3)). Eqs. (10c) and (10d) are Robin boundary conditions [27] applied to the epicardial surface (in the normal direction only) and the valve plane surface (in all directions), with stiffnesses *k*_(·)_ and damping coefficients *c*_(·)_ ^4^. The epicardial Robin BC stiffness *k*_epi_(**X**) is spatially varying and is described further in Section 2.3.4. The problem is initialized in the reference configuration (Section 2.4.2) with zero displacement and velocity.

#### 2.3.2 Constitutive model

Following Eq. (9), the passive hyperelastic response of the material is modeled using an isochoric potential *G*_iso_ and a volumetric potential *G*_vol_. In addition, the contraction of the myocardium is performed using an active stress formulation, so that 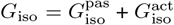, where 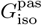 and 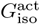represent the passive and active contributions to the strain energy, respectively. Further, the macroscopic viscous behavior in the tissue is modeled using a Newtonian fluid-like viscosity model. Here, we provide the details on these material models.

For the myocardium in Ω_myo_ (Figure 3D), the passive material behavior is modeled with the orthotropic Holzapfel-Ogden (HO) model in its decoupled form [104],

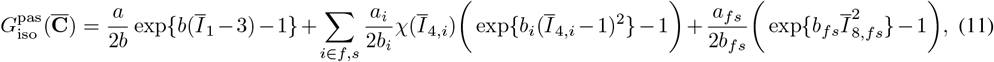

where {*a, a*_*f*_, *a*_*s*_, *a*_*fs*_} are stiffness-like parameters for the isotropic ground matrix, fibers, sheets, and fiber-sheet interactions, respectively, and {*b, b*_*f*_, *b*_*s*_, *b*_*fs*_} control the rate of strain stiffening for the corresponding components. The strain invariants are defined as 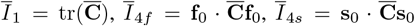, and 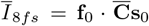, where **f**_0_ and **s**_0_ are the fiber and sheet vectors in the reference configuration, respectively. Finally, *χ*(*η*) is a smoothed Heaviside function centered at *η* = 1, which enforces that fiber and sheet components only provide support in tension, 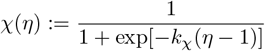[103].

For the volumetric potential, we use the model proposed by Simo and Taylor (ST91) [112], written in terms of the volumetric Gibbs free energy [110]

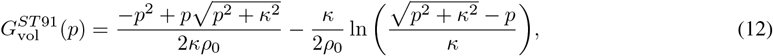

where *κ* is the bulk modulus.

The active stress model free energy may be written,

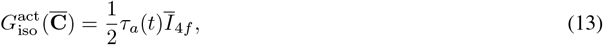

which yields an active stress

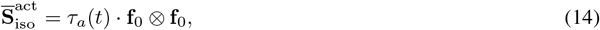

where *τ*_*a*_(*t*) is the prescribed, time-dependent active stress magnitude. We model this active stress with a maximum value *τ*_max_ multiplied by an activation function *a*(*t*),

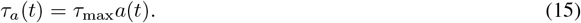

In this work, we use a double Hill activation function [113], defined as

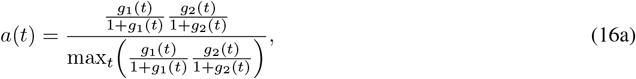

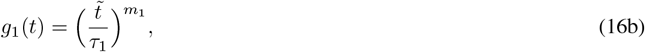

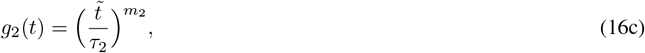

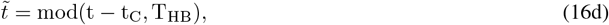

where *τ*_1_, *τ*_2_, *m*_1_, and *m*_2_ control the shape of the curve, *t*_*C*_ is the time at which contraction begins, and *T*_*HB*_ is the cardiac cycle period. A sample active stress curve is shown in Figure 5C. Contraction times are calculated from the collected ECG data, while literature values are used for *τ*_1_, *τ*_2_, *m*_1_, and *m*_2_. More details are provided in Appendix D.

**Figure 5.**
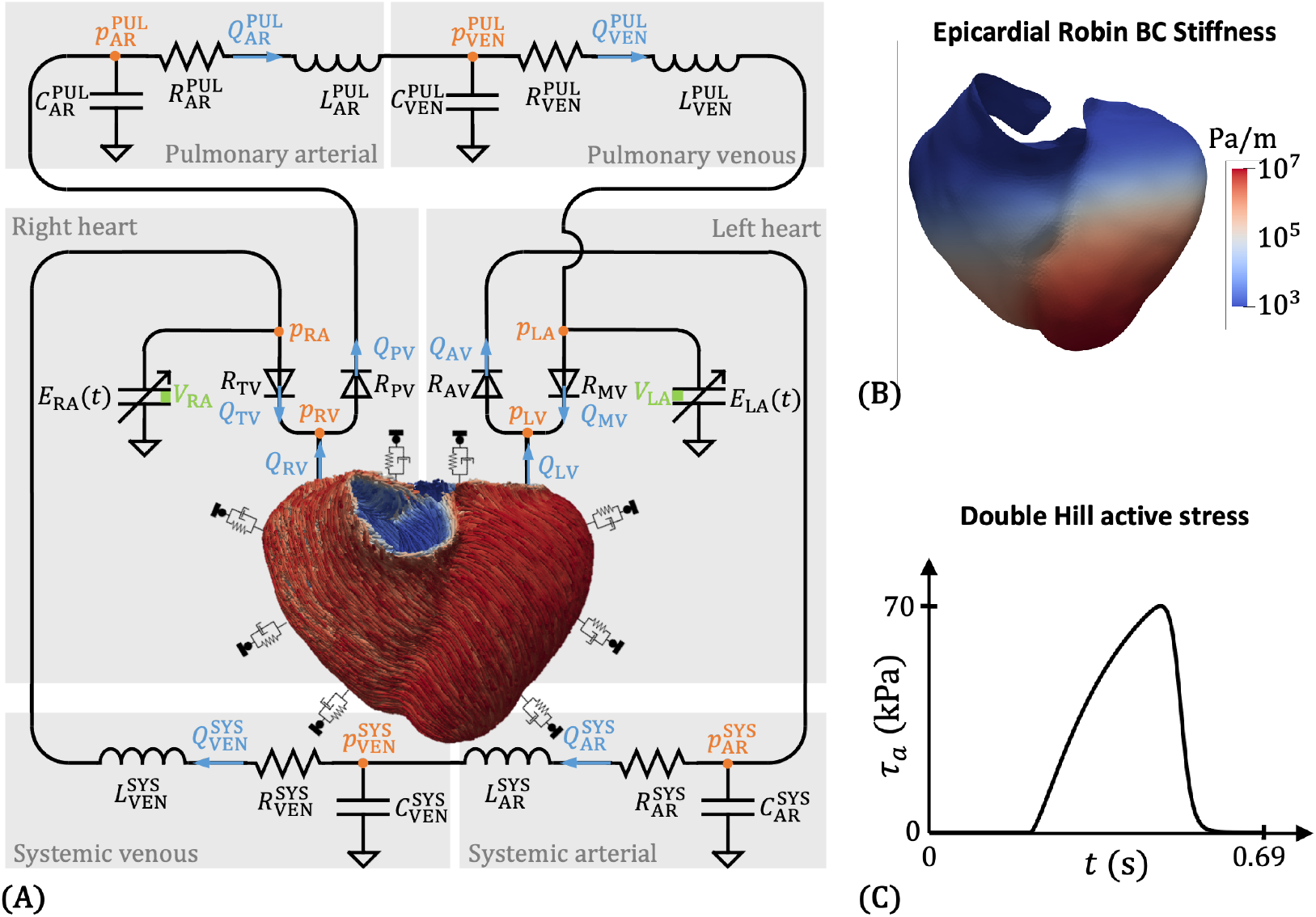
Multiscale BiV model setup. **A**: The BiV model is coupled to a closed-loop 0D circulation model (Section 2.3.3). **B**: A spatially varying Robin boundary condition (BC) is applied on the epicardium to model the mechanical support from the pericardium and surrounding thoracic anatomy (Section 2.3.4). **C**: An active stress formulation with a double Hill activation function is used to model myocardial contraction (Eq. (15)). Activation timing parameters are obtained directly from the collected ECG data (Appendix D).

Viscosity in the myocardium is modeled with a Newtonian viscous stress [92],

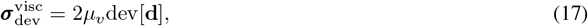

where *µ*_*v*_ is the dynamic viscosity, dev[**d**] is the deviatoric part of the rate of deformation tensor, where

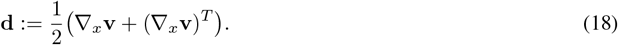

The stiff valvular tissue in Ω_ring_ (Figure 3D) is modeled as described above, with the following differences. To make it isotropic [108], the same HO model (Eq. (11)) is used, but all *a*_(_·_)_ parameters are set to zero except for the isotropic ground matrix term, which we denote *a*_valves_. In addition, we denote the isotropic exponential term as *b*_valves_. Both are fixed to values that were found to be effective in preventing excessive dilation of the valve annuli. To make it passive, no active stress is applied in Ω_ring_.

#### 2.3.3 3D-0D coupling

To model the effects of preload and afterload on the ventricles, we couple our 3D mechanics model to a 0D lumped-parameter network (LPN) model of the systemic and pulmonary circulations [87, 92]. The closed-loop LPN, shown in Figure 5A, is composed of capacitor-resistance-inductor (C-R-L) assemblies for four compartments – systemic arterial, systemic venous, pulmonary arterial, and pulmonary venous. In addition, the atria are modeled as time-varying elastances (TVEs) with double Hill activation functions (Eq. (16) and Appendix D), while the heart valves are represented by non-ideal resistive diodes. Appendix A provides the governing system of ordinary differential equations (ODEs) for a full LPN model (using TVEs for all cardiac chambers, including the LV and RV), and Appendix B explains how the equations are modified when coupling the LPN to a 3D finite element model of the LV and RV.

We couple the 3D BiV mechanical model with the 0D LPN model using our recently developed modular and fully implicit approach [85], which is inspired by the Approximate Newton Method of Chan [114]. The coupling scheme involves the bidirectional exchange of flow rates and pressures: flow rates are passed from the 3D model to the 0D model, while pressures are communicated from the 0D model back to the 3D domain (Figure 5A). Specifically, flow rates *Q*_*LV*_ and *Q*_*RV*_ are calculated as the rates of change of volume enclosed by the LV and RV endocardial surfaces (Γ_endo,LV_ and Γ_endo,LV_), respectively. The cap surfaces Γ_cap,LV_ and Γ_cap,RV_ are used here to close the endocardial surfaces and are essential for accurately computing *Q*_*LV*_ and *Q*_*RV*_. The computed flow rates are supplied as inputs to the 0D solver, which integrates the ODE system of the LPN model. The resulting pressures at the corresponding nodes (*p*_*LV*_ and *p*_*RV*_) are passed to the 3D solver, where they are then applied as follower pressure loads on the LV and RV endocardial surfaces. This exchange occurs at every Newton iteration of the 3D solver within each time step until convergence. To improve the convergence robustness, we include a coupling-related term in the 3D tangent matrix based on the sensitivity of pressure to flow rate (e.g., 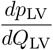). This sensitivity, essentially the boundary resistance seen by the 3D domain due to the 0D domain, is estimated using finite differences [85, 115].

#### 2.3.4 Spatially varying epicardial BC

To account for the mechanical support provided by the pericardium and the surrounding thoracic anatomy [27], including the diaphragm and the lungs [91], we apply a spatially varying Robin BC stiffness on the epicardial surface, similar to the approach employed in Strocchi et al. [42, 116]. We assume that the stiffness decreases from apex to base, where the presence of epicardial adipose tissue is believed to reduce the constraining effect of the pericardium [46]. We first define an apicobasal scaling field *s*_ab_ based on a sigmoidal transformation of the long-axis coordinate Ψ_long_ (Eqs. (2) and (3)),

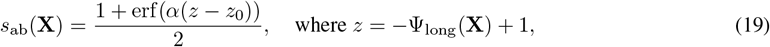

where *z*_0_ determines the location of *s*_ab_ = 0.5, and *α* controls the steepness of the sigmoid. We also define a left-to-right scaling field based on the left-to-right Laplace field Φ_l2r_

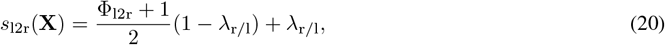

where *λ*_r/l_ is a ratio between the RV and LV maximum epicardial stiffnesses. We found that introducing left-to-right variation was necessary for the model to reproduce the RV free wall motion shown in Figure 4.

The epicardial stiffness is then given as

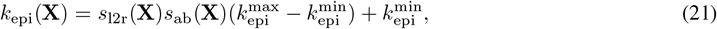

where 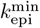 and 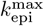are the minimum and maximum stiffnesses over the epicardial surface. Ψ_long_, Φ_l2r_, *s*_ab_, *s*_l2r_, and *k*_epi_ are shown in Figure 6. Parameter values for this BC can be found in Table 6.

**Figure 6.**
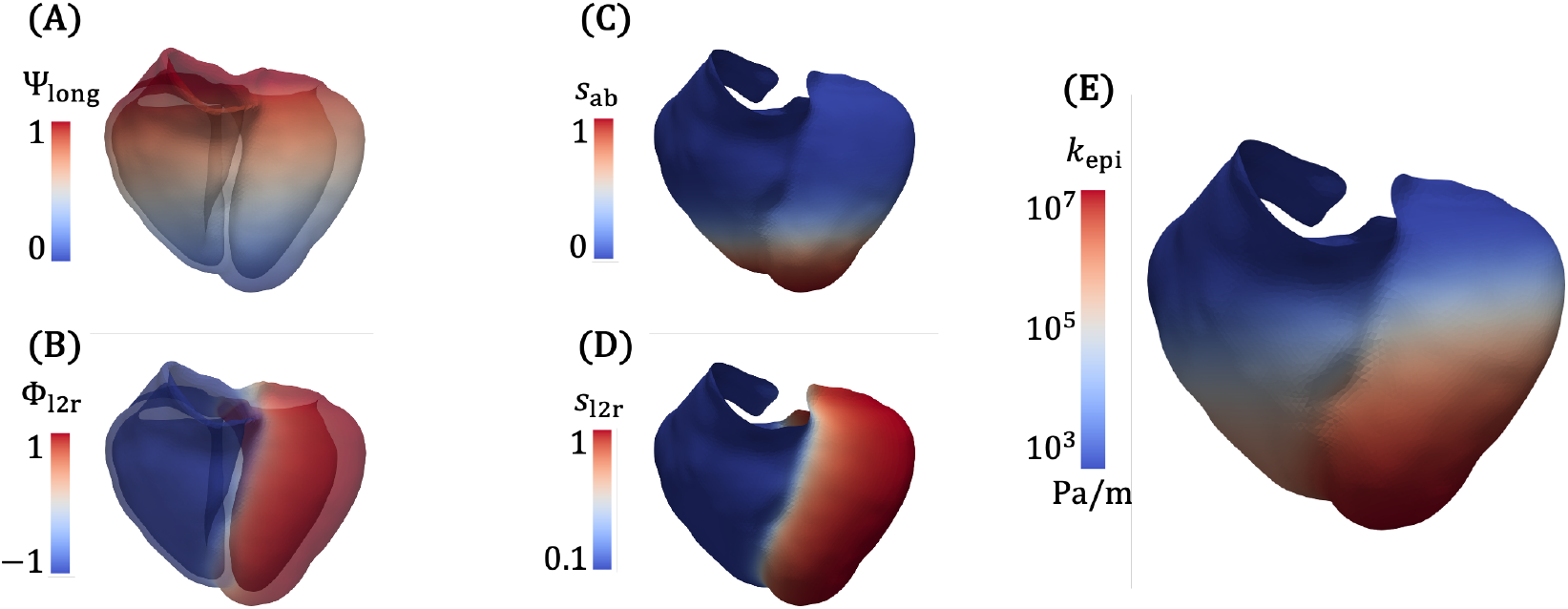
Visualization of various field quantities used to define the spatially varying stiffness for the epicardial Robin BC. **A**: Long-axis coordinate Ψ_long_. **B**: Left-to-right field Φ_l2r_. **C**: Apicobasal scaling field *s*_ab_, computed from Ψ_long_ with *α* = 5 and *z*_0_ = 0.7. **D**: Left-to-right scaling field *s*_l2r_, computed from Φ_l2r_ with *λ*_r/l_ = 0.1. **E**: spatially varying epicardial stiffness *k*_epi_, with 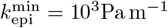 and 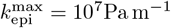.

#### 2.3.5 Numerical implementation

All simulations are performed using an in-house multiphysics finite element solver adapted from SimVascular’s `svFSI` solver [117], which has been applied for a variety of cardiovascular applications [85, 118, 119, 120, 121, 122, 123, 124, 125, 126], and verified for cardiac mechanics modeling [103, 111]. For all simulations, we use four-noded linear tetrahedral elements (TET4), which are chosen for their versatility in meshing complex shapes, such as our patient-specific ventricular models. Our mechanics formulation employs a variational multiscale (VMS) stabilization, which allows equal-order interpolation for velocity and pressure (P1-P1) and mitigates potential volumetric locking [110, 111]. The semi-discrete equations are integrated in time using the implicit generalized-*α* method with a predictor and multi-corrector scheme [110], using a spectral radius of infinite time step (*ρ*_∞_) of 0.5. The resulting nonlinear algebraic system is solved using Newton’s method, and at each Newton iteration, the linear system is solved using the generalized minimal residual (GMRES) method [127]. Linear and nonlinear solver tolerances are both set to 10^−4^. In 3D-0D coupled simulations, a resistance-based preconditioner [85, 128] is used to improve linear solver convergence. In pure 3D passive mechanics simulations, performed during iFEA (Section 2.4.2), we use the thresholded incomplete LU (ILUT) preconditioner available in the Trilinos library (https://trilinos.github.io/).

### 2.4 Multistep personalization procedure

The multiscale mechanics model described thus far depends on dozens of parameters. In this section, we describe our strategy for tuning these parameters so that the model outputs match clinical data. We adopt a multistep personalization strategy, similar to the one developed in Shi et al. [92] for the left atrium, adapted here for BiV mechanics. This approach consists of the following steps, visualized in Figure 7.

**Figure 7.**
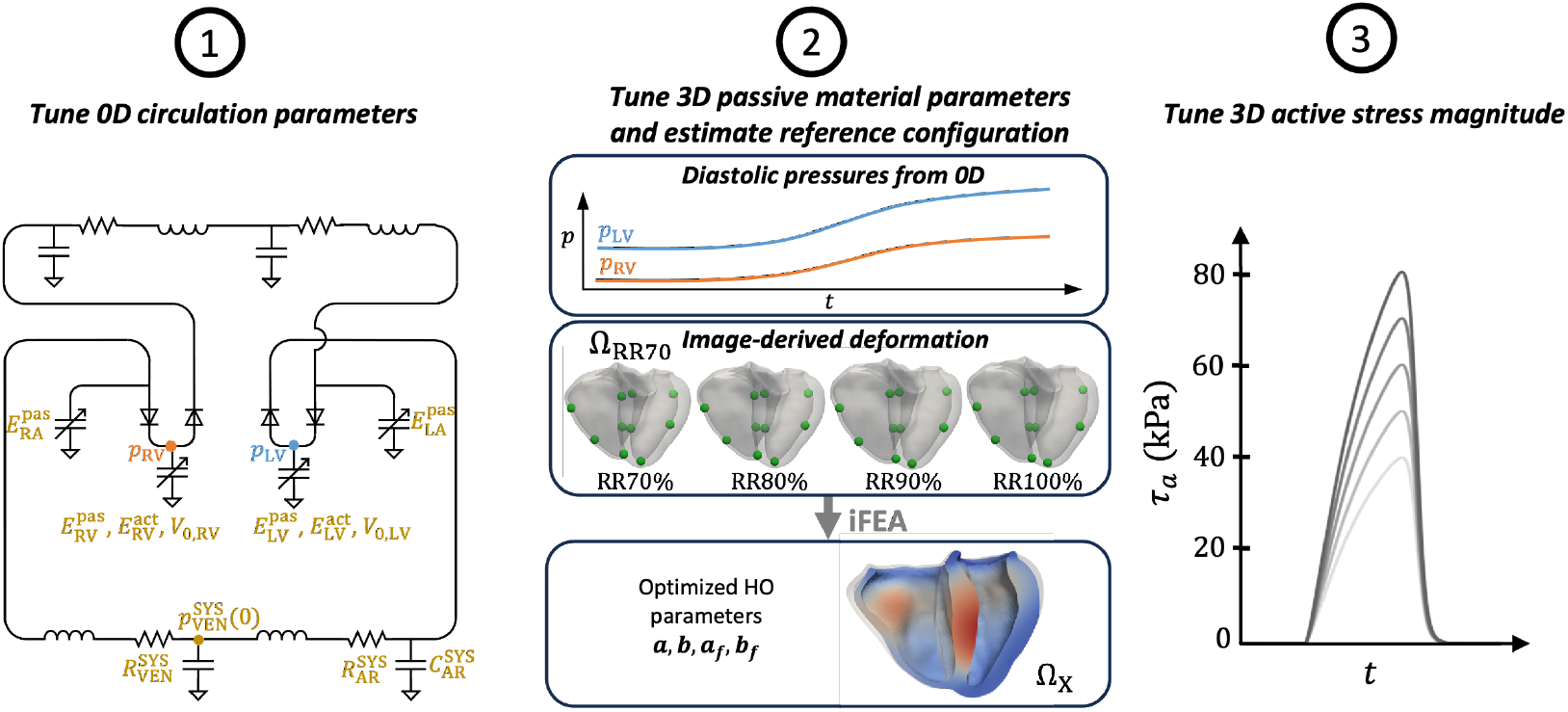
The multiscale BiV model is tuned using a multistep personalization procedure. **Step 1**: We use a full 0D surrogate of the multiscale model (Figure 5A) to efficiently tune key circulation parameters (shown in gold). **Step 2**: Using the LV and RV diastolic pressure from the tuned full 0D surrogate, as well as image-derived diastolic mesh motion with landmark points (green), an iFEA algorithm is used to estimate the HO constitutive model parameters *a, b, a*_*f*_ and *b*_*f*_, as well as the reference configuration Ω_*X*_ [111]. **Step 3**: To personalize the active myocardial behavior, we evaluate the multiscale BiV model over a range of active stress magnitudes *τ*_max_ and choose the value that yields the best fit to the clinical data. iFEA: inverse finite element analysis; HO: Holzapfel-Ogden constitutive model

1. Estimate key 0D circulation parameters using an evolutionary algorithm with a full 0D surrogate.
2. Estimate key HO model parameters Eq. (11) and the reference configuration Ω_*X*_ using an iterative iFEA algorithm [111].
3. Estimate the active stress magnitude Eq. (15) using a parameter sweep. These three steps are elaborated in the following sections.

#### 2.4.1 0D parameter estimation

To tune the 0D circulation parameters, we adopt a full 0D surrogate of the 3D-0D multiscale model, in which the 3D FE model of the ventricles is replaced with TVEs for the LV and RV (Figure 7 Step 1). The governing system of equations is provided in Section A. This full 0D surrogate can be evaluated orders of magnitude faster than the 3D-0D model, allowing us to efficiently estimate the circulation model parameters to fit clinical data. We tune the parameters of our full 0D circulation model, ***θ***, using the following procedure.

The objective function for optimization is based on “target” values of pressure and volume described in Section 2.1. Summarizing here, our targets are

- Patient-specific time-series, minimum, and maximum cardiac volumes 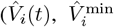 and 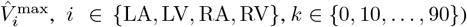(Table 1).
- Patient-specific systolic and diastolic cuff blood pressures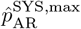 and 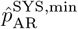(Table 2)
- Reference values for the pulmonary arterial systolic and diastolic blood pressures (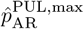 and 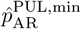 or PAP in Table 3), the peripheral venous pressure (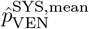 or PVP in Table 3), the central venous pressure (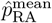or CVP in Table 3), and the pulmonary arterial wedge pressure (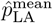 or PAWP in Table 3). These additional literature-based constraints promote physiological circulatory dynamics.

Given measurement uncertainties, we only penalize deviations greater than 5% for the measured clinical data. For the reference pressure values, we penalize deviations outside the ranges provided in Table 3. The objective function is thus defined as,

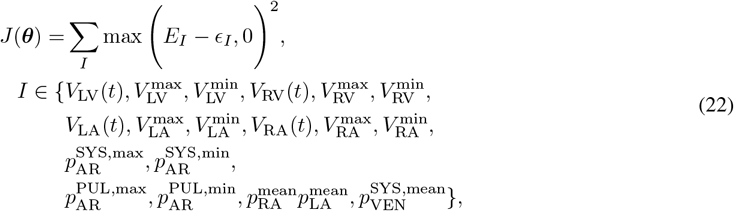

where *E*_*I*_ is the relative error and *ϵ*_*I*_ is the threshold above which we penalize deviations for target *I*. If *I* is a scalar quantity, the error is defined as

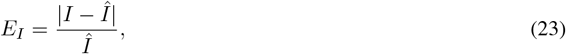

while if *I* is a time-series quantity,

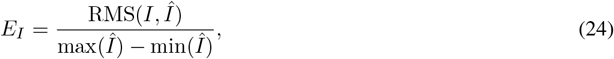

Where

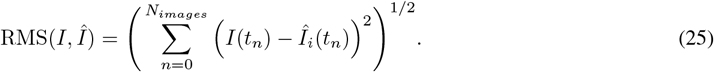

Minimum, maximum, and mean values in time are evaluated over the last two cardiac cycles after reaching a limit cycle, which was obtained by running for 30 cardiac cycles.

Using literature values [30, 129], we obtain preliminary parameter values that result in physiological results, assessed through atrial and ventricular PV loops, as well as systemic and pulmonary pressures and flow waveforms. Based on the sensitivity analysis for this LPN presented in Salvador et al. [30], the most important parameters for the target metrics are

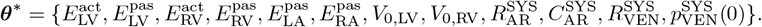

This subset includes the passive 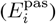 elastances of all cardiac chambers, the active 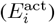 elastances and rest volumes (*V*_0,*i*_) of the LV and RV, the systemic arterial resistance 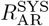 and capacitance 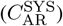, the systemic venous resistance 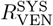, and the initial pressure in the systemic venous compartment 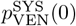. 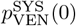 was chosen because, since 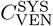 is large (Table 5), it is a major determinant of the total blood volume, which was shown to be a very important parameter [30, 130]. We tune only this subset, leaving all other parameters unchanged, since they have only a secondary effect on the target quantities of interest.

To solve the optimization problem, we use the `differential_evolution` algorithm in the `scipy.optimize` library. The parameters are bounded in the range 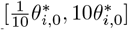, where 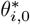 is the preliminary value of the parameter. We use a population size multiplier `popsize` of 20 with Sobol sequence initialization. Additionally, we configure the algorithm using “deferred updating,” which permits parallel function evaluations in each generation.

We conclude this section by noting that, in addition to providing optimized 0D parameters, this process also generates 0D initial conditions that immediately produce a limit cycle for the full 0D model. When applied in the 3D-0D model, these initial conditions help the simulation reach a limit cycle within just a few cardiac cycles [131].

#### 2.4.2 Personalizing passive mechanics

In step 2, we personalize the 3D myocardial passive mechanics, simultaneously estimating passive material parameters and the reference configuration Ω_*X*_ (Figure 7, Step 2). We employ a recently developed iFEA framework involving a nested iteration scheme [111]. In the outer iteration, material parameters are estimated to match LV and RV volumes and landmark point displacements using successive simulations of the passive inflation of the BiV model from diastasis to end-diastole (RR70% to RR100%). Five landmark points each were selected on the LV and RV endocardial surfaces using a semi-automatic method. We use a custom-designed genetic algorithm presented in Shi et al. [111], although other optimization algorithms may be effective. In the inner iteration, a modified Sellier’s algorithm is used to estimate a reference configuration Ω_*X*_ that, when pressurized, matches the (loaded) RR70% configuration Ω_RR70_ with fixed material parameters from the outer iteration.

Since the reference configuration Ω_*X*_ is estimated during the iFEA procedure, it is important to emphasize that quantities originally defined in the imaged RR70% configuration Ω_RR70_ must be consistently mapped to the current estimate of Ω_*X*_ at each iFEA iteration. In particular, the spatially varying epicardial boundary stiffness *k*_epi_ is specified in Ω_RR70_ (Section 2.3.4), but is applied on Ω_*X*_ and remains active in each iteration. This means that the epicardial boundary “springs” are unstressed in Ω_*X*_ rather than in Ω_RR70_. Importantly, no explicit computation is required for this mapping, because *k*_epi_ is a scalar quantity defined nodewise on the Ω_RR70_ mesh, and the node ordering is preserved when generating Ω_*X*_. Similarly, myocardial fiber orientations {**f, n, s**} are defined in Ω_RR70_ (Section 2.2) and must be mapped to the evolving estimate of Ω_*X*_. Unlike the boundary condition mapping, this step requires additional computation, in that the fibers must be transformed as material directions using deformation gradient from Ω_RR70_ to Ω_*X*_ and subsequently renormalized. Once the final estimate of Ω_*X*_ is obtained at the conclusion of the iFEA procedure, both *k*_epi_ and **f, n, s** are again mapped onto Ω_*X*_, in preparation for the subsequent 3D–0D simulations.

The method requires as inputs the diastolic LV and RV pressures during passive inflation, as well as the deformation of the BiV model during the same phase. While the deformation is obtained from the measured CTA data (Section 2.2 and Figure 4), *p*_LV_ and *p*_RV_ were not measured clinically. Instead, we use *p*_LV_ and *p*_RV_ profiles generated by the tuned full 0D model from the previous step (Section 2.4.1). We apply the iFEA framework to estimate four HO constitutive model parameters, *a, b, a*_*f*_, and *b*_*f*_ (Eq. 11), which were shown in Lazarus et al. [132] to be the most important for LV passive mechanics. These parameters are optimized in the bounds *a* ∈ [10^1^, 10^3^], *a*_*f*_ ∈ [10^1^, 10^4^], *b* ∈ [1, 20], and *b*_*f*_ ∈ [1, 20]. The other HO parameters (*a*_*s*_, *b*_*s*_, *a*_*fs*_, *b*_*fs*_) are fixed at literature values [104].

#### 2.4.3 Tuning active stress

Finally, in step 3, we tune the magnitude of active stress *τ*_max_ (Eq. (15)) by evaluating the 3D-0D BiV model over a range of *τ*_max_ and assessing the goodness of fit in terms of pressure and volume (as in Section 2.4.1), as well the local deformation compared to the CTA-derived motion (Figure 4). In preliminary tests, we found 60 kPa a good baseline value for our model, so we perform simulations with the following active stress magnitudes, *τ*_max_ = {40, 50, 60, 70, 80} kPa. All simulations use the optimized 0D parameters identified in Section 2.4.1 and the optimized HO model parameters and estimated reference configuration obtained in Section 2.4.2. Additionally, all simulations are run for five cardiac cycles, which we found to be adequate for reaching a limit cycle [35].

### 2.5 Simulation setup for t-BiV and LV

Thus far, we have described the simulation setup and personalization for the BiV model only. In this section, we describe the simulation setup for the other two geometries – t-BiV and LV. Our philosophy when establishing the t-BiV and LV models is to isolate the effect of *geometry* as much as possible. To this end, we keep all model inputs identical to those for the personalized BiV model. This is straightforward for inputs like the 0D circulation parameters and active/passive material parameters, which were optimized in Section 2.4. However, some inputs, namely myofiber directions, the spatially varying epicardial BC stiffness, and the iFEA-estimated reference configuration, require careful treatment, which we describe here.

As shown in Figure 8, these three inputs are generated for the t-BiV and LV models by interpolating the necessary mesh fields from the BiV model. First, we recall that the t-BiV and LV geometric models were constructed from the BiV geometric model by removing the appropriate portions, and that all models represent the RR70% configuration Ω_RR70_. Myofiber directions t-BiV and LV are generated by interpolating the Laplace-Dirichlet solution fields from BiV, then executing the fiber generation algorithm with identical epicardial and endocardial fiber angles (Eq. (1)). The stiffness for the spatially varying epicardial Robin BC is obtained by interpolating the long-axis coordinate (Ψ_long_, Eqs. (2) and (3)) and the left-to-right coordinate (Φ_l2r_), which is then transformed into a stiffness map using identical parameters as those of the BiV model (Eqs. (19), (20), and (21)). Finally, to obtain the reference configuration Ω_*X*_ for t-BiV and LV, we first compute the displacement field **u** from Ω_RR70_ to the iFEA-estimated reference configuration for the BiV model (Section 2.4.2). Then, we interpolate **u** onto t-BiV and LV and warp the meshes by this displacement to obtain corresponding reference configuration meshes^5^. For all interpolations, we use Gaussian interpolation and set the kernel radius to be the average edge length of the BiV mesh. This procedure guarantees that, at the same anatomical location across the three models, the fiber orientations, epicardial stiffness, and reference configuration of material points are identical.

**Figure 8.**
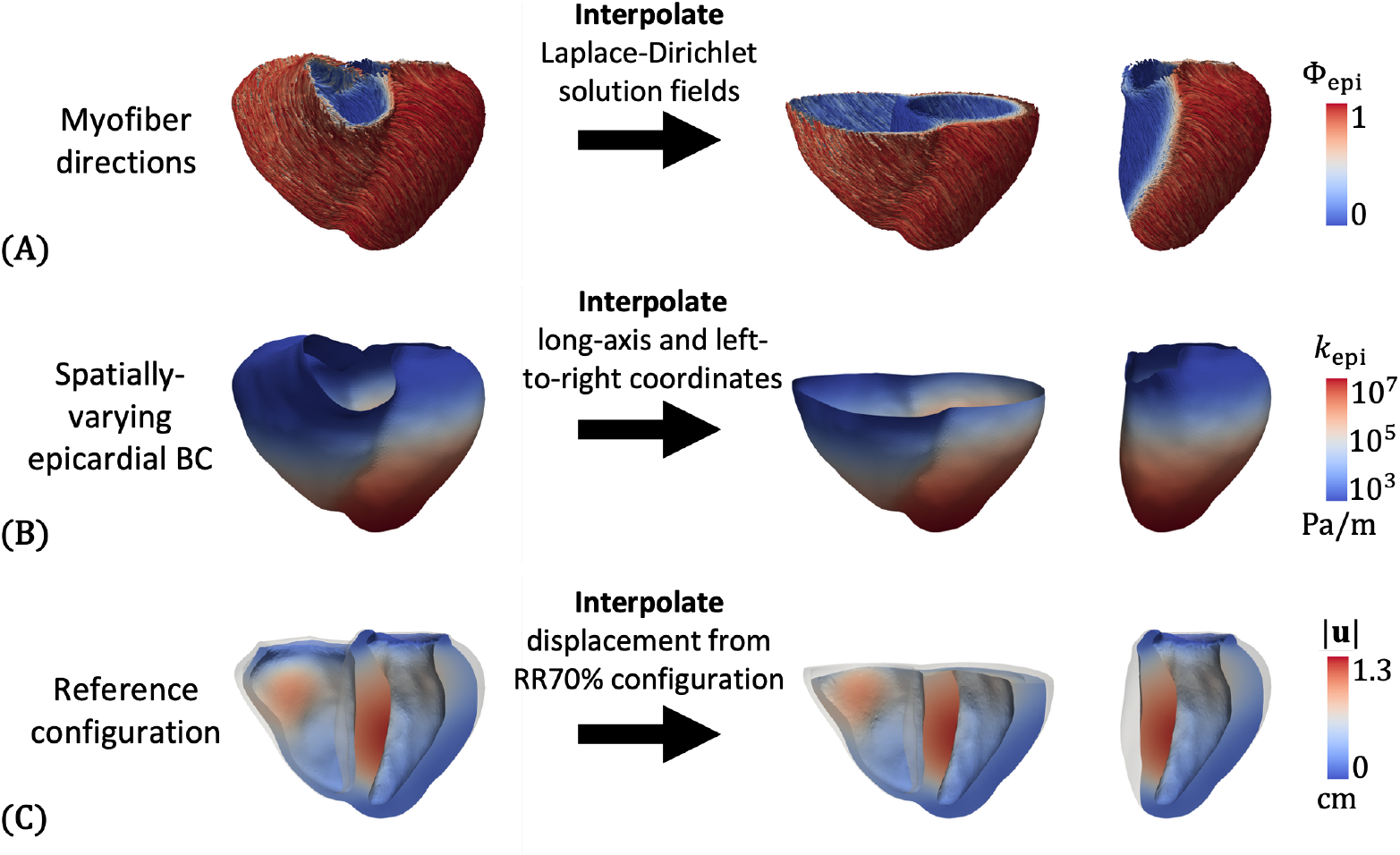
Details on simulation inputs for models t-BiV and LV models. **A**: Myofiber directions are generated by interpolating the Laplace-Dirichlet solution fields, then generating fiber directions using the same parameters as for the BiV model (Section 2.2). **B**: The spatially varying epicardial BC is generated by interpolating the long-axis (Ψ_long_) and left-to-right coordinates (Φ_l2r_), then generating the spatially varying stiffness profile (*k*_epi_) using the same parameters as for the BiV model (Section 2.3.4). **C**: The reference configuration is generated by first computing the nodal displacements **u** from the RR70% configuration to the reference configuration for the BiV model (Section 2.4.2), then interpolating **u** onto t-BiV and LV models, and finally warping these models by this interpolated displacement field.

We also note that for the LV model, since we no longer have a 3D RV, we modify the LPN circulation model to include a TVE model for the RV (Appendix A). We use the RV elastance parameters from the tuned full 0D model from Section 2.4.1 for the coupled 3D-0D simulations of LV mechanics.

The new geometries also introduce new surfaces on which we must define boundary conditions. On t-BiV, truncation creates an artificial basal plane face Γ_base_ (Figure 9A). On LV, part of the outer surface is the septal part of the RV endocardial surface. We label this Γ_epi,septum_, to distinguish it from the free wall portion of the LV epicardium, Γ_epi,free_ (Figure 9B).

**Figure 9.**
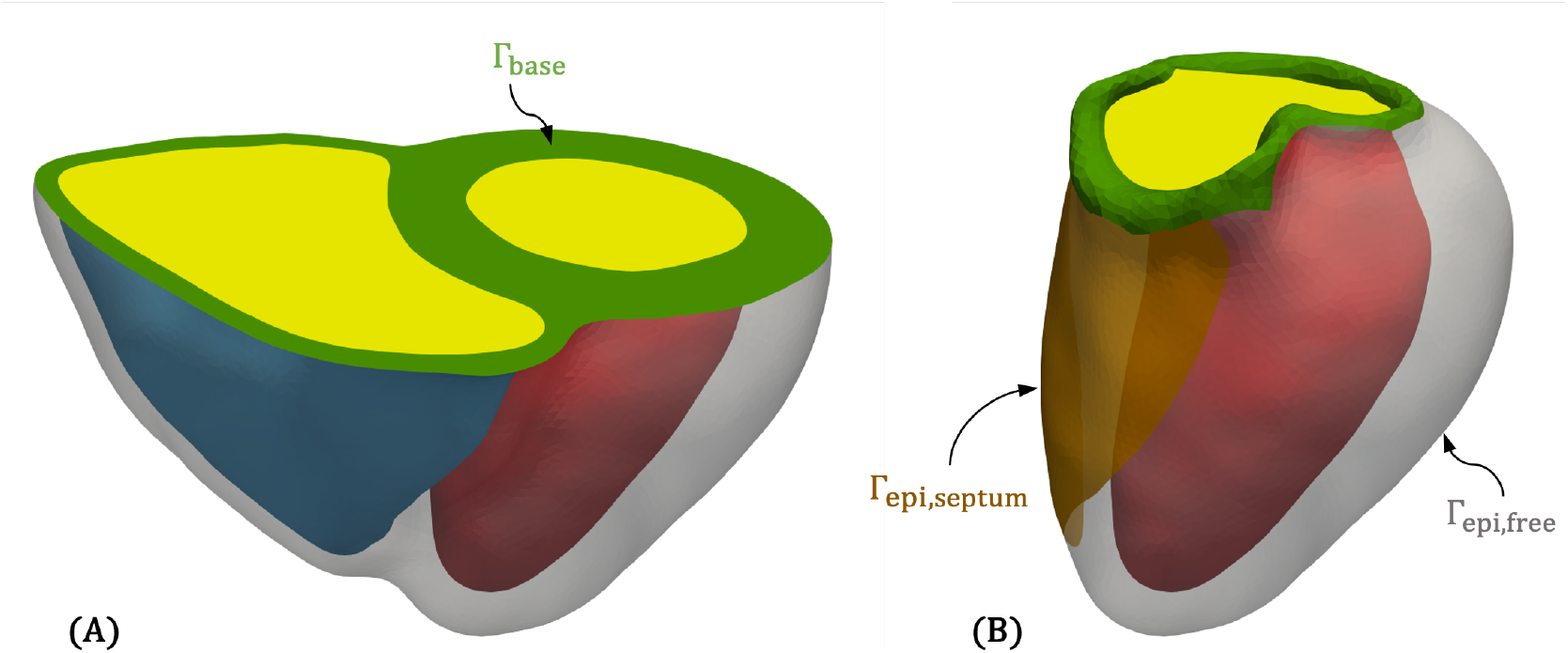
Labeled surfaces on models t-BiV and LV. **A**: On t-BiV, the truncation introduces an artificial basal plane face Γ_base_. **B**: On LV, we divide the epicardial boundary into a septal portion Γ_epi,septum_ and a free wall portion Γ_epi,free_.

For the t-BiV model, we apply a Robin BC (in all directions) on Γ_base_, as was done for Γ_valves_ on the BiV model. We consider three variations (Table 4). For t-BiV(same base), we prescribe the basal Robin BC stiffness, denoted *k*_base,t_ −_BiV_, with the same value as was used for the BiV model (*k*_base,t_ − _BiV_ = *k*_valves_). As we show in Section 3.3, this can lead to unphysiological deformation of the basal surfaces. Thus, we also consider a case with a stiffness ten times greater (*k*_base,t−BiV_ = 10*k*_valves_), denoted t-BiV(stiffer base). Finally, we consider a case with both a stiffer base and active stress with two times greater magnitude (*τ*_max,t−BiV_ = 2*τ*_max_), denoted t-BiV(higher stress).

**Table 4.** Summary of cases simulated in this work. We consider the BiV case as the baseline, and evaluate three variations for truncated BiV (t-BiV) model and three variations for the LV model. The surfaces Γ_base_, Γ_epi,free_, and Γ_epi,septum_ are shown in Figure 9.

| Case | Boundary condition description | Active stress magnitude |
| --- | --- | --- |
| BiV | Robin on $\Gamma_{\text{valves}}$ with stiffness $k_{\text{valves}}$ | $\tau_{\text{max}}$ |
| t-BiV(same base) | Robin on $\Gamma_{\text{base}}$ with stiffness $k_{\text{base,t-BiV}} = k_{\text{valves}}$ | $\tau_{\text{max}}$ |
| t-BiV(stiffer base) | Robin on $\Gamma_{\text{base}}$ with stiffness $k_{\text{base,t-BiV}} = 10k_{\text{valves}}$ | $\tau_{\text{max}}$ |
| t-BiV(higher stress) | Robin on $\Gamma_{\text{base}}$ with stiffness $k_{\text{base,t-BiV}} = 10k_{\text{valves}}$ | $2\tau_{\text{max}}$ |
| LV(all Robin) | Robin on $\Gamma_{\text{epi}} = \Gamma_{\text{epi,septum}} \cup \Gamma_{\text{epi,free}}$ | $\tau_{\text{max}}$ |
| LV(free septum) | Robin on $\Gamma_{\text{epi,free}}$, traction-free on $\Gamma_{\text{epi,septum}}$ | $\tau_{\text{max}}$ |
| LV(loaded septum) | Robin on $\Gamma_{\text{epi,free}}$, RV pressure on $\Gamma_{\text{epi,septum}}$ | $\tau_{\text{max}}$ |

For the LV model, there is some ambiguity about what BC should be applied to Γ_epi,septum_. We test the following three cases (Table 4). For case LV(all Robin), we define a single epicardial surface Γ_epi_ = Γ_epi,septum_ ∪Γ_epi,free_ and apply a spatially varying Robin BC over the entire epicardial surface (as shown in Figure 8). For case LV(free septum), we apply a spatially varying Robin BC on the free wall portion Γ_epi,free_, and apply a traction-free (i.e., zero pressure) BC on Γ_epi,septum_. Finally, in case LV(loaded septum), we again impose a spatially varying Robin BC on Γ_epi,free_, and apply RV pressure on Γ_epi,septum_ [80]. The RV pressure is obtained from the 0D RV element within the LPN that is coupled to the LV model.

## 3 Results

### 3.1 Personalization results

We begin by presenting the results of our multistep personalization procedure. In Section 3.1.1, we detail the outcomes of the 0D parameter estimation using a full 0D surrogate model. Section 3.1.2 follows with personalized passive mechanics results obtained from the iFEA framework. Finally, Section 3.1.3 presents the results of the active stress magnitude parameter sweep. All parameter values used in this study can be found in Tables 5, 6, and *D*.1.

**Table 5.** Parameters of the tuned full 0D circulation model. Values optimized using the method described in Section 2.4.1 are marked by a red dagger *†*.

| Parameter | Symbol | Value | Units |
| --- | --- | --- | --- |
| *General parameters* |  |  |  |
| Time step size | $\Delta t_{0D}$ | $10^{-3}$ | s |
| Number of cardiac cycles | $N_{\text{cycles},0D}$ | 30 | — |
| *Chamber time-varying elastance* |  |  |  |
| LA active elastance | $E_{\text{LA}}^{\text{act}}$ | 0.20 | mmHg mL<sup>-1</sup> |
| LA passive elastance | $E_{\text{LA}}^{\text{pas}}$ | 0.153† | mmHg mL<sup>-1</sup> |
| RA active elastance | $E_{\text{RA}}^{\text{act}}$ | 0.06 | mmHg mL<sup>-1</sup> |

Table 5 – continued from previous page
| Parameter | Symbol | Value | Units |
| --- | --- | --- | --- |
| RA passive elastance | $E_{RA}^{pas}$ | 0.037<sup>†</sup> | mmHg mL<sup>-1</sup> |
| LV active elastance | $E_{LV}^{act}$ | 2.504<sup>†</sup> | mmHg mL<sup>-1</sup> |
| LV passive elastance | $E_{LV}^{pas}$ | 0.095<sup>†</sup> | mmHg mL<sup>-1</sup> |
| RV active elastance | $E_{RV}^{act}$ | 0.493<sup>†</sup> | mmHg mL<sup>-1</sup> |
| RV passive elastance | $E_{RV}^{pas}$ | 0.041<sup>†</sup> | mmHg mL<sup>-1</sup> |
| LA rest volume | $V_{0,LA}$ | 4.0 | mL |
| RA rest volume | $V_{0,RA}$ | 4.0 | mL |
| LV rest volume | $V_{0,LV}$ | 1.976<sup>†</sup> | mL |
| RV rest volume | $V_{0,RV}$ | 54.316<sup>†</sup> | mL |
| **Systemic circulation** |  |  |  |
| Systemic arterial resistance | $R_{AR}^{SYS}$ | 0.677<sup>†</sup> | mmHg s mL<sup>-1</sup> |
| Systemic arterial capacitance | $C_{AR}^{SYS}$ | 0.925<sup>†</sup> | mL mmHg<sup>-1</sup> |
| Systemic arterial inductance | $L_{AR}^{SYS}$ | 0.005 | mmHg s<sup>2</sup> mL<sup>-1</sup> |
| Systemic venous resistance | $R_{VEN}^{SYS}$ | 0.064<sup>†</sup> | mmHg s mL<sup>-1</sup> |
| Systemic venous capacitance | $C_{VEN}^{SYS}$ | 60.0 | mL mmHg<sup>-1</sup> |
| Systemic venous inductance | $L_{VEN}^{SYS}$ | 0.0005 | mmHg s<sup>2</sup> mL<sup>-1</sup> |
| **Pulmonary circulation** |  |  |  |
| Pulmonary arterial resistance | $R_{AR}^{PUL}$ | 0.032 | mmHg s mL<sup>-1</sup> |
| Pulmonary arterial capacitance | $C_{AR}^{PUL}$ | 10.0 | mL mmHg<sup>-1</sup> |
| Pulmonary arterial inductance | $L_{AR}^{PUL}$ | 0.0005 | mmHg s<sup>2</sup> mL<sup>-1</sup> |
| Pulmonary venous resistance | $R_{VEN}^{PUL}$ | 0.035 | mmHg s mL<sup>-1</sup> |
| Pulmonary venous capacitance | $C_{VEN}^{PUL}$ | 16.0 | mL mmHg<sup>-1</sup> |
| Pulmonary venous inductance | $L_{VEN}^{PUL}$ | 0.0005 | mmHg s<sup>2</sup> mL<sup>-1</sup> |
| **Valve resistances** |  |  |  |
| Open valve resistance | $R_{min}$ | 0.005 | mmHg s mL<sup>-1</sup> |
| Closed valve resistance | $R_{max}$ | 50.0 | mmHg s mL<sup>-1</sup> |
| **Initial conditions** |  |  |  |
| LA volume | $V_{LA}(0)$ | 75.61 | mL |
| RA volume | $V_{RA}(0)$ | 75.11 | mL |
| LV volume | $V_{LV}(0)$ | 140.87 | mL |
| RV volume | $V_{RV}(0)$ | 125.17 | mL |
| Systemic arterial pressure | $p_{AR}^{SYS}(0)$ | 99.49 | mmHg |
| Systemic venous pressure | $p_{VEN}^{SYS}(0)$ | 3.703<sup>†</sup> | mmHg |
| Pulmonary arterial pressure | $p_{AR}^{PUL}(0)$ | 26.25 | mmHg |
| Pulmonary venous pressure | $p_{VEN}^{PUL}(0)$ | 24.83 | mmHg |
| Systemic arterial flow | $Q_{AR}^{SYS}(0)$ | 74.80 | mL s<sup>-1</sup> |
| Systemic venous flow | $Q_{VEN}^{SYS}(0)$ | 91.78 | mL s<sup>-1</sup> |
| Pulmonary arterial flow | $Q_{AR}^{PUL}(0)$ | 47.00 | mL s<sup>-1</sup> |
| Pulmonary venous flow | $Q_{VEN}^{PUL}(0)$ | 46.21 | mL s<sup>-1</sup> |

#### 3.1.1 Personalized full 0D model

The tuned circulation model parameters are provided in Table 5, where we use a red dagger *†* to mark the values that are optimized according to the procedure described in Section 2.4.1. The full 0D model outputs with these parameters are shown in Figure 10. PV loops for both the atria and ventricles are shown in Figure 10A-B, where the colors indicate the phase of the cardiac cycle – atrial contraction (green), isovolumic contraction (yellow), ventricular ejection (red), isovolumic relaxation (purple), and ventricular passive filling (blue). The ventricular PV loops display physiological profiles, with stroke volumes of 73.5 mL for LV and 72.6 mL for RV. The atrial PV diagram exhibits an a-loop, associated with atrial contraction, while the v-loop, associated with the atrial reservoir and conduit phases [92], collapses to a straight line with a slope equal to the atrial passive elastance.

**Figure 10.**
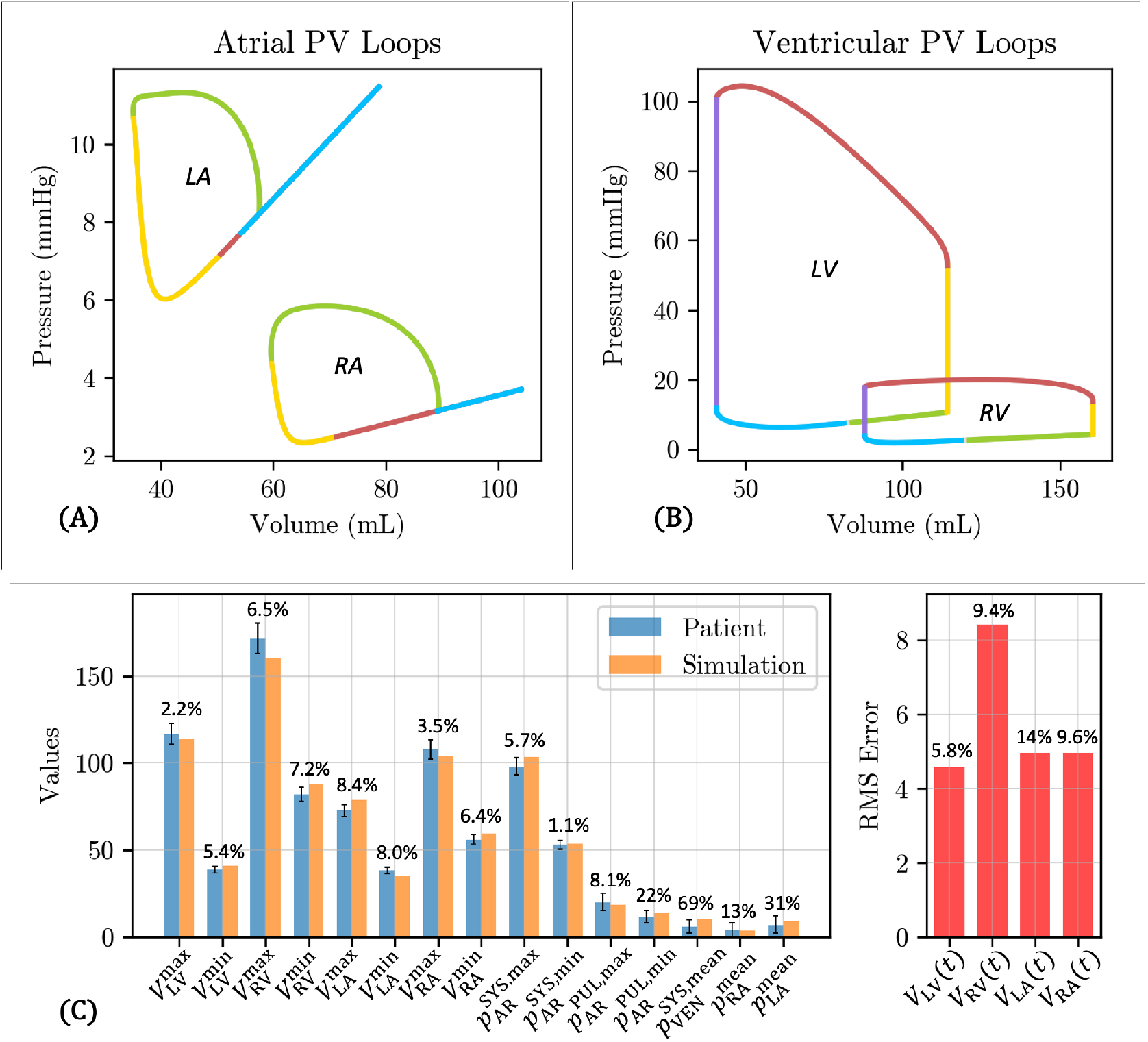
Results of the tuned full 0D circulation model. **A-B**: PV loops for LA, RA, LV, and RV. Colors indicate the phase of the cardiac cycle – atrial contraction (green), isovolumic contraction (yellow), ventricular ejection (red), isovolumic relaxation (purple), and ventricular passive filling (blue). **C**: (Left) Comparison between simulation outputs (orange) and scalar clinical targets (blue) for the studied patient. Error bars indicate the uncertainty in target patient values. (Right) RMS errors for the chamber volumes represent the average volume error across all imaged cardiac phases (RR0%, …, RR90%) and are computed using Eq. (25). *V*_(_·_)_ quantities in mL and *p*_(_·_)_ quantities are in mmHg. Percentage values above each bar indicate the percent relative error (Eqs. (23) and (24)).

The tuned full 0D model outputs show a decent agreement with the target clinical data (Figure 10C). For the measured clinical data (chamber volumes and cuff blood pressure), the highest relative error (Eq. (23)) is 8.4% for the maximum LA volume 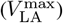. The relative errors are substantial for the reference pressure values 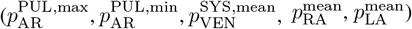, although these errors fall within the uncertainty ranges for these values (Table 3). The normalized RMS errors in the four chamber volumes (Eq. (24)), which measure the discrepancy in chamber volume over the entire cardiac cycle, are all below 10%, except for *V*_LA_(*t*), which has a slightly higher relative error of 14%. However, the absolute value of the RMS error is reasonably low at 5 mL measured over the entire cardiac cycle. Overall, the tuned full 0D circulation model produces physiological PV loops and reasonably matches the measured clinical data.

In terms of computational performance, 0D parameter estimation using the full 0D surrogate takes approximately 4 hours using 20 CPUs of an Intel Gold 5118 2.3 GHz processor. The evolutionary algorithm converged after 269 steps, cumulatively performing nearly 70,000 model evaluations and taking approximately 4 CPU seconds per model evaluation.

#### 3.1.2 Personalized passive mechanics from iFEA

The HO model parameters (*a, b, a*_*f*_, and *b*_*f*_) optimized using the iFEA framework are marked by a red dagger *†* in Table 6. Figure 11A shows the estimated reference configuration Ω_*X*_, colored by displacement magnitude relative to the loaded configuration Ω_RR70_ (shown in transparent gray). The largest displacements occur at the ventricular septum and the basal region of the RV free wall.

**Table 6.** 3D model parameters. Values optimized using the methods described in Section 2.4.2 and Section 2.4.3 are marked by a red dagger *†*.

| Parameter | Symbol | Value | Units |
| --- | --- | --- | --- |
| **General parameters** |  |  |  |
| Time step size | $\Delta t$ | $10^{-3}$ | s |
| Number of cardiac cycles | $N_{\text{cycles}}$ | 5 | |
| Tissue density | $\rho$ | $1.055 \times 10^3$ | $\text{kg m}^{-3}$ |
| Tissue viscosity | $\mu_v$ | $10^2$ | $\text{Pa s}$ |
| Tissue bulk modulus | $\kappa$ | $10^6$ | Pa |
| **Boundary conditions (Robin)** |  |  |  |
| Maximum epicardial stiffness | $k_{\text{epi}}^{\text{max}}$ | $10^7$ | $\text{Pa m}^{-1}$ |
| Minimum epicardial stiffness | $k_{\text{epi}}^{\text{min}}$ | $10^3$ | $\text{Pa m}^{-1}$ |
| Apicobasal steepness | $\alpha$ | 5 | |
| Apicobasal midpoint | $z_0$ | 0.7 | |
| RV-to-LV stiffness ratio | $\lambda_{\text{r/l}}$ | 0.1 | |
| Epicardial damping | $c_{\text{epi}}$ | $5 \times 10^3$ | $\text{Pa s m}^{-1}$ |
| Valve ring stiffness | $k_{\text{valves}}$ | $10^5$ | $\text{Pa m}^{-1}$ |
| Valve ring damping | $c_{\text{valves}}$ | $5 \times 10^3$ | $\text{Pa s m}^{-1}$ |
| **Myocardium** |  |  |  |
| Isotropic stiffness | $a$ | $164.8^\dagger$ | Pa |
| Isotropic exponential | $b$ | $1.723^\dagger$ | Pa |
| Fiber stiffness | $a_f$ | $777.3^\dagger$ | Pa |
| Fiber exponential | $b_f$ | $6.980^\dagger$ | Pa |
| Sheet stiffness | $a_s$ | 2481 | Pa |
| Sheet exponential | $b_s$ | 11.12 | Pa |
| Fiber-sheet stiffness | $a_{fs}$ | 216.0 | Pa |
| Fiber-sheet exponential | $b_{fs}$ | 11.44 | Pa |
| Heaviside smoothness | $k_\chi$ | 100 | |
| **Valvular tissue** |  |  |  |
| Isotropic stiffness | $a_{\text{valves}}$ | $10^6$ | Pa |
| Isotropic exponential | $b_{\text{valves}}$ | 5 | Pa |
| **Active contraction** |  |  |  |
| Active stress magnitude<br>(See table D.1 for timing parameters) | $\tau_{\text{max}}$ | $70^\dagger$ | kPa |

**Figure 11.**
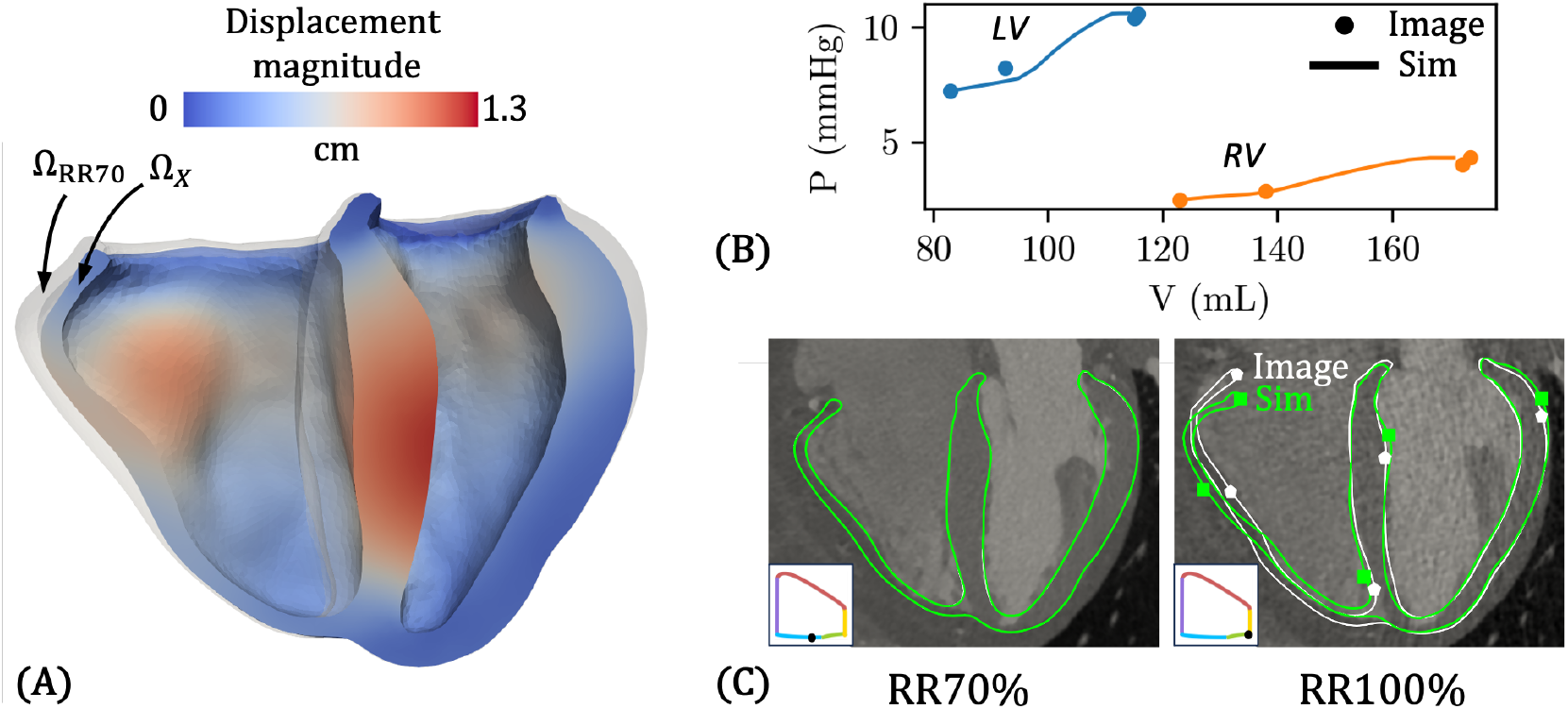
Results from characterizing biventricular passive mechanics using inverse finite element analysis (iFEA). **A**: The estimated reference configuration Ω_*X*_, colored by the magnitude of displacement from the RR70% configuration Ω_RR70_, shown in transparent gray. The largest displacements are at the ventricular septum and the basal portion of the RV free wall. **B**: The simulated late-diastolic passive PV curve (lines) over the target data (dots) using the estimated reference configuration and optimized material parameters. **C**: 4-chamber slice view showing the simulated deformation (green squares) over the image-derived target deformation (white pentagons) at the beginning (RR70%) and end (RR100%) of late-diastolic passive inflation. The background is the CTA image. The black dot on the inset PV loop denotes the phase of the cardiac cycle.

Figures 11B–C illustrate the effectiveness of iFEA in personalizing the passive mechanical response. In Figure 11B, we show the pressure–volume (PV) curves for the LV and RV during late-diastolic filling (RR70% to RR100%). Dots indicate the target PV data at RR70%, RR80%, RR90%, and RR100%, with volumes derived from CTA images and pressures from the tuned 0D model (Figure 10B). The simulated PV curves, obtained using the optimized HO parameters and Ω_*X*_, are shown as lines. These curves show good agreement with the target data and exhibit a physiologic concave-up shape [133] across most of the loading range, with some deviation at the end.

While Figure 11B confirms good agreement in chamber volumes, we also find a decent agreement in the overall myocardial deformation. In Figure 11C we compare simulated deformation (green) to image-derived deformation (white), overlaid on the raw CTA image in a 4-chamber view, defined by a plane containing the apex, mitral valve center, and tricuspid valve center [95]. The white boundary is obtained from the image-morphed BiV surface meshes Γ_img,RR*i*_, *i* ∈ [0, 10, …, 90], described in Section 2.2 and visualized in Figure 4. At RR70%, the simulated myocardium perfectly aligns with the image data. At RR100%, the agreement remains strong, though some discrepancies remain at the apical and basal regions of the RV free wall.

#### 3.1.3 Results from active stress parameter sweep

The results from sweeping the active stress magnitude parameter (*τ*_max_) are shown in Figure 12. Figures 12A-B show the temporal traces of important left and right side pressures and volumes for values of *τ*_max_ between 40 kPa and 80 kPa. As expected, as *τ*_max_ increases, the maximum value of *p*_LV_ increases, as does the maximum value of 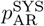 (Figure 12A). At *τ*_max_ = 40 kPa, the LV does not contract quickly and strongly enough to raise 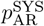to the target value of 98 mmHg, denoted by the upper horizontal green dotted line. However, with *τ*_max_ = 60 kPa, the peak of 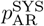 is already very close to the target value. Interestingly, the minimum value of 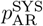 appears relatively unaffected by *τ*_max_, with all cases slightly undershooting the target value of 53 mmHg, denoted by the lower horizontal green dotted line. Similar trends are observed on the right side of the heart (Figure 12B).

**Figure 12.**
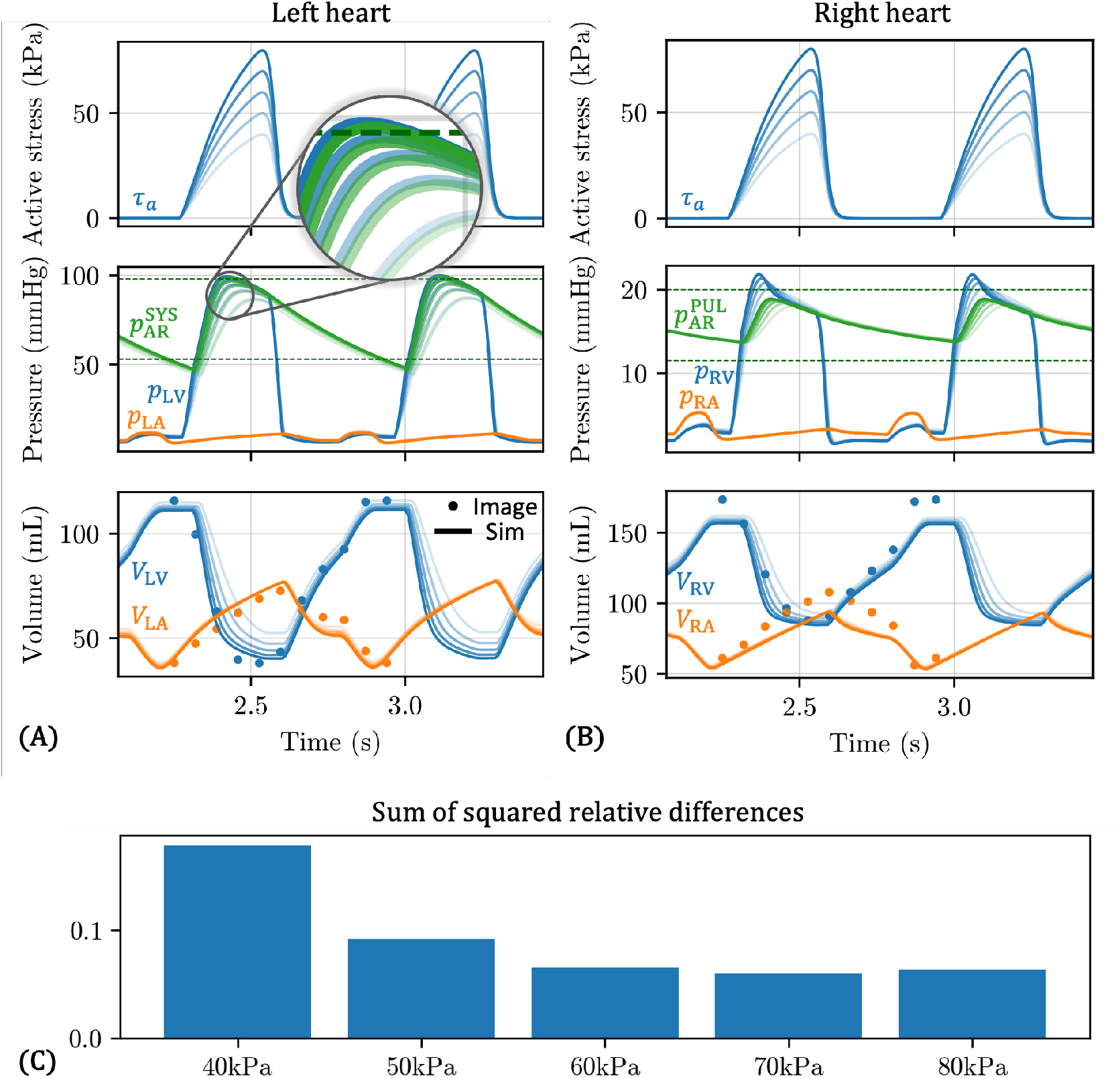
Results from varying the maximum active stress parameter (*τ*_max_), incorporating optimized 0D parameters (Table 5), estimated reference configuration (Ω_*X*_ in Figure 11A), and optimized passive material parameters (Table 6). *τ*_max_ ranges from 40 kPa (lightest lines) to 80 kPa (darkest lines) in 10 kPa increments. All simulations were run for five cardiac cycles, and data was extracted from the last two cardiac cycles. **A**: Left heart active stress, pressures, and volumes. Top: Double Hill active stress curve *τ*_*a*_ (Eq. (15)). Middle: LV pressure *p*_LV_ (blue), LA pressure *p*_LA_ (orange), and systemic arterial pressure 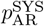 (green). A magnified inset is provided to better distinguish *p*_LV_, which is in all cases slightly greater than 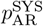. Horizontal dark green dotted lines represent the target minimum and maximum values for 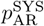. Bottom: LV volume *V*_LV_ (blue) and LA volume *V*_LA_ (orange). Dots represent the target image-derived volume data for RR0%, RR10%, …, RR100%. **B**: Right heart active stress, pressures, and volumes. Top: Double Hill active stress curve *τ*_*a*_ (Eq. (15)). The same *τ*_*a*_ is used for the RV as for the LV. Middle: RV pressure *p*_RV_ (blue), RA pressure *p*_RA_ (orange), and pulmonary arterial pressure 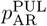 (green). Horizontal dark green dotted lines represent the target minimum and maximum values for 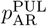. Bottom: RV volume *V*_RV_ (blue) and RA volume *V*_RA_ (orange). Dots represent the target image-derived volume data for RR0%, RR10%, …, RR100%. **C**: The sum of squared relative differences between clinical targets and model predictions (Eq. (22)) for each value of *τ*_max_. A magnitude of 70 kPa yields the best fit.

We also observe an intuitive relationship between *τ*_max_ and LV and RV volumes. As *τ*_max_ increases, the minimum values of *V*_LV_ and *V*_RV_ decrease. Furthermore, the rate of volume change also increases, which explains the increase in the peak *p*_LV_ and *p*_RV_ with *τ*_max_; higher flowrates out of the LV and RV lead to higher LV and RV pressure. Interestingly, *τ*_max_ appears to have a minimal effect on the maximum volumes. It also has minimal impact on atrial volumes, although it is worth noting that in this model, the atria are only represented by 0D TVE elements.

In Figure 12C, we quantify the overall agreement with the target pressure and volume data for each value of *τ*_max_ by computing the sum of squared relative differences, which was also used as the objective function for tuning the full 0D surrogate (Eq. (22)). We find that *τ*_max_ = 70 kPa provides the overall best fit to data.

### 3.2 Personalized multiscale BiV model

In this section, we evaluate the optimized multiscale BiV model in terms of the agreement with target pressure and volume data (Figure 13), the agreement with the image-based myocardial deformation (Figure 14), and myocardial stress and strain (Figure 15). To reiterate, the parameters of this model are listed in Tables 5, 6, and D.1, where the values marked by a red dagger *†* have been optimized using our multistep personalization procedure (Section 2.4). The simulation was run for five cardiac cycles, and data was extracted from the last two cardiac cycles.

**Figure 13.**
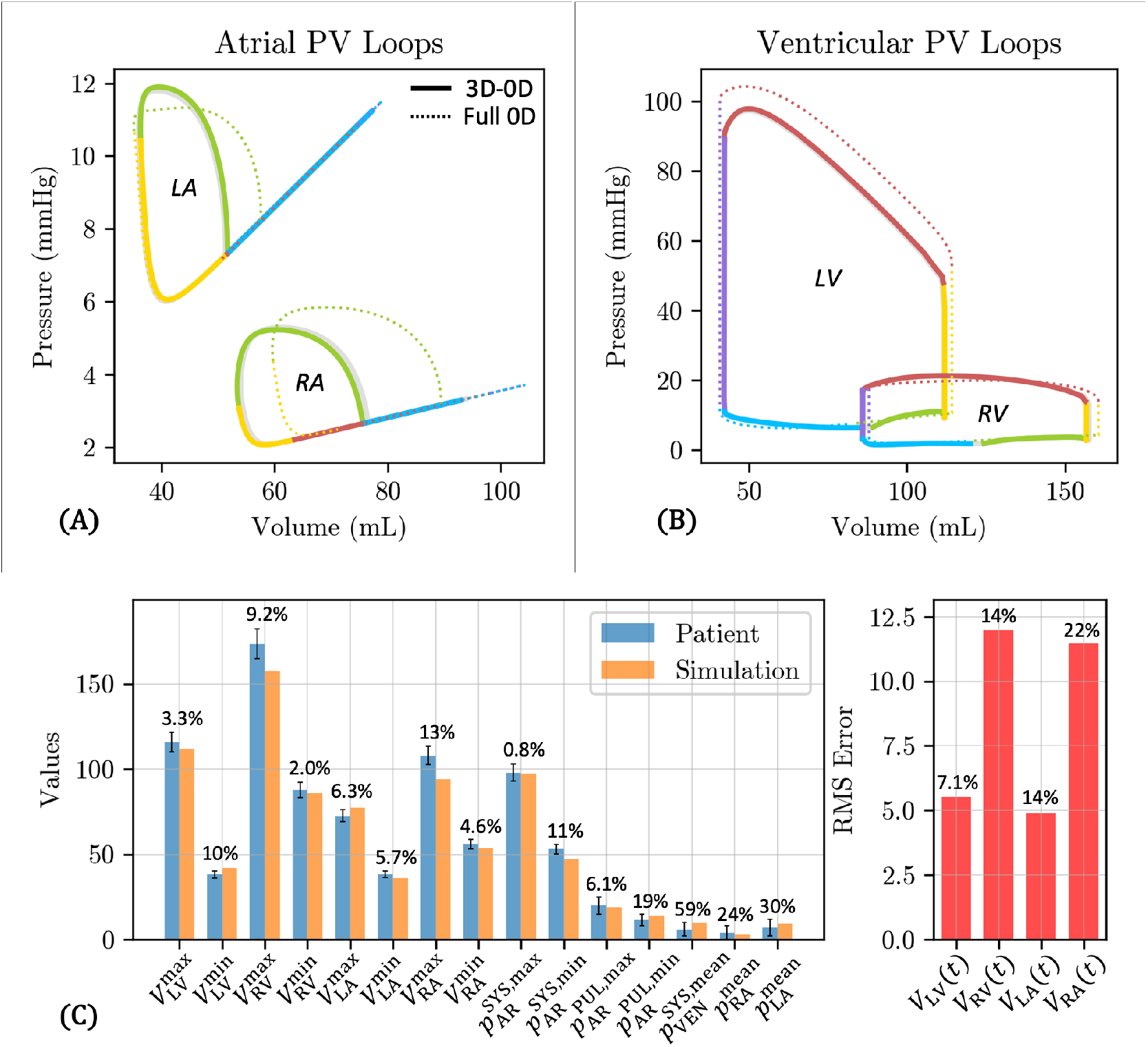
Pressure and volume outputs of the personalized coupled multiscale (3D-0D) BiV model. The simulation was run for five cardiac cycles, and data was extracted from the last two cardiac cycles. **A-B**: PV loops for LA, RA, LV, and RV (solid lines). For reference, we also plot the full 0D model results (identical to Figure 10A-B) in dotted lines. Colors indicate the phase of the cardiac cycle: atrial contraction (green), isovolumic contraction (yellow), ventricular ejection (red), isovolumic relaxation (purple), and ventricular passive filling (blue). **C**: (Left) Comparison between simulation outputs (orange) and scalar clinical targets (blue) for the studied patient. Error bars indicate the uncertainty in target patient values. (Right) RMS errors for the chamber volumes represent the average volume error across all imaged cardiac phases (RR0%, …, RR90%) and are computed using Eq. (25). *V*_(_·_)_ quantities in mL and *p*_(_·_)_ quantities are in mmHg. Percentage values above each bar indicate the percent relative error (Eqs. (23) and (24)).

**Figure 14.**
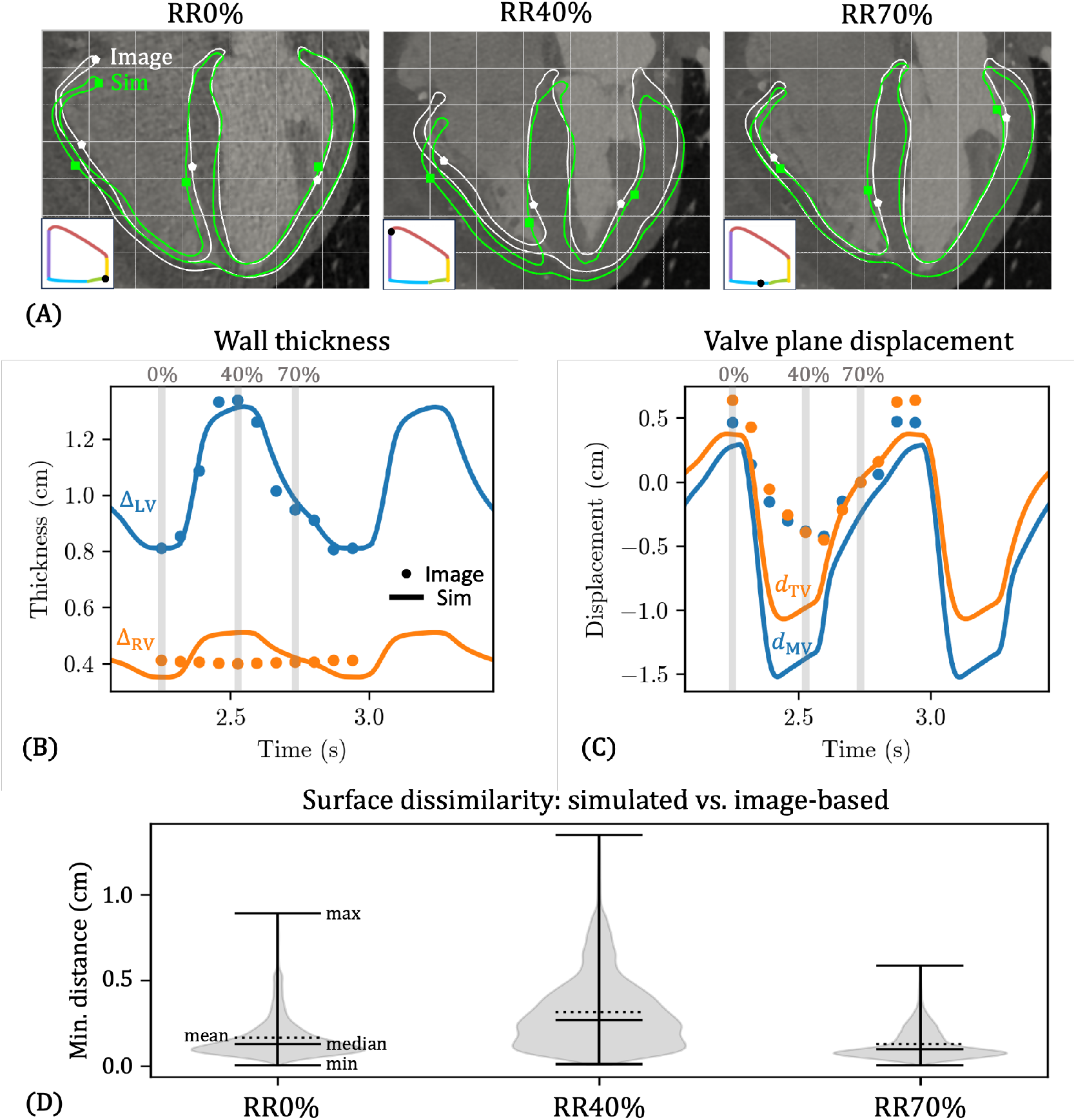
**A**: 4-chamber slice view showing the deformation of the personalized 3D-0D BiV model, at three phases during the cardiac cycle: RR0% (max volume), RR40% (min volume), and RR70% (diastasis at which the model was constructed). The image-morphed BiV surface (Γ_img,RR*i*_, *i* ∈ [0, 40, 70]) is shown in white with pentagon markers, while the corresponding simulated surface (Γ_sim,RR*i*_, *i* ∈ [0, 40, 70]) is shown in green with square markers. The background is the CTA image. Light gray grid lines provide a fixed reference to observe deformation patterns. The black dot on the inset PV loop denotes the phase of the cardiac cycle. Animations are provided in Supplemental Data. **B**: LV (Δ_LV_) and RV (Δ_RV_) wall thicknesses versus time (two cardiac cycles). **C**: Mitral valve (*d*_MV_) and tricuspid valve (*d*_TV_) plane displacements versus time. In panels **B** and **C**, lines indicate quantities extracted from the personalized BiV simulation, while dots represent quantities obtained from image-morphed BiV surface. **D**: To quantify the dissimilarity between image-based (white) and simulated (green) deformation, we generate violin plots of the distribution of distances from each node on the green boundary to the nearest node on the white boundary, *H*(Γ_img,RR*i*_; Γ_sim,RR*i*_) for *i* ∈ [0, 40, 70] (Eq. (26)). Note that while we visualize the green and white boundaries with 2D slices, the distance computation is performed with the analogous 3D surfaces. Solid horizontal lines indicate the minimum, median, and maximum of the distance histogram, while the dotted line indicates the mean.

**Figure 15.**
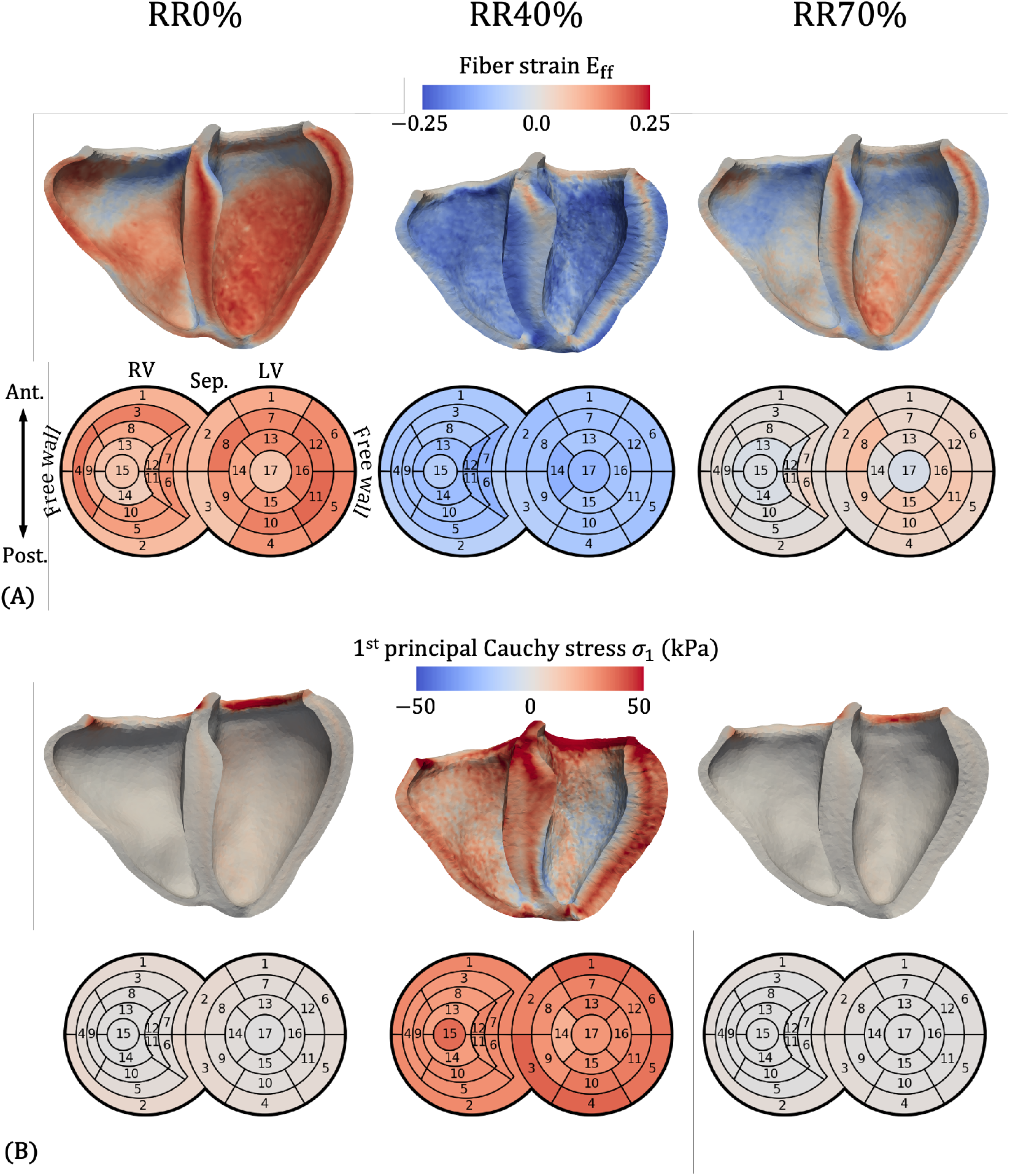
Fiber strain (**A**) and 1^st^ principal Cauchy stress (**B**) of the personalized 3D-0D BiV model, at three phases during the cardiac cycle. Strain and stress are also shown on BiV bullseye plots, using a standard 17-segment model for the LV [138] and a 15-segment model for the RV [139].

#### 3.2.1 0D outputs

Figure 13 shows pressure and volume outputs. Note that this data is identical to that shown in Figure 12 for the 70 kPa case, but we visualize the results in terms of PV loops (Figure 13A-B) and provide a more detailed breakdown of the model fit to target pressure and volume data (Figure 13C). The atrial PV loops for the 3D-0D model (solid lines) are similar to those in the full 0D model (dotted lines), since the atria are modeled the same in both cases, namely with linear TVE elements. However, the a-loops for both LA and RA are shifted to lower volumes, and the maximum volume for the RA is also reduced (94.0 mL vs. 104.2 mL, 9.8% smaller). The ventricular PV loops also resemble their full 0D counterparts, albeit with some differences. The LV PV loop exhibits slightly smaller maximum volume (111.9 mL vs. 114.2 mL, 2.0% smaller) and slightly greater minimum volume (42.0 mL vs. 40.7 mL, 3.2% larger), as well as a reduction in peak systolic pressure (97.9 mmHg vs. 104.3 mmHg, 6.1% smaller). The RV reaches a slightly higher systolic pressure than the full 0D model (21.4 mmHg vs. 20.0 mmHg, 7% larger), but is shifted to smaller volumes. The resulting stroke volumes for the LV (70.0 mL vs. 73.5 mL, 4.8% smaller) and the RV (71.7 mL vs. 72.6 mL, 1.2% smaller) are both slightly lower than for the full 0D model. These correspond to ejection fractions (EFs) of 62.5% for the LV and 45.5% for the RV. Additionally, in the 3D-0D model, the ventricular PV loops, especially the LV, display a change in concavity between the passive filling phase (blue) and the atrial contraction phase (green), which is not apparent in the full 0D model data (see also Figure 10B).

Figure 13C shows a decent agreement with the target clinical data, although the fit is not as good as the full 0D model (Figure 10C). In matching the measured patient data, over 5% errors are found for the minimum LV volume (10%, 3.8 mL), the maximum RV volume (9.2%, 15.9 mL), the maximum RA volume (13%, 13.9 mL), and the minimum arterial systemic pressure (11%, 5.8 mmHg). Higher relative errors are obtained for the reference pressure values 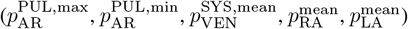, but as discussed previously, these quantities have a greater uncertainty in the literature (Table 3) and are used in this study only to encourage overall physiological behavior. Likewise, although the normalized RMS errors are marginally higher than those of the full 0D model, they are still within acceptable limits. *V*_RV_ (14%) and *V*_RA_ (22%) show the largest disagreement, primarily attributed to the underprediction of the maximum values for these quantities. These discrepancies can also be observed in Figures 12A-B for the 70 kPa case, which provide a good qualitative understanding of how well our model outputs align with the target pressure and volume data.

#### 3.2.2 Myocardial deformation

In Figure 14A, we compare the myocardial deformation predicted by our model (green) with that derived from clinical CTA image data (white), overlaid on the gray-scale CTA image in a 4-chamber view. The white boundary corresponds to the image-morphed BiV surface meshes Γ_img,RR*i*_, *i* ∈ [0, 10, …, 90], described in Section 2.2 and shown in Figure 4. We analogously denote the simulated BiV surface meshes (green boundary) as Γ_sim,RR*i*_. At RR0% (end-diastole), the agreement between the simulated and image-based geometries is generally good. Some discrepancies are visible near the apex and base of the RV free wall, as well as at the apical portion of the septum. At RR40% (end-systole), the agreement deteriorates, particularly toward the apex and the RV free wall. Nonetheless, the simulation captures the overall deformation pattern from RR0% to RR40%, including downward (apical) displacement of the atrioventricular (AV) plane, systolic wall thickening, and radial contraction of the free walls. However, the magnitude of AV plane displacement is overestimated by the model, especially on the LV and RV free walls. Additionally, the image-based deformation shows upward (basal) motion of the epicardial apex, especially on the RV side. In contrast, the RV apex in the simulation displaces downward, while the LV apex remains virtually stationary. From RR40% to RR70% (approximately the ventricular early filling or E-wave), the deformation reverses direction: the AV plane displaces upward, wall thickness decreases, and the free walls expand radially. The simulation reproduces these features reasonably well. At RR70%, the match between simulation and image is strongest, with only minor discrepancies at the basal regions of the LV and RV. Finally, from RR70% back to RR0%, corresponding to the late diastolic phase, we observe continued upward displacement of the AV plane and radial expansion of the free walls. These motions are again captured by the model, although the predicted AV plane displacement on the RV free wall, as well as radial expansion of the apical RV free wall, falls short of the image-based deformation.

To quantify the deformation, we measured the wall thicknesses of the LV (Δ_LV_) and the RV (Δ_RV_); the method to compute wall thicknesses is described in Appendix C. Figure 14B shows simulated (lines) and image-based (dots) Δ_LV_ and Δ_RV_ over two cardiac cycles, reinforcing the qualitative patterns observed in Figure 14A. For the LV, the image-derived systolic wall thickening is approximately 65% from RR0% to RR40%, which is in the range of previously reported experimental values [134, 135, 136]. Both the magnitude and shape are matched very well by the BiV simulation. For the RV, the image-derived wall thickness shows virtually no change over the cardiac cycle. This is because the automatic segmentation tool, MeshDeformNet, only tracks the RV endocardium, causing the RV wall thickness to remain nearly constant (a nominal thickness of 3.5 mm was prescribed in Section 2.2, but the actual thickness is approximately 4.0 mm after smoothing). The simulation, however, does predict RV systolic thickening, which is approximately 46% in our model.

We also calculated the displacements of the mitral valve (*d*_MV_) and tricuspid valve (*d*_TV_) planes (Appendix C). Figure 14C shows *d*_MV_ and *d*_TV_ over two cardiac cycles, comparing simulation results (lines) with image-derived motion (dots). Both datasets show the characteristic systolic downward displacement from RR0% to RR40%, followed by an upward rebound during early diastole (RR40–70%) and continued upward motion through late diastole (RR70–100%). The simulation captures the overall shape of the image-based displacement curves, although the peak systolic displacement occurs slightly earlier. It also overestimates the magnitude of systolic AV plane motion, particularly for the MV plane, consistent with the qualitative observations in Figure 14A. Similarly, during late diastole, the simulated increase in both *d*_MV_ and *d*_TV_ follows the image-based trends but remains smaller in magnitude.

To quantify the dissimilarity in myocardial boundary deformation between the image-based BiV data(Γ_img,RR*i*_) and simulated BiV model (Γ_sim,RR*i*_) in 3D, we compute the set of directed minimum distances (DMDs) from Γ_sim,RR*i*_ to Γ_img,RR*i*_. Dropping the subscript RR*i* for the moment, we define the set of DMDs as

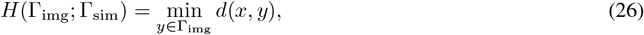

where *x* and *y* are points of Γ_sim_ and Γ_img_, respectively, and *d*(*x, y*) is the Euclidean distance between points *x* and *y*. Note that the directed Haussdorff distance (DHD) [137] can be obtained as the maximum of *H*(Γ_img_; Γ_sim_) over all points *x* ∈ Γ_sim_. Figure 14D visualizes *H*(Γ_img,RR*i*_; Γ_sim,RR*i*_) for *i* ∈ [0, 40, 70] using violin plots, where the width of the violin plot indicates the frequency of distances over all points *x* ∈ Γ_sim_. As seen qualitatively in the 4-chamber slice views (Figure 14A), the deformation agreement is worst at RR40% (mean 0.314 cm, max 1.347 cm), best at RR70% (mean 0.128 cm, max 0.586 cm), and intermediate at RR0% (mean 0.164 cm, max 0.892 cm). Moreover, the distributions of distances are skewed toward larger values, as evidenced by median values below mean values. This indicates that while most of Γ_sim,RR*i*_ lies relatively close to Γ_img,RR*i*_, some parts of the surface lie significantly further away, which is also apparent in the 4-chamber slice views (Figure 14A).

#### 3.2.3 Strain and stress

Figure 15A shows fiber strain (*E*_ff_ = **f**_**0**_ · (**Ef**_**0**_)) in a long-axis anterior cut view (top) and a bullseye plot (bottom). *E*_ff_ measures the change in length of myocardial fibers (*l*) relative to their length in the reference configuration (*L*). Specifically, 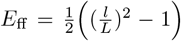. The bullseye plot uses the standard 17-segment model for the LV [138] and a 15-segment model for the RV adapted from Tokodi et al. [139]. From the clipped views, we note a transmural variation in *E*_ff_ in all cardiac phases, particularly in the septum and the LV free wall. At end-diastole (RR0%) and diastasis (RR70%), the midwall portion of the LV free wall, where fibers are oriented roughly circumferentially, experiences the greatest fiber strain. At end-systole (RR40%), the midwall strain is weakly positive, while the endocardial and epicardial layers, where fibers are oriented at *±*60 relative to circumferential, show strong contraction up to −0.25 strain. At all phases, the most basal portions of both the LV and RV show little strain. This corresponds to Ω_ring_, where a stiff isotropic material was prescribed to model the valvular tissue in that region. From the bullseye plot, we note some regional variations in *E*_ff_. At RR0%, the highest fiber strain is seen in the LV free wall mid-cavity segment 11 (0.178). The lowest fiber strain is at the RV posterior apical segment 14 (0.0445), and the mean across all LV and RV segments is 0.117. At RR40%, the highest magnitude strain is at the RV septal apical segment 12 (−0.135), while the lowest is at RV segment 2 (−0.0625). The mean across all segments is −0.0928. LV segment 11 experiences the greatest change in strain from RR0% to RR40% (0.261), while LV segment 3 experiences the smallest change (0.127), and the mean change in *E*_ff_ is 0.210.

Figure 15B shows the 1^st^ principal Cauchy stress *σ*_1_, representing the maximum tensile stress over all directions experienced by the myocardium at that point. At RR0% and RR70%, most of the tissue has relatively low stress, except for the Ω_ring_ region at the LV and RV base. We also note a slight transmural variation in stress, which becomes more pronounced at RR40%. In line with the fiber strain patterns, the midwall portions of the LV free wall and septum experience greater stress than the endocardial and epicardial layers. Interestingly, at RR40%, the mid-to-lower ventricular portions of the LV endocardial layer actually experience negative stress, indicating that this material is being compressed. From the bullseye plot, we note some regional variations in stress. At RR0%, the highest stress is at RV segment 2 (4.42 kPa), while the lowest is at LV segment 17 (0.127 kPa). The mean across all segments is 1.77 kPa. At RR40%, the highest stress is at LV segment 3 (37.1 kPa), and the lowest is at LV segment 14 (18.8 kPa). The mean across all segments is 2.89 kPa. LV segment 14 also experiences the smallest change in stress from RR0% to RR40% (17.6 kPa), while the largest change is found in RV segment 15 (34.5 kPa).

### 3.3 BiV vs. t-BiV vs. LV comparison

Figure 16 compares the simulation results for the three anatomical models: BiV (green), t-BiV (orange), and LV (teal). For t-BiV, we choose the case with the same value of the basal Robin stiffness as applied on the valve surfaces of the BiV model, denoted t-BiV(same base) (Table 4). For the LV-only model, we choose the case with a Robin BC over the entire epicardial surface, including the septal portion, denoted LV(all Robin) (Table 4). We reiterate that all model inputs are identical, except for the geometry and boundary conditions as stated (Section 2.5). All simulations were run for five cardiac cycles, and data is extracted from the last two cardiac cycles for further comparison.

**Figure 16.**
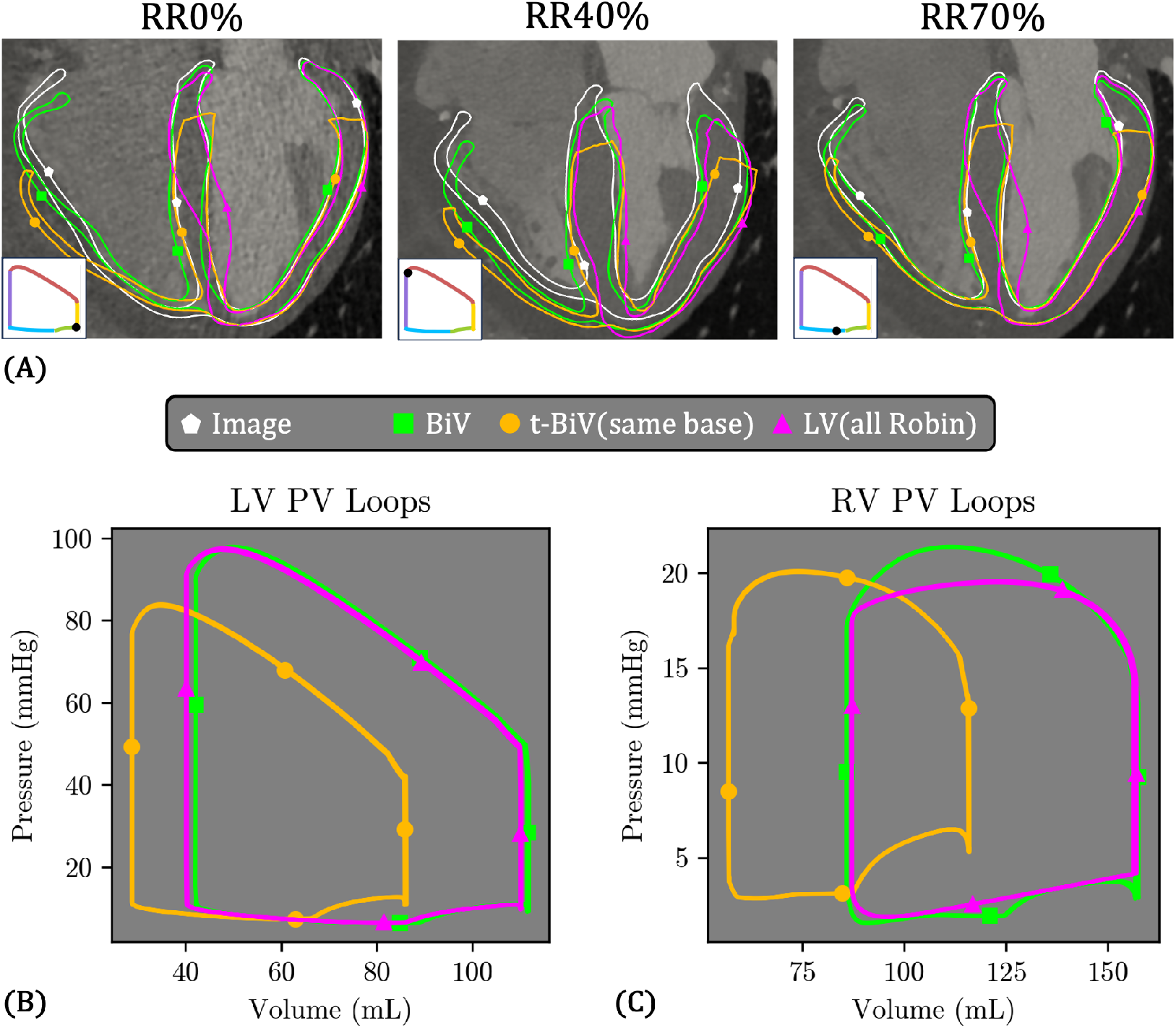
Comparison of the simulated local deformation and global hemodynamics among the three anatomical models considered in this work. Green squares: BiV, the biventricular model. Orange circles: t-BiV(same base), the truncated biventricular model (t-BiV), with the same value of the Robin BC stiffness applied on the base as is applied on the valve surfaces on BiV. Pink triangles: LV(all Robin), the left ventricle model, with Robin BC applied on the entire epicardium, including the septal portion. All other inputs are identical among the three models. All simulations were run for five cardiac cycles, and data is extracted from the last two cardiac cycles. **A**: 4-chamber slice view showing the deformed configuration of the three models at key phases during the cardiac cycle. For reference, the gray-scale CTA image is shown in the background, and the image-morphed BiV surface (Γ_img,RR*i*_, *i* ∈ [0, 40, 70]) is overlaid in white with pentagon markers. The black dot on the inset PV loop denotes the phase of the cardiac cycle. **B-C**: LV and RV PV loops for the three models, following the same color code used in frame **A**.

Figure 16A shows the deformed configuration of the three models at key cardiac phases. We note some differences in the deformation patterns of t-BiV(same base) and LV(all Robin) compared to the baseline BiV case and the image-derived deformation. At RR0% and RR40%, t-BiV(same base) matches BiV well in the LV free wall and septum, but exhibits excess outward radial displacement in the RV free wall. For the LV(all Robin) model, at both RR0% and RR70%, the lower portion of the septum bulges into the LV cavity. Finally, during systole (RR40%), the epicardial apex of LV(all Robin) moves downward, in contrast to virtually no apex motion in BiV and t-BiV, and a slight upward motion of the apex in the image-based deformation.

PV loops for the three models are shown in Figure 16B–C. In terms of the LV function, LV(all Robin) is remarkably similar to BiV, with only slightly smaller (~2%) minimum and maximum volumes. In contrast, t-BiV(same base) significantly underpredicts minimum (28.7 mL vs. 41.9 mL, 31.5% smaller) and maximum (86.3 mL vs. 112 mL, 22.9% smaller) LV volumes, as well as peak systolic pressure (83.9 mmHg vs. 97.9 mmHg, 14.3% smaller), compared to the BiV model. While not visually obvious, LV stroke volume is also substantially reduced (57.6 mL vs. 70.0 mL, 17.7% smaller), though LV ejection fraction is slightly larger (66.8% vs. 62.5%, 6.9% larger). In terms of RV function, t-BiV(same base) similarly exhibits reduced minimum and maximum RV volumes, as well as reduced peak systolic pressure (20.1 mmHg vs. 21.4 mmHg, 6.1% smaller), although the reduction is much smaller than for the LV. The RV PV loop for LV(all Robin) has a smoother shape, since it is represented by a TVE element in this case, with very similar minimum and maximum RV volumes to the BiV model, but a reduced peak systolic pressure.

### 3.4 t-BiV variations

Due to the excessive deformation observed in t-BiV(same base) (Figure 17A), we simulated an additional case with a tenfold increase in basal stiffness, referred to as t-BiV(stiffer base) (Table 4). Anticipating that this would further reduce predicted volumes and pressures, we also simulated a third case, t-BiV (higher stress), which combines increased basal stiffness with double the magnitude of active stress to enhance contractile function. Results from all three cases are shown in Figure 17.

**Figure 17.**
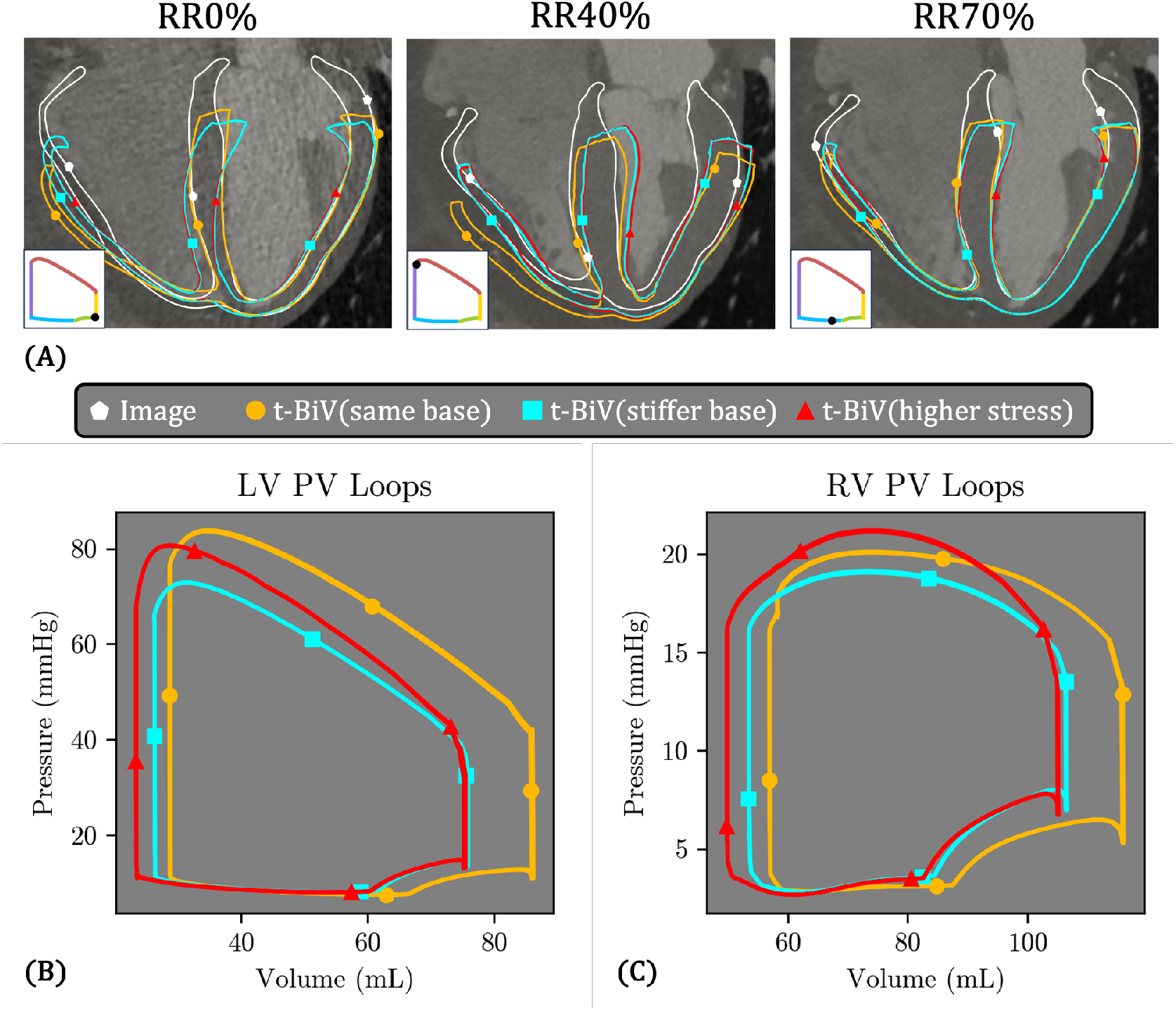
Comparison of the simulation results among the three truncated biventricle (t-BiV) cases considered in this study (Table 4). Orange circles: t-BiV (same base), with the same basal Robin stiffness of *k*_base,t−BiV_ = 10^5^Pa m^−1^ as that of the full biventricle (BiV) model (Figure 16) and an active stress magnitude of *τ*_max,t−BiV_ = 70 kPa. Light blue squares: t-BiV (stiffer base) with *k*_base,t−BiV_ = 10^6^Pa m^−1^ and *τ*_max,t−BiV_ = 70 kPa. Red triangles: t-BiV (higher stress) with *k*_base,t_ − _BiV_ = 10^6^Pa m^−1^ and *τ*_max,t_−_BiV_ = 140 kPa. All simulations were run for five cardiac cycles, and data is extracted from the last two cardiac cycles. **A**: 4-chamber slice view showing the deformed configuration of the three models at key phases during the cardiac cycle. For reference, the gray-scale CTA image is shown in the background, and the image-morphed BiV surface (Γ_img,RR*i*_, *i* ∈ [0,40,70]) is overlaid in white with pentagon markers. The black dot on the inset PV loop denotes the phase of the cardiac cycle.Animations are provided in Supplemental Data. **B-C**: LV and RV PV loops for the three models, following the same color code used in frame **A**.

As expected, both t-BiV(stiffer base) and t-BiV(higher stress) exhibit markedly reduced basal plane motion throughout the cardiac cycle (Figure 17A), effectively limiting the excessive outward radial displacement of the RV free wall observed in t-BiV(same base). However, the increased basal stiffness introduces some unusual deformation patterns. At RR0%, pronounced bending occurs near the basal plane, particularly in the RV. The elevated stiffness also constrains septal motion, resulting in poorer agreement with the image data. Interestingly, despite a two-fold increase in active stress magnitude, the deformations of t-BiV(stiffer base) and t-BiV(higher stress) are very similar at all cardiac phases shown, with only small differences at the septum and RV free wall.

The PV loops for all three models are shown in Figure 17B–C. Compared to t-BiV(same base), the increased basal stiffness in both t-BiV(stiffer base) and t-BiV(higher stress) substantially reduces the maximum LV volume and, to a lesser extent, the minimum volume. Consequently, LV stroke volume (49.7 mL vs. 57.6 mL, 13.7% smaller) and LV ejection fraction (65.5% vs. 66.8%, 1.9% smaller) are diminished in t-BiV(stiffer base), leading to a much lower peak systolic pressure (72.9 mmHg vs. 83.9 mmHg, 13.1% smaller). Comparing t-BiV(higher stress) with t-BiV(stiffer base), the two-fold increase in active stress decreases the minimum LV volume and increases peak systolic LV pressure, but not enough to match t-BiV(same base). Notably, this increase in active stress has minimal impact on the diastolic phase of the PV loop. Similar trends are observed in the RV, except that the peak systolic pressure in t-BiV(higher stress) slightly exceeds that of t-BiV(same base).

### 3.5 LV variations

Figure 18 compares the three LV models, which differ in the boundary conditions applied to the septal epicardial surface Γ_epi,septum_ (Figure 9B). To remind the reader, in LV(all Robin), a spatially varying Robin boundary condition is applied over the entire epicardial surface, including the septum (Table 4). In contrast, LV(free septum) applies no pressure or Robin BC to Γ_epi,septum_, while LV(loaded septum) applies RV pressure on Γ_epi,septum_.

**Figure 18.**
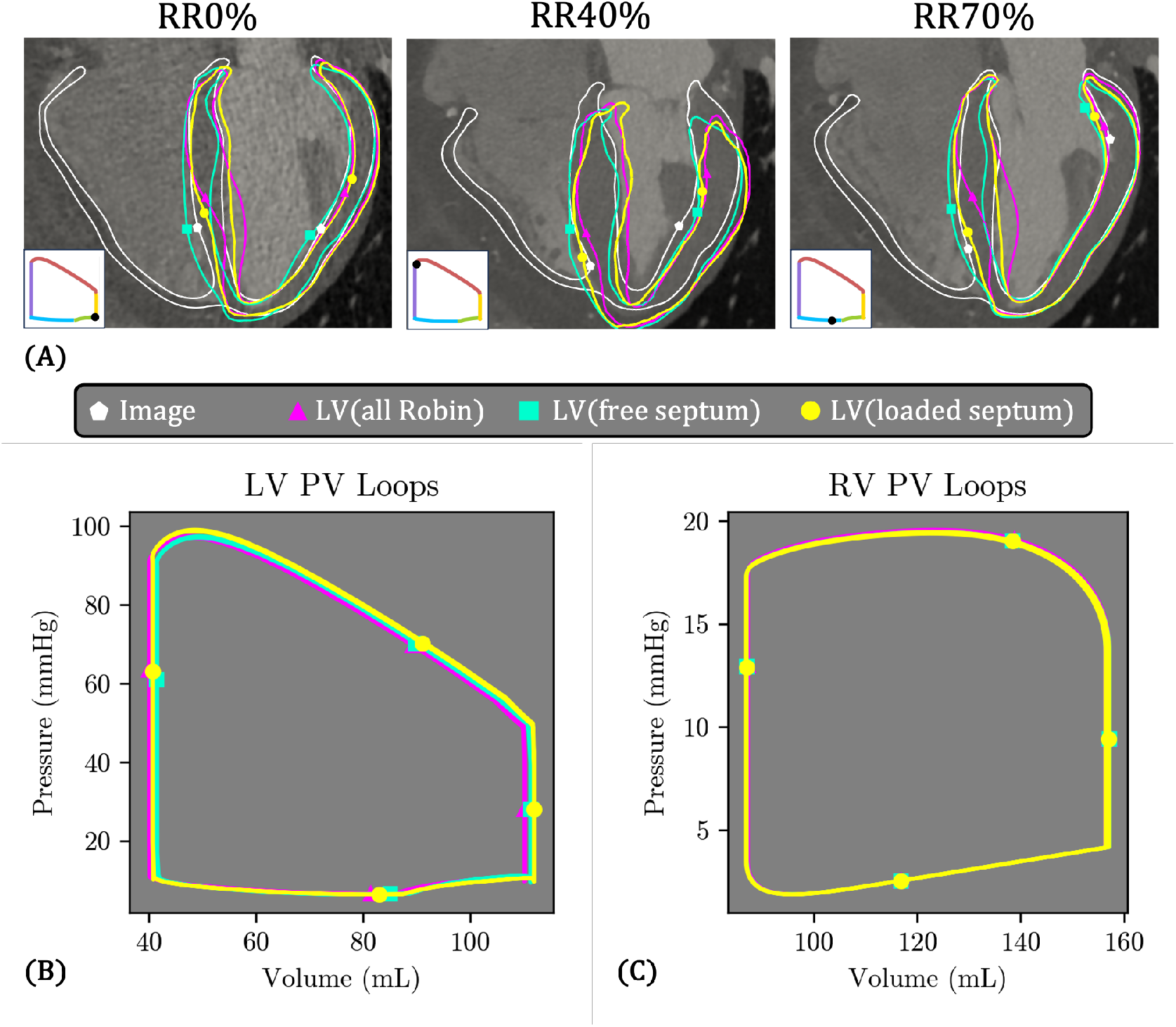
Comparison of the simulation results among the three left ventricular (LV) cases considered (Table 4). Pink triangles: LV(all Robin), with Robin BC on the entire epicardium, including the septal portion. Teal squares: LV(free septum), with no load or Robin BC on the septal epicardium. Yellow circles: LV(loaded septum), with RV pressure applied to the septal epicardium. All simulations were run for five cardiac cycles, and data is extracted from the last two cardiac cycles. **A**: 4-chamber slice view showing the deformed configuration of the three models at key phases during the cardiac cycle. For reference, the gray-scale CTA image is shown in the background, and the image-morphed BiV surface (Γ_img,RR*i*_, *i* ∈ [0, 40, 70]) is overlaid in white with pentagon markers. The black dot on the inset PV loop denotes the phase of the cardiac cycle. Animations are provided in Supplemental Data. **B-C**: LV and RV PV loops for the three models, following the same color code used in frame **A**.

Overall, the deformation patterns are broadly similar across the three models (Figure 18A), though some notable differences emerge. The septal bulging observed in LV(all Robin) is eliminated in LV(free septum), and to a lesser degree in LV(loaded septum). Also, at RR40%, LV(free septum) generally aligns best with the image data, while LV(loaded septum) appears “tilted” toward the left side, and LV(all Robin) is also tilted, but to a lesser degree.

Despite these variations in tissue deformation, the LV and RV PV loops are nearly identical across all three cases (Figure 18B–C).

## 4 Discussion

In this study, we developed a patient-specific model of BiV cardiac mechanics. The model closely reproduces the patient’s cuff blood pressure and phase-resolved chamber volumes, with reasonable agreement in myocardial deformation compared to gated CTA images. Although localized discrepancies in deformation are observed, particularly at end-systole, the model captures essential physiological features, including atrioventricular plane displacement, radial contraction, and wall thickening. We also observed interesting transmural and regional variations in myocardial stress and strain. To investigate the impact of frequently adopted anatomical simplifications, we compared the BiV model with the t-BiV model, which truncates the BiV above the basal plane, and the LV model, which excludes the RV. Analyzing the t-BiV model, we found that truncation not only reduces total ventricular volumes, but also reduces stroke volumes and peak systolic pressures. Additionally, applying the same basal BC stiffness to the t-BiV model resulted in unrealistic deformation of the RV free wall. While increasing the basal stiffness ameliorated this issue, it introduced new, non-physiological deformation patterns near the basal plane. Finally, doubling the active stress magnitude increased peak systolic pressure, but not enough to match the clinical target. Considering the LV model, we found that while removing the RV resulted in moderate differences in deformation, particularly in the apical septum, it had virtually no effect on the LV PV loop. Similarly, variations in BCs applied to the septal epicardium in the LV model produced notable changes in local deformation, but had a negligible impact on global pressure-volume behavior. We conclude that a carefully designed LV-only model may serve as a reasonable surrogate for a BiV model in capturing both global and local LV function, depending on the problem complexity and clinical application. However, truncation at the basal plane substantially distorts global hemodynamics and regional wall mechanics and should therefore be used with caution.

### 4.1 Multistep personalization procedure

Our multistep personalization strategy proved to be an effective and efficient approach for identifying key patient-specific model parameters. By initially replacing the 3D finite element BiV model with simplified TVE representations, we were able to calibrate circulatory (0D) parameters independently, prior to tuning myocardial passive and active mechanical properties. This surrogate-based strategy significantly reduced computational cost and complexity, and was found to be sufficiently accurate for pressure-volume calibration, particularly when embedded in a multistep optimization workflow. The effectiveness of the TVE representation is likely enhanced by its use of the same activation function as the 3D mechanics model, namely the double Hill model, which provides a more realistic description of activation than earlier TVE formulations [87]. The TVE representation of the ventricles is much simpler than neural network–based surrogates used in Salvador et al. [30], which require extensive training data. Although we found linear TVEs to be adequate in this work, the ventricles can also be represented with more complex reduced-order models at little additional computational cost [33, 140, 141, 142, 143].

Sequentially personalizing the passive mechanical response (HO model parameters and reference configuration), then tuning the active stress magnitude *τ*_max_, was effective because *τ*_max_ has a relatively minor influence on diastolic function. This is evident in the active stress magnitude parameter sweep (Figure 12A-B), which shows approximately constant end-diastolic volume and pressure for both LV and RV over a range of *τ*_max_ from 40 kPa to 80 kPa. This behavior is also evident in the comparison between t-BiV(stiffer base) and t-BiV(higher stress); despite a twofold increase in *τ*_max_, the diastolic portions of the PV loops are nearly identical. Instead, *τ*_max_ predominantly affects systolic function, being correlated with decreased end-systolic volume and increased stroke volume and peak systolic pressures (Figure 12A-B). Diastolic function, instead, is primarily influenced by the passive material properties and reference configuration, which were effectively personalized using the iFEA framework (Figure 11). These observations are consistent with the sensitivity analysis performed by Niederer et al. [79].

Our estimated HO model parameters (*a* = 0.1648 kPa, *b* = 1.723, *a*_*f*_ = 0.7773 kPa, and *b*_*f*_ = 6.980), marked by a red dagger *†* in Table 6, are broadly consistent with values reported in other *in vivo* inverse modeling studies, although notable differences persist across the literature. Holzapfel et al. [104] fit their HO model to *ex vivo* shear data [144], and obtained the following parameter values: *a* = 0.057 kPa, *b* = 8.094, *a*_*f*_ = 21.503 kPa, and *b*_*f*_ = 15.819. However, when fitting to biaxial data [145], they obtained substantially different values: *a* = 2.280 kPa, *b* = 9.726, *a*_*f*_ = 1.685 kPa, and *b*_*f*_ = 15.779. *In vivo* stiffnesses obtained by inverse modeling approaches are generally smaller than *ex vivo* values [146, 147]. Gao et al. [148], for example identified *in vivo* HO model parameters of *a* ≈ 0.1 kPa, *b* ≈ 3, *a*_*f*_ ≈ 3 kPa, *b*_*f*_≈ 5. Palit et al. [147] and Shi et al. [111] obtained roughly similar values using inverse finite element analysis on human clinical data. Other studies employing the transversely-isotropic Guccione model for myocardium [149] have reported *in vivo* stiffnesses of a similar order of magnitude [61, 150]. However, direct comparisons with these studies are complicated by several factors. First, the Fung-type material models exhibit issues with parameter identifiability stemming from correlations among parameters; for example, parameters such as *a* and *b* can trade off to produce similar stress-strain responses [148, 151]. Second, most prior studies assumed the diastasis configuration was stress-free [147, 148]; in contrast, the iFEA method employed in this study simultaneously estimates the reference configuration of the heart and material parameters [111]. Also, previous work neglected epicardial BCs [132, 147, 148]; we instead apply Robin BCs to model the effect of the pericardium, which has been shown to strongly influence cardiac mechanics [27, 42]. Finally, other studies have relied on literature values or generic pressure profiles when performing passive inverse modeling [61, 111], which has been shown to strongly affect estimated stiffnesses [132, 147]; we take a slightly more personalized approach, using the passive pressure profile from the tuned full 0D surrogate. The best option would be *in vivo* ventricular pressure recordings synchronized with the CTA data, as obtained for dogs in Wang et al. [150], but this is typically not possible for human subjects. Despite the substantial differences in parameter values, the key outcome is that the target passive mechanical behavior is well captured in this work, as demonstrated in Figure 11. This aligns with the observation of Gao et al. that fairly large differences in parameter values can produce largely similar mechanical responses [148].

The optimal active stress magnitude estimated in this study (*τ*_max_ = 70 kPa) is comparable to values reported in the literature. Experimental studies of isolated ventricular cardiomyocytes report a peak contractile stress of around 40 kPa to 50 kPa [152, 153, 154], while prior computational studies have typically used higher values. For example, Pfaller et al. [27] calibrated a value of 90.7 kPa for a model with *±*60^°^ fiber orientations and epicardial boundary conditions. Strocchi et al. [42] used a higher value of 125 kPa in a whole-heart model incorporating a spatially varying epicardial boundary condition. Fedele et al. [46] also reported active tension exceeding 80 kPa in certain regions of the ventricles. As with the passive material parameters, direct comparisons are challenging due to substantial methodological differences across studies, including the passive material parameter values themselves, as well as the boundary conditions and underlying clinical datasets used for personalization. Nonetheless, we are satisfied that our tuned value lies within reasonable deviations from other studies while faithfully reproducing the observed clinical data.

### 4.2 Personalized multiscale BiV model

#### 4.2.1 0D outputs

In the diastolic portion of the ventricular PV loops (Figure 13B), the 3D-0D BiV model exhibited concave-down curvature during the atrial contraction phase, in contrast to the straight line in the full 0D model, which reflects the linear TVE model used (Eqs. A.4–A.5). Similar late-diastolic concave-down behavior has been reported in the biventricular model of Augustin et al. [52] as well as the whole-heart model of Fedele et al. [46]. This has also been observed experimentally and described as “excess pressure” during atrial systole for a given volume [155], being attributed to the viscous properties of the tissue. In our case, rate-dependent deformation could originate from myocardial viscosity in the material model (Eq. 17) or damping within the Robin boundary conditions (Eq. 10).

The multiscale BiV model reproduced the target pressure and volume data well, though not perfectly (Figures 12 and 13). The underprediction of RV maximum volume (Figure 13) may be partly due to errors in calculating image-based chamber volumes. Indeed, while LV and RV stroke volumes were nearly identical in our model (70.0 mL and 71.7 mL), stroke volumes based on the CTA image data were larger and more unequal (77.6 mL and 85.5 mL). In healthy subjects, LV and RV stroke volumes should be very similar [156]. Differential stroke volume can occur, but this is usually associated with valve regurgitation, which was not reported for the studied patient [156]. Because regurgitation was negligible in our model, we found that the model tends to equalize stroke volumes, as reported in other computational studies with closed-loop circulatory models [33], making it difficult to match clinical data for which LV and RV stroke volumes differ. Image-based volume errors may arise from whether papillary muscles and trabeculae are included or excluded during segmentation, as this choice significantly influences chamber volume estimation [157] and can bias LV-to-RV comparisons since LV papillary muscles are typically larger than those in the RV. The deep-learning segmentation method used in the present work generally excludes the papillary muscles and most of the trabeculae from the myocardial segmentation, although these criteria depend on its training data.

#### 4.2.2 Myocardial deformation

From RR0% to RR40% (isovolumic contraction and ejection), we reproduced physiological patterns of LV deformation observed in the CTA data, including AV plane displacement, wall thickening, and radial contraction of the free wall [27, 158] (Figure 14A-C). Greater longitudinal shortening was observed on the LV free wall than on the septum, which corresponds to the “tilting” deformation mode described in Remme et al. [158]. However, compared to the image data, this mode was exaggerated in our simulation. For the RV, we similarly observed physiological mechanisms of contraction described by Tokodi et al. [139], including AV plane displacement and free wall radial contraction. Tokodi et al. also describe septal bulging into the RV during systole as an RV contraction mechanism. In our simulation, we did not observe septal bulging, although it was also not visible in the CTA image data. The prevalence and functional significance of this mechanism deserve further investigation.

At RR40% (end-systole), the simulated deformation exhibits the greatest discrepancy against the image-based data. While LV wall thickening was matched very well ((Figure 14B), we observed excessive AV plane displacement (Figure 14C), most notably along the free walls, but also in the septum (Figure 14A). Strocchi et al. [42] likewise reported excessive free wall longitudinal shortening at end-systole relative to image data, although the effect was less pronounced than in our study. Pfaller et al. [27] found that endocardial and epicardial fiber angles strongly influenced AV plane displacement, with steeper angles producing greater systolic displacement of the AV plane. This occurs because steeper angles generate a larger net contractile force in the longitudinal direction, pulling the AV plane toward the apex. In our preliminary tests, fiber angles of *±*45^°^ substantially reduced systolic AV plane displacement (data not shown), but we ultimately used the more standard *±*60^°^. We also observed discrepancies at the epicardial apex: in the image data, the apical region moved slightly upward during systole, whereas in our simulations, the RV apex shifted downward and the LV apex remained nearly stationary. Upward systolic apical motion may be reproduced by prescribing steeper fiber angles [27] or incorporating cross-fiber stresses in the sheet-normal direction [55], though these modifications were not explored in the present study.

From RR40% to RR70% (early ventricular filling), we noted the reversal of these deformations. Considering the upward motion of the AV plane during this phase, we discuss two possible mechanisms. The first is the rebound of the compressed passive tissue following the release of active stress. The compression of the passive tissue is evident in Figure 15A, where at RR40%, the majority of the BiV model experiences negative fiber strain. Additionally, from Figure 15B, it is evident that at RR40%, some parts of the LV endocardial region under compressive stress, as indicated by negative values of *σ*_1_. The second is tension from the valve plane Robin BC, which in our BiV model surrogates the mechanical influence of the atria. As seen in Figure 11, the resting position of the AV plane is approximately the same as in the RR70% configuration. Thus, because the RR40% AV plane is significantly lower than in the RR70% state, the valve plane Robin BC is “under tension”, to analogize the Robin BC with a simple spring. This tension tends to pull the AV plane back up to the resting position once the active stress is released. Both mechanisms are related to the concept of ventricular suction, in which elastic recoil (by either of the mechanisms described here) creates a suction effect, actively drawing blood into the ventricles [159]. A criterion for ventricular suction advocated in Zhang et al. [159] is *dP/dV <* 0 during early diastolic filling. Observing ventricular PV loops in Figure 13B (early diastolic phase in blue), we indeed found *dP/dV <* 0 for the LV, but not for the RV. Future work should investigate this difference, as well as alternative definitions of ventricular suction[160].

In our preliminary tests, we found that accounting for the mechanical support of the pericardium and surrounding thoracic anatomy (Section 2.3.4) was critical in reproducing the observed deformation patterns, consistent with previous studies using both uniform [27, 161] and spatially varying [42, 89] epicardial BCs. Pericardial support favors AV plane motion over radial motion, although in our model, the reduced stiffness towards the base permits some radial contraction during systole and expansion during diastole, as observed in the image data. Increased stiffness near the apex helps keep it relatively stationary throughout the cardiac cycle. On the RV side, reduced stiffness was necessary to allow the moderate free wall motion seen in the images (Figure 4 and Figure 14A); however, even with this adjustment, the model underestimated systolic radial contraction. Lower RV stiffness also allowed apex displacement, although in the simulation, this motion occurred in the opposite direction to that observed in the image data.

#### 4.2.3 Strain and stress

We found strong transmural gradients in fiber strain *E*_ff_ at all cardiac phases (Figure 15A). In contrast, experimental studies have reported relatively uniform *E*_ff_ across the myocardial wall, at both end-diastole and end-systole, with systolic fiber shortening typically ranging from 10–20% [162, 163]. However, these studies computed fiber strain relative to the *end-diastolic configuration* (*E*_ff,ED_), whereas *E*_ff_ is computed relative to the *reference configuration*. To facilitate direct comparison with these studies, we recomputed fiber strain relative to the end-diastole configuration (Appendix E, Figure E.1A). Interestingly, *E*_ff,ED_ exhibited smaller transmural variations than *E*_ff_ (Figure E.1B). In the mid-cavity free wall region of the LV (AHA segments 10, 11, 12, and 7), a common focus of experimental studies, both *E*_ff_ and *E*_ff,ED_ peaked at the same mid-wall location at RR40%, but the range of *E*_ff,ED_ (0.102) was smaller than that of *E*_ff_ (0.164). The mean *E*_ff,ED_ was −0.193, comparable to experimentally measured values [162, 163], though on the high side.

High 1^st^ principal stresses were observed in the valve ring region Ω_ring_, particularly in the LV and across all cardiac phases (Figure 15), indicating that the stiff isotropic material in this region was under tension and played an important role in preventing excessive valve ring expansion. This finding supports our design choice to include this stiff region, which represents the collagen-dense tissue of the valve annuli [107], in the model.

### 4.3 Effect of anatomical model and boundary conditions

**t-BiV** We demonstrated that truncating the biventricle model results significantly reduced LV stroke volume as well as LV peak systolic pressure (BiV vs. t-BiV(same base) in Figure 16, Section 3.3). These differences stem from the t-BiV geometry excluding the portion of the ventricle above the basal truncation plane, which reduces the overall chamber volume. The smaller chamber reduces both end-diastolic and end-systolic volumes, and also results in a smaller stroke volume, generating a lower systolic pressure upon ejection into the systemic circulation. Similar trends were observed in the RV PV diagram, though the drop in RV systolic pressure was smaller. We also observed excessive RV free wall motion in t-BiV(same base), suggesting that the tissue above the basal plane likely plays an important role in constraining RV free wall motion.

Increasing the basal stiffness in t-BiV(stiffer base) reduced this excessive deformation, but also caused unphysiological bending at the basal plane (Figure 17A). This arises because ventricular pressurization pushes the LV and RV free walls outward, while the stiff Robin BC at the base resists outward displacement. The stiffer base also further reduced LV stroke volume and LV peak systolic pressure (Section 3.4). Interestingly, these findings are in contradiction to the work by Peirlinck et al. [50], who compared different kinematic BCs applied on the basal surface of a truncated BiV model. They found that stricter constraints on basal plane motion *increased* LV stroke volumes and peak systolic pressures. The discrepancy may be due to different choices in BCs; in their work, all the basal BCs they investigated prevented longitudinal motion of the base, and they did not apply any epicardial BC. This combination leads to a fundamentally different mode of deformation in which the base is fixed and the apex moves, which is usually not observed in patient images, and may account for the contradictory conclusion reached in the present work.

Comparisons between t-BiV(higher stress) and t-BiV(stiffer base) showed nearly identical deformation patterns at the imaged time points (Figure 17A). However, t-BiV(higher stress) produced higher peak systolic pressures and smaller minimum volumes in both LV and RV (Figure 17B). The slightly smaller minimum volume is due to subtle differences in deformation at RR40%, while the higher systolic pressure is explained by a faster increase in active stress. Although not apparent in the snapshots, this faster contraction increases systolic flow rate out of the ventricles, thereby boosting peak pressure. This increase in LV pressure, however, is not enough to match t-BiV(same base), much less BiV.

Overall, these findings underscore the limitations and challenges associated with using models truncated at the basal plane. Despite reasonable variations to BCs and contractility, all t-BiV models significantly underpredicted LV peak systolic pressure and exhibited unphysiological deformations. This is not to say that truncated models cannot produce accurate results. With different parameterizations of BCs, material properties, and the circulatory LPN model, the t-BiV model would likely be able to reproduce physiological deformations and match hemodynamic targets. Indeed, several studies have successfully personalized truncated geometries using modeling assumptions and parameter tuning approaches similar to those employed in the present work [33, 49, 57]. However, we showed that parameter sets that perform well for a full BiV model produce very different results when used in a truncated model. Furthermore, to the extent such parameters are intended to represent real physical quantities, truncated models risk introducing systematic errors in their estimated values. These issues do not make truncated models inherently unsuitable, but they do require that their limitations be clearly understood and accounted for in both research and clinical applications.

**LV** The nearly identical LV PV loops between LV(all Robin) and BiV (Figure 16B) is generally consistent with the findings of Palit et al. [78], who compared LV diastolic filling in an LV-only and a BiV model. They found that at a pressure of 10 mmHg, LV volume was 3-5% greater in the BiV model versus the LV-only model. However, we caution that their model used fixed base and free epicardium BCs, which complicates the comparison of these results. Also, their LV-only model is more similar to our LV(free septum) model, for which the LVEDV is only marginally smaller (*<* 1%) than that of the BiV case.

Some differences were observed in local myocardial deformation, particularly near the apex. One key difference was an apparent bulging of the lower part of the septum into the LV during diastole (Figure 16A). This behavior arises from a combination of the septal position in the estimated reference configuration (Figure 11A) and the Robin BC applied to the septal portion of the LV epicardium Γ_epi,septum_ (Section 2.5). In the reference configuration (Figure 11A), the septum is displaced into the LV cavity relative to its in vivo position. In the full BiV model, LV pressurization acts to push the septum uniformly back toward the RV cavity, restoring a more physiological in vivo position. In contrast, in LV(all Robin), the Robin BC applied on Γ_epi,septum_ restricts motion in the lower septum where the stiffness is greatest, while allowing greater displacement in the upper septum. As a result, pressurization pushes the upper septum toward the RV cavity but leaves the lower septum relatively stationary, producing the observed bulging of the lower septum into the LV. This constraint also generally reduces septal motion throughout the cardiac cycle. Given the clinical relevance of septal motion in both health and disease [164, 165, 166], restricting it in computational models with a Robin BC may be inappropriate.

The LV-only models also exhibited downward motion of the apex during systole (Figure 16A and Figure 18A) in contrast to the stationary apex in the BiV model. Together with the slightly greater basal displacement, this points to an overall bulk downward displacement of the LV. This motion may be explained by the net downward force on the LV due to blood pressure, which arises because the LV endocardial surface is not closed. In the BiV model, we observed much smaller downward motion of the LV, suggesting that RV tissue support helps resist the LV’s downward motion.

We tested three different boundary conditions on the LV septal epicardium. Overall scalar outputs were remarkably insensitive to these changes, in agreement with previous whole-heart simulations of Pfaller et al. [27], who also found only minor differences in global hemodynamics when varying epicardial boundary conditions on LV. However, local myocardial deformation patterns did vary. In LV(loaded septum), the action of RV pressure applied to the septal epicardium, combined with spatially varying epicardial BCs, pushed the LV toward the anatomical left during systole, enhancing the tilting mode described by Remme et al. [158]. LV(free septum) agreed best with image data, exhibiting no tilting. In LV(all Robin), the Robin BC moderated this behavior, somewhat resisting the tilting moment and producing an intermediate tilt between those of LV(loaded septum) and LV(free septum). In BiV, the presence of the RV also reduced tilting. These results suggest that while RV pressure tends to push the LV leftward anatomically, the competing tissue support from RV tends to pull the LV back toward the septum. For this reason, RV pressure should not be applied unless the RV’s tethering effect is also modeled, either by including the RV explicitly, as in BiV, or by using a Robin BC on the septal epicardium, as in LV(all Robin), although the latter may inappropriately constrain septal motion. Without such tethering, RV pressure should be omitted, as in LV(free septum). These findings complement prior computational work on RV–LV mechanical interactions, such as Hadjicharalambous et al. [167], who modeled RV traction on the LV septum during passive filling, and Asner et al. [49], who showed that RV epicardial boundary conditions can constrain LV/RV junction deformation to a more anatomically realistic position, thereby reducing model error.

### 4.4 Limitations

The primary discrepancy between our personalized BiV model and the clinical data lies in the suboptimal agreement of the simulated tissue deformation with the collected CTA image data. Mainly, the model overestimates systolic AV plane displacement and predicts unphysiological systolic apex motion (Figure 14). Such local mismatches in myocardial motion are common in many personalized models [27, 41, 42]. Nonetheless, it is worth noting that some recent personalized whole-heart models, such as that of Strocchi et al. [32], have reported close agreement with image-derived AV plane displacement, even though other aspects of myocardial motion were not explicitly validated.

A contributing factor is the simplified treatment of boundary conditions, which play an important role in determining myocardial deformation. In this work, we manually tuned the parameters of the spatially varying epicardial BC and the valve ring BC to constrain the tissue from unphysiological deformation, while permitting enough deformation to roughly match the image data and clinical pressure and volume targets. A comprehensive BC tuning procedure could improve deformation agreement, but we leave this for future work. Prior studies have already investigated the sensitivity of simulation results to BCs [27, 29, 47, 50], but more work needs to be done. Although a spatially varying epicardial BC was used in this work, this is still a simplification of the complex pattern of tissue support on the epicardial surface [27]. While a more complex spatial variation could improve the deformation agreement, it would also introduce additional parameters that must be personalized, which in turn may require more comprehensive clinical measurements and more advanced parameter tuning techniques. Future work should also investigate the valve ring BC. Preliminary tests suggest a tradeoff: increasing the valve ring stiffness may reduce excessive systolic shortening but also inhibit the necessary upward AV plane movement during diastole. Modeling the effect of the atria could address this issue, as atrial contraction has been shown to influence ventricular diastolic mechanics [46, 74]. One approach to approximate this effect without modeling the atria explicitly would be to apply a time-varying Robin stiffness at the base. Future work may also consider the energy-consistent boundary condition proposed by Regazzoni et al. [86] and Piersanti et al. [55] to address this issue.

The discrepancy in the simulated deformation may also be due to our use of rule-based fiber orientations, which are not patient-specific. Indeed, fiber direction has a strong influence on deformation patterns [168, 169, 170]. Pfaller et al. [27] found that modest changes in rule-based method parameters can lead to substantially different deformation patterns; specifically, steeper endocardial and epicardial fiber angles increase AV plane displacement and reduce radial contraction of the free walls. Ogiermann et al. [171] discovered that the sheetlet orientation **s** has a strong influence on many kinematic measures of left ventricular function. The rule-based fiber orientations also likely contribute to the spatially nonuniform fiber strains observed in our simulations (Figure 15), which persist even when recomputing fiber strains relative to the end-diastolic configuration (Figure E.1). Transmural gradients were also observed in Palit et al. [78] using a rule-based method over a range of fiber angles. These findings diverge from the largely uniform transmural fiber strain reported experimentally [163]. Future work could personalize fiber angles in a rule-based method, optimize fiber angles to homogenize active tension generation and fiber shortening [172], employ a nested toroidal structure for myofibers [173], or account for truly personalized mesostructure by incorporating *in vivo* diffusion-tensor MRI (DTMRI) [174, 175, 176, 177, 178]. *In vivo* DTMRI, however, still faces several challenges, such as long acquisition times, limited spatial resolution, and the obstacles associated with compensating for bulk cardiac motion [179, 180].

Our model would benefit from the inclusion of additional clinical data in general. In particular, we relied on literature-derived values for pulmonary, atrial, and peripheral venous pressures (Table 3) to constrain the model, but ideally, these parameters would be measured directly in each patient. Catheterization could provide such data [51], although it is minimally invasive and not routinely performed. Alternatively, Doppler echocardiography can be used to noninvasively estimate pressure gradients [181, 182], which could in turn be used to approximate absolute pressures. For the passive mechanics personalization (Section 2.4.2), we used pressure waveforms from the tuned 0D surrogate model. While this approach was effective, greater accuracy could be achieved with *in vivo* ventricular pressure measurements synchronized to the imaging data [32]. Additional modalities could further improve model personalization. For instance, tagged MRI can capture ventricular twist [183, 184], a feature not resolved by conventional CTA or MRI [27].

Given the size and complexity of the optimization problem, the loss landscape is likely to contain numerous local minima. While we did not perform formal uncertainty quantification (UQ) [30, 132] in this study, exploring the sensitivity of the calibrated parameters to different initial conditions or optimizer seeds could provide valuable insight into the variability of the resulting parameters, as well as the predicted PV loops and deformation patterns [143]. In both the 0D parameter estimation and iFEA cases, we employed evolutionary algorithms, which are generally effective at locating global optima [185]. Nevertheless, parameter sensitivity and identifiability remain notable limitations, and a systematic UQ analysis would be an important extension for future work. Future work should also explore alternative approaches to estimating the reference configuration, such as the method proposed by Barnafi et al. [186], which addresses identifiability of the inverse mechanics problem in complex cardiac modeling contexts.

Finally, we aim to extend this analysis of the effect of anatomical model to heart models that include not only the atria, but also other less studied cardiac structures like the roots of the great vessels [42, 45, 74], the trabeculae and papillary muscles [187, 188, 189], and epicardial adipose tissue [27, 45, 190]. While several studies have begun to explore this direction [27, 46, 74, 75, 76], further investigation is needed to fully understand the importance of these structures in the context of patient-specific modeling.

## 5 Conclusion

Personalized computational heart models are powerful tools for investigating cardiac function with direct clinical relevance. Yet, it remains unclear how the choice of anatomical representation (LV vs. BiV, truncated vs. not) affects simulation outcomes and the conclusions drawn from them. To anchor our study in a physiologically meaningful context, we first developed a workflow to identify personalized parameters for a patient-specific BiV model. This model then served as a reference for evaluating the impact of anatomical simplifications under plausible variations in boundary conditions and contractile strength. Our results indicate that while LV-only models can reasonably approximate the BiV model in terms of global hemodynamics and regional myocardial mechanics, truncation at the basal plane leads to substantial deviations in both. These findings provide guidance for cardiac modelers in choosing anatomical representations that balance computational efficiency with physiological fidelity.

## Acknowledgements

AB and AM would like to acknowledge financial support from National Institutes of Health (R01HL129727, R01HL173845) and and the National Science Foundation (1663671, 2310909). VV would like to acknowledge financial support from the American Heart Association’s Second Century Early Faculty Independence Award (24SCE-FIA1260268) and the National Science Foundation’s CAREER Award (2443726).

## APPENDIX A LPN equations

We use the 0D closed-loop circulation model proposed in [87], which is governed by the following ODE system. Here, we provide the equations for the full LPN model, which also uses TVEs for the LV and RV. In Appendix B, we explain how these equations are modified when coupling the LPN to the 3D ventricular model.

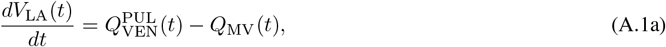

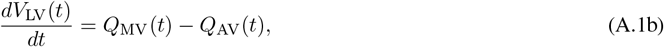

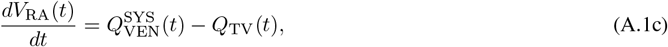

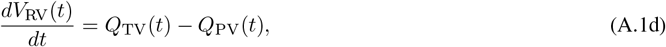

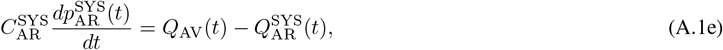

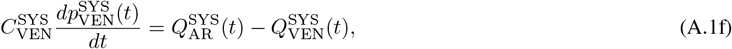

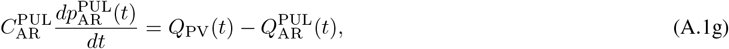

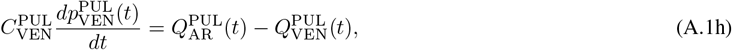

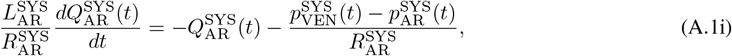

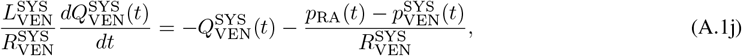

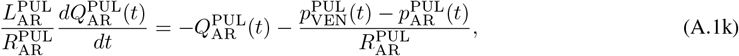

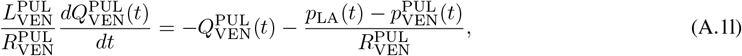

with initial conditions 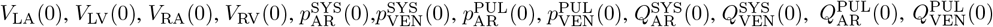, and *V* denote flowrate, pressure, and volume, respectively, and *R, C*, and *L* denote resistance, capacitance, and inductance, respectively. The subscripts and superscript indicate location in the LPN: LA = left atrium, LV = left ventricle, RA = right atrium, RV = right ventricle, PUL = pulmonary, SYS = systemic, AR = arterial, VEN = venous.

The valvular flowrates are given by

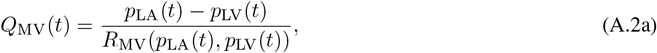

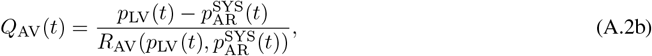

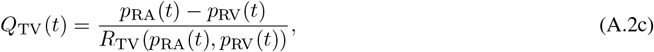

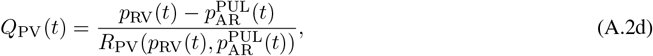

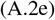

Where

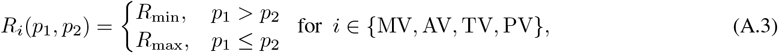

where *p*_1_ denotes the pressure upstream of the valve, and *p*_2_ the pressure downstream.

The cardiac chambers are modeled as TVE elements, according to the following equation

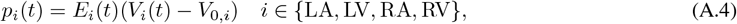

where *p*_*i*_ is the pressure of chamber *C, V*_*i*_ is the volume, *E*_*i*_ is the TVE, and *V*_0,*i*_ is the unloaded or rest volume. *E*_*i*_ is given by

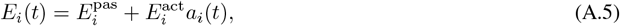

where *a*_*i*_ is a double Hill activation function (Eq. (16)) with different parameters *τ*_1,*i*_, *τ*_2,*i*_, *m*_1,*i*_, *m*_2,*i*_, *t*_*C,i*_ for each chamber *i* ∈ {LA, RA, LV, RV}. These parameters can be found in Table D.1.

## Appendix B Revised LPN equations for coupled 3D-0D model

Our 3D-0D coupling relies on passing flowrate from the 3D to the 0D domain, and passing pressure from the 0D to the 3D domain (Figure 5A). Because of this, when coupling the circulation model to a 3D model of the left and right ventricles, the circulation model must be modified to reflect flow rate boundary conditions into the LV and RV nodes. This results in the following modifications to the LPN equations presented in Appendix A. *V*_LV_ and *V*_RV_ are replaced by *p*_LV_ and *p*_RV_ as state variables of the circulation system. Consequently, Eqs. (A.1b) and (A.1d) are replaced by

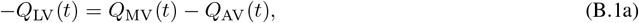

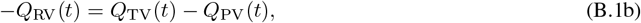

where *Q*_LV_ and *Q*_RV_ are the flowrates passed from the 3D biventricular model. Recalling Eq. (A.2) for the valvular flowrates, we obtain implicit algebraic equations for *p*_LV_ and *p*_RV_,

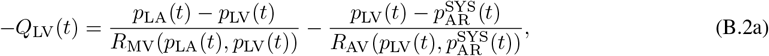

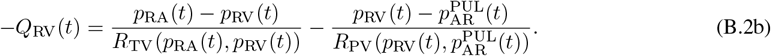

These equations replace the original equations for the LV and RV pressure, given by the TVE model (Eq. (A.4)). Eqs. (B.2a) and (B.2b) can be manipulated into a quasi-explicit form,

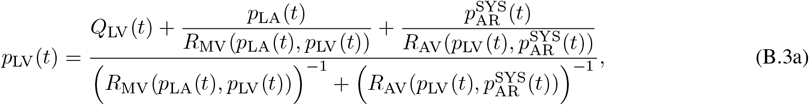

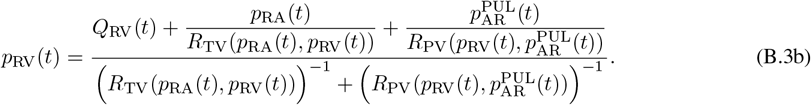

When coupling to 3D, the circulation model is integrated from timestep *n* to *n* + 1 every Newton iteration of the 3D solver, and Eqs. (B.3a) and (B.3b) are implicit equations for 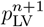 and 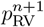. To avoid numerical issues associated with valves opening and closing between Newton iterations, and to make Eqs. (B.3a) and (B.3b) explicit and thus easier to solve, we use 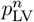 and 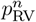 to evaluate valve resistances [85].

## Appendix C Calculating wall thickness and valve plane displacement

Here, we describe how LV and RV wall thicknesses, as well as MV and TV valve plane displacements, are computed (Figure 14B–C). For clarity, we present the wall thickness calculation for the LV and the valve plane displacement for the MV; the same procedures are applied to the RV wall thickness and TV plane displacement, respectively.

LV wall thickness is computed by identifying, for each node on the epicardial surface Γ_epi_, the nearest node on the LV endocardial surface Γ_endo,LV_ (Figure 3E). The nearest-neighbor search uses the `query()` method of the `cKDTree` class in `scipy.spatial`, which provides efficient distance computation. This produces a distribution of minimum distances representing the separation between the LV endocardium and the entire epicardial surface. Because Γ_epi_ includes both LV and RV regions, the resulting distribution is bimodal: one mode corresponds to the LV free wall and the other to the more distant RV portion of the epicardial surface. To separate these regions, a Gaussian mixture model (GMM) with two components is fitted to the distance data using `sklearn.mixture.GaussianMixture`. The smaller of the two Gaussian means is taken as the LV wall thickness, Δ_LV_.

MV plane displacement is computed using a mitral valve ring surface Γ_MV_, identified as a subset of the entire valve ring surface Γ_valves_ (Figure 3E). A plane *P*_MV_ is fitted to the points of Γ_MV_ using PyVista’s `fit_plane_to_points()` function. The plane is sized to the extent of the points so that its center, **c**_MV_, approximates the centroid of Γ_MV_. We choose the image-based RR70% configuration (Figures 3 and 4) as the zero-displacement state. At this state, a reference plane center, **c**_MV,ref_, and a reference plane normal, **n**_MV,ref_, are computed. The MV plane displacement at any time *t, d*_MV_, is then defined as the projection of the displacement of the plane center, **c**_MV_, onto the reference normal,

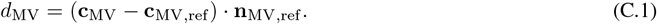

## Appendix D ECG-based double Hill cardiac activation

In this work, the double Hill activation model Eq. (16) is used to prescribe ventricular and atrial contraction. In the 3D-0D model, it is used for the 3D ventricular active stress (Eq. (15)) and the 0D atrial TVEs (Section 2.3.3). In the full 0D model, it is used for both ventricular and atrial TVEs (Eq. (A.5)).

The double Hill parameters used in this study for atria and ventricles can be found in Table D.1. Parameters are identical for both atria (LA and RA) and both ventricles (LV and RV). These parameters yield the atrial (*a*_LA/RA_(*t*)) and ventricular (*a*_LV/RV_(*t*)) activation curves shown in Figure D.1. Values from [113] are used for *τ*_1,*i*_, *τ*_2,*i*_, *m*_1,*i*_, and *m*_2,*i*_, for *i* ∈ {LA/RA, LV/RV}. In contrast, the contraction times *t*_*C,i*_ are calculated from the collected ECG data, as described below.

**Table D.1.** ECG and double Hill activation model parameters.

| Parameter | Symbol | Value | Units |
| --- | --- | --- | --- |
| **ECG and timing parameters** |  |  |  |
| Heart rate | BPM | 87.0 | beats/min |
| PR interval | PR | 0.182 | s |
| Electromechanical delay | EMD | 0.025 | s |
| Cardiac cycle period ($\frac{60}{\text{BPM}}$) | $T_{HB}$ | 0.690 | s |
| **Double Hill [113]** |  |  |  |
| Atrial contraction rate constant | $m_{1,LA/RA}$ | 1.32 | - |
| Atrial relaxation rate constant | $m_{2,LA/RA}$ | 13.1 | - |
| Atrial contraction time constant | $\tau_{1,LA/RA}$ | $0.110 T_{HB}$ | s |
| Atrial relaxation time constant | $\tau_{2,LA/RA}$ | $0.180 T_{HB}$ | s |
| Atrial contraction start time | $t_{C,LA/RA}$ | EMD | s |
| Ventricular contraction rate constant | $m_{1,LV/RV}$ | 1.32 | - |
| Ventricular relaxation rate constant | $m_{2,LV/RV}$ | 27.4 | - |
| Ventricular contraction time constant | $\tau_{1,LV/RV}$ | $0.269 T_{HB}$ | s |
| Ventricular relaxation time constant | $\tau_{2,LV/RV}$ | $0.452 T_{HB}$ | s |
| Ventricular contraction start time | $t_{C,LV/RV}$ | PR + EMD | s |

**Figure D.1.**
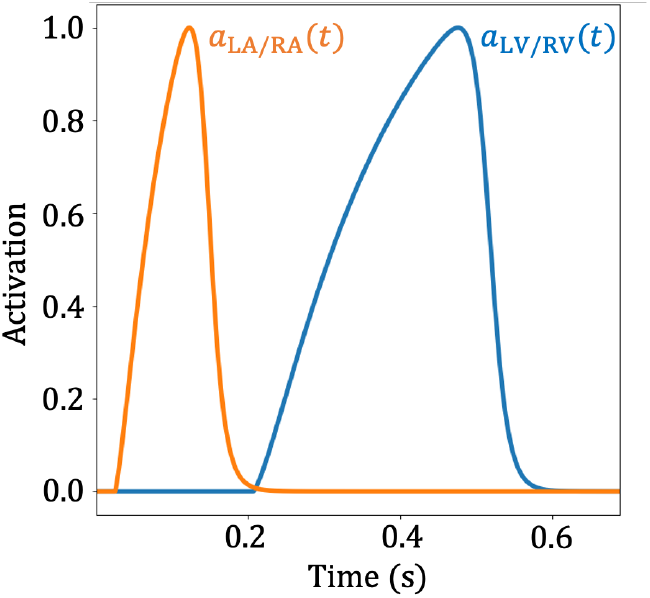
Double hill activation curves over one cardiac cycle for the atria (*a*_LA/RA_) and the ventricles (*a*_LV/RV_).

We assume that for any chamber, mechanical contraction begins at electrical depolarization, plus an electromechanical delay (EMD), which we assume to be ~25 ms based on literature [191]. EMD is generally spatially heterogeneous, but for simplicity we use a constant EMD for all chambers. It is known that the features of the ECG reflect electrical events at various locations in the heart. In particular, the P wave is associated with atrial depolarization, and the QRS complex with ventricular depolarization [192]. Using these principles, we write the following equations for the start times of atrial (*t*_*C*,LA/RA_) and ventricular (*t*_*C*,LV,RV_) mechanical contraction. Let the start of the P wave be *t* = 0 ms.

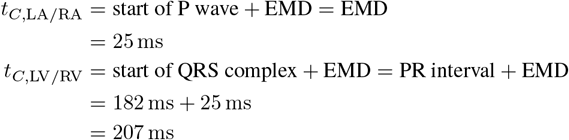

These equations mean that for both full 0D and 3D-0D models, the beginning of the simulation *t* = 0 corresponds to the start of the P wave. This is used to synchronize the time-dependent image data and simulation results.

## Appendix E Fiber strain relative to end-diastolic state

Experimental results for *in vivo* fiber strain typically measure strain at end-systole relative to end-diastole. To provide a fair comparison, we recompute fiber strain relative to the end-diastolic state, rather than to the reference configuration. We use the following equation:

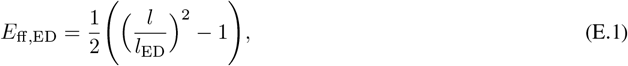

where 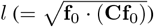 is the length of a fiber in the deformed configuration and *l*_ED_ is the length of the fiber in the end-diastolic configuration.

*E*_ff,ED_ exhibits a smaller transmural variation at RR40% than *E*_ff_ (Figure 15). To quantify this, in Figure E.1B, we plot both *E*_ff,ED_ and *E*_ff_ against the transmural Laplace field Φ_epi_. The data is taken from the LV mid-cavity free wall region, composed of LV segments 10, 11, 12, and 7. While *E*_ff,ED_ still displays transmural variation at RR40%, the magnitude of variation is reduced. For *E*_ff,ED_, the mean value is −0.193 and the range (max-min) is 0.102, while for *E*_ff_, the mean value is −0.094 and the range is 0.164.

**Figure E.1.**
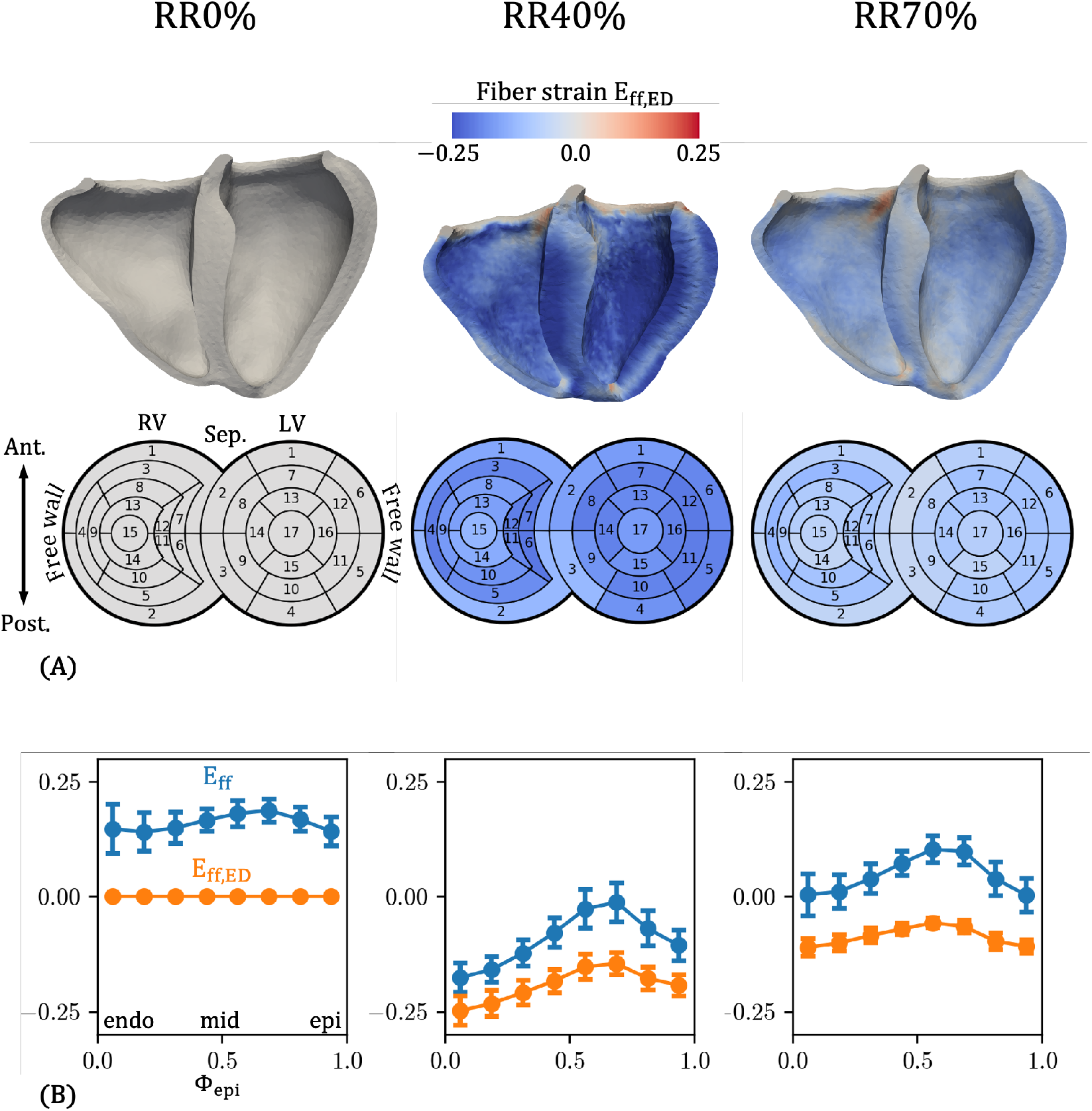
**A**: Fiber strain relative to the end-diastolic state (*E*_ff,ED_ for the personalized 3D-0D BiV model, at three phases during the cardiac cycle. Strain is also plotted on BiV bullseye plots, using a standard 17 segment model for the LV [138] and a 15 segment model for the RV from [139]. **B**: Transmural variation of *E*_ff_ and *E*_ff,ED_ for the LV mid-cavity free wall region, composed of LV segments 10, 11, 12, and 7.

## Footnotes

1 The coordinate system {**f, n, s**} corresponds to {*F, S, T*} in Bayer et al. [93]. We adopt the notation {**f, n, s**} because it is now standard in the current literature. Moreover, it is used in the Holzapfel–Ogden material model for myocardium [104], which we employ in the present study.

2 There is some confusion in the literature regarding the definition of the *β* angles, which we address in our recent review [105].

3 Φ_ab_ also parameterizes the BiV long axis, but Ψ_long_ is convenient in practice due to its uniform variation over the domain (Figure 6A). An alternative would be to smooth Φ_ab_ by mapping it to a geodesic from apex to base, as done in the universal ventricular coordinates framework [109].

4 Using displacements **u** and velocities **v** in this boundary condition violates the principle of objectivity (see “[Simulating overdamped dynamics through a surface drag force](https://forums.febio.org/forum/febio-forums/users-forum/25978-simulating-overdamped-dynamics-through-a-surface-drag-force#post26469)” on the FEBio Forum). This limitation can be avoided by explicitly representing the pericardium as a solid body, as done by Fritz et al. [41]. Nevertheless, the boundary condition proposed by Pfaller et al. [27] remains very practical and popular, and the implications of its nonobjectivity merit further investigation.

5 The spatially varying stiffnesses and myofiber directions are mapped from Ω_RR70_ to Ω_*X*_ using the same procedure as described in Section 2.4.2.

## References

[1] Roel Meiburg, Jesse H J Rijks, Ahmed S Beela, Edoardo Bressi, Domenico Grieco, Tammo Delhaas, Justin GLM Luermans, Frits W Prinzen, Kevin Vernooy, and Joost Lumens. Comparison of novel ventricular pacing strategies using an electro-mechanical simulation platform. EP Europace, 25(6):euad144, June 2023. ISSN 1099-5129. doi:10.1093/europace/euad144. URL https://doi.org/10.1093/europace/euad144.

[2] Emilia Capuano, Francesco Regazzoni, Massimiliano Maines, Silvia Fornara, Vanessa Locatelli, Domenico Catanzariti, Simone Stella, Fabio Nobile, Maurizio Del Greco, and Christian Vergara. Personalized computational electro-mechanics simulations to optimize cardiac resynchronization therapy. Biomechanics and Modeling in Mechanobiology, August 2024. ISSN 1617-7940. doi:10.1007/s10237-024-01878-8. URL https://doi.org/10.1007/s10237-024-01878-8.

[3] Marina Strocchi, Jack W Samways, Akriti Naraen, Nadine Ali, Matthew J Shun-Shin, Karli Gillette, Christopher Aldo Rinaldi, Ahran D Arnold, Gernot Plank, Edward J Vigmond, Zachary I Whinnett, and Steven A Niederer. An in silico guide for ventriculo-ventricular delay programming for left bundle branch-optimized cardiac resynchronization therapy. EP Europace, 27(5):euaf089, May 2025. ISSN 1099-5129. doi:10.1093/europace/euaf089. URL https://doi.org/10.1093/europace/euaf089.

[4] Adityo Prakosa, Hermenegild J. Arevalo, Dongdong Deng, Patrick M. Boyle, Plamen P. Nikolov, Hiroshi Ashikaga, Joshua J. E. Blauer, Elyar Ghafoori, Carolyn J. Park, Robert C. Blake, Frederick T. Han, Rob S. MacLeod, Henry R. Halperin, David J. Callans, Ravi Ranjan, Jonathan Chrispin, Saman Nazarian, and Natalia A. Trayanova. Personalized virtual-heart technology for guiding the ablation of infarct-related ventricular tachycardia. Nature Biomedical Engineering, 2(10):732–740, October 2018. ISSN 2157-846X. doi:10.1038/s41551-018-0282-2. URL https://www.nature.com/articles/s41551-018-0282-2. Publisher: Nature Publishing Group.

[5] Patrick M. Boyle, Tarek Zghaib, Sohail Zahid, Rheeda L. Ali, Dongdong Deng, William H. Franceschi, Joe B. Hakim, Michael J. Murphy, Adityo Prakosa, Stefan L. Zimmerman, Hiroshi Ashikaga, Joseph E. Marine, Aravindan Kolandaivelu, Saman Nazarian, David D. Spragg, Hugh Calkins, and Natalia A. Trayanova. Computationally guided personalized targeted ablation of persistent atrial fibrillation. Nature Biomedical Engineering, 3(11):870–879, November 2019. ISSN 2157-846X. doi:10.1038/s41551-019-0437-9. URL https://www.nature.com/articles/s41551-019-0437-9. Publisher: Nature Publishing Group.

[6] Lik Chuan Lee, Liang Ge, Zhihong Zhang, Matthew Pease, Serjan D. Nikolic, Rakesh Mishra, Mark B. Ratcliffe, and Julius M. Guccione. Patient-specific finite element modeling of the Cardiokinetix Parachute® device: effects on left ventricular wall stress and function. Medical & Biological Engineering & Computing, 52(6): 557–566, June 2014. ISSN 1741-0444. doi:10.1007/s11517-014-1159-5. URL https://doi.org/10.1007/s11517-014-1159-5.

[7] Ethan Kung, Masoud Farahmand, and Akash Gupta. A Hybrid Experimental-Computational Modeling Framework for Cardiovascular Device Testing. Journal of Biomechanical Engineering, 141(051012), March 2019. ISSN 0148-0731. doi:10.1115/1.4042665. URL https://doi.org/10.1115/1.4042665.

[8] Tain-Yen Hsia, Timothy Conover, and Richard Figliola. Computational Modeling to Support Surgical Decision Making in Single Ventricle Physiology. Seminars in Thoracic and Cardiovascular Surgery: Pediatric Cardiac Surgery Annual, 23:2–10, January 2020. ISSN 1092-9126. doi:10.1053/j.pcsu.2020.01.001. URL https://www.sciencedirect.com/science/article/pii/S1092912620300016.

[9] Yue-Hin Loke, Francesco Capuano, Elias Balaras, and Laura J. Olivieri. Computational Modeling of Right Ventricular Motion and Intracardiac Flow in Repaired Tetralogy of Fallot. Cardiovascular Engineering and Technology, 13(1):41–54, February 2022. ISSN 1869-4098. doi:10.1007/s13239-021-00558-3. URL https://link.springer.com/article/10.1007/s13239-021-00558-3. Company: Springer Distributor: Springer Institution: Springer Label: Springer Publisher: Springer International Publishing.

[10] Francisco Sahli-Costabal, Kinya Seo, Euan Ashley, and Ellen Kuhl. Classifying Drugs by their Arrhythmogenic Risk Using Machine Learning. Biophysical Journal, 118(5):1165–1176, March 2020. ISSN 0006-3495. doi:10.1016/j.bpj.2020.01.012. URL https://www.sciencedirect.com/science/article/pii/S0006349520300382.

[11] M. Peirlinck, J. Yao, F. Sahli Costabal, and E. Kuhl. How drugs modulate the performance of the human heart. Computational Mechanics, 69(6):1397–1411, June 2022. ISSN 1432-0924. doi:10.1007/s00466-022-02146-1. URL https://doi.org/10.1007/s00466-022-02146-1.

[12] Ryan P O’Hara, Edem Binka, Adityo Prakosa, Stefan L Zimmerman, Mark J Cartoski, M Roselle Abraham, Dai-Yin Lu, Patrick M Boyle, and Natalia A Trayanova. Personalized computational heart models with T1-mapped fibrotic remodeling predict sudden death risk in patients with hypertrophic cardiomyopathy. eLife, 11:e73325, January 2022. ISSN 2050-084X. doi:10.7554/eLife.73325. URL https://doi.org/10.7554/eLife.73325. Publisher: eLife Sciences Publications, Ltd.

[13] Ekaterina Kovacheva, Tobias Gerach, Steffen Schuler, Marco Ochs, Olaf Dössel, and Axel Loewe. Causes of altered ventricular mechanics in hypertrophic cardiomyopathy: an in-silico study. BioMedical Engineering OnLine, 20(1):69, July 2021. ISSN 1475-925X. doi:10.1186/s12938-021-00900-9. URL https://doi.org/10.1186/s12938-021-00900-9.

[14] Alejandro Gonzalo, Christoph M. Augustin, Savannah F. Bifulco, Åshild Telle, Yaacoub Chahine, Ahmad Kassar, Manuel Guerrero-Hurtado, Eduardo Durán, Pablo Martínez-Legazpi, Oscar Flores, Javier Bermejo, Gernot Plank, Nazem Akoum, Patrick M. Boyle, and Juan C. del Alamo. Multiphysics simulations reveal haemodynamic impacts of patient-derived fibrosis-related changes in left atrial tissue mechanics. The Journal of Physiology, 602(24):6789–6812, 2024. ISSN 1469-7793. doi:10.1113/JP287011. URL https://onlinelibrary.wiley.com/doi/abs/10.1113/JP287011. _eprint: https://onlinelibrary.wiley.com/doi/pdf/10.1113/JP287011.

[15] Abigail E. Teitgen, Marcus T. Hock, Kimberly J. McCabe, Matthew C. Childers, Gary A. Huber, Bahador Marzban, Daniel A. Beard, J. Andrew McCammon, Michael Regnier, and Andrew D. McCulloch. Multiscale modeling shows how 2’-deoxy-ATP rescues ventricular function in heart failure. Proceedings of the National Academy of Sciences, 121(35):e2322077121, August 2024. doi:10.1073/pnas.2322077121. URL https://www.pnas.org/doi/abs/10.1073/pnas.2322077121. Publisher: Proceedings of the National Academy of Sciences.

[16] Adriana Gaia Cairelli, Alex Gendernalik, Wei Xuan Chan, Phuc Nguyen, Julien Vermot, Juhyun Lee, David Bark, and Choon Hwai Yap. Role of tissue biomechanics in the formation and function of myocardial trabeculae in zebrafish embryos. The Journal of Physiology, 602(4):597–617, February 2024. ISSN 1469-7793. doi:10.1113/JP285490.

[17] Jing Wang, Aaron L. Brown, Seul-Ki Park, Charlie Z. Zheng, Adam Langenbacher, Enbo Zhu, Ryan O’Donnell, Peng Zhao, Jeffrey J. Hsu, Tomohiro Yokota, Jiandong Liu, Jau-Nian Chen, Alison L. Marsden, and Tzung K. Hsiai. Mechanically activated snai1b coordinates the initiation of myocardial delamination for trabeculation. Nature Communications, 16(1):8363, September 2025. ISSN 2041-1723. doi:10.1038/s41467-025-62285-w. URL https://www.nature.com/articles/s41467-025-62285-w. Publisher: Nature Publishing Group.

[18] Johane H. Bracamonte, Sarah K. Saunders, John S. Wilson, Uyen T. Truong, and Joao S. Soares. Patient-Specific Inverse Modeling of In Vivo Cardiovascular Mechanics with Medical Image-Derived Kinematics as Input Data: Concepts, Methods, and Applications. Applied Sciences, 12(8):3954, January 2022. ISSN 2076-3417. doi:10.3390/app12083954. URL https://www.mdpi.com/2076-3417/12/8/3954. Number: 8 Publisher: Multidisciplinary Digital Publishing Institute.

[19] Marina Strocchi, Cristobal Rodero, Caroline H. Roney, Caroline Mendonca Costa, Gernot Plank, Pablo Lamata, and Steven A. Niederer. A Semi-automatic Pipeline for Generation of Large Cohorts of Four-Chamber Heart Meshes. In Michael Regnier and Matthew Childers, editors, Familial Cardiomyopathies: Methods and Protocols, pages 117–127. Springer US, New York, NY, 2024. ISBN 978-1-07-163527-8. doi:10.1007/978-1-0716-3527-8_7. URL https://doi.org/10.1007/978-1-0716-3527-8_7.

[20] Turki Nasser Alnasser, Lojain Abdulaal, Ahmed Maiter, Michael Sharkey, Krit Dwivedi, Mahan Salehi, Pankaj Garg, Andrew James Swift, and Samer Alabed. Advancements in cardiac structures segmentation: a comprehensive systematic review of deep learning in CT imaging. Frontiers in Cardiovascular Medicine, 11, January 2024. ISSN 2297-055X. doi:10.3389/fcvm.2024.1323461. URL https://www.frontiersin.org/journals/cardiovascular-medicine/articles/10.3389/fcvm.2024.1323461/full. Publisher: Frontiers.

[21] Lukasz Romaszko, Agnieszka Borowska, Alan Lazarus, David Dalton, Colin Berry, Xiaoyu Luo, Dirk Husmeier, and Hao Gao. Neural network-based left ventricle geometry prediction from CMR images with application in biomechanics. Artificial Intelligence in Medicine, 119:102140, September 2021. ISSN 0933-3657. doi:10.1016/j.artmed.2021.102140. URL https://www.sciencedirect.com/science/article/pii/S0933365721001330.

[22] Fanwei Kong, Nathan Wilson, and Shawn Shadden. A deep-learning approach for direct whole-heart mesh reconstruction. Medical Image Analysis, 74:102222, December 2021. ISSN 1361-8415. doi:10.1016/j.media.2021.102222. URL https://www.sciencedirect.com/science/article/pii/S136184152100267X.

[23] Fanwei Kong and Shawn C. Shadden. Learning Whole Heart Mesh Generation From Patient Images for Computational Simulations. IEEE Transactions on Medical Imaging, 42(2):533–545, February 2023. ISSN 1558-254X. doi:10.1109/TMI.2022.3219284. URL https://ieeexplore.ieee.org/abstract/document/9936657. Conference Name: IEEE Transactions on Medical Imaging.

[24] Fanwei Kong, Sascha Stocker, Perry S. Choi, Michael Ma, Daniel B. Ennis, and Alison L. Marsden. SDF4CHD: Generative modeling of cardiac anatomies with congenital heart defects. Medical Image Analysis, 97:103293, October 2024. ISSN 1361-8415. doi:10.1016/j.media.2024.103293. URL https://www.sciencedirect.com/science/article/pii/S1361841524002184.

[25] Arjun Narayanan, Fanwei Kong, and Shawn Shadden. LinFlo-Net: A Two-Stage Deep Learning Method to Generate Simulation Ready Meshes of the Heart. Journal of Biomechanical Engineering, 146(071005), March 2024. ISSN 0148-0731. doi:10.1115/1.4064527. URL https://doi.org/10.1115/1.4064527.

[26] Daniel H. Pak, Minliang Liu, Theodore Kim, Liang Liang, Andres Caballero, John Onofrey, Shawn S. Ahn, Yilin Xu, Raymond McKay, Wei Sun, Rudolph Gleason, and James S. Duncan. Patient-Specific Heart Geometry Modeling for Solid Biomechanics Using Deep Learning. IEEE Transactions on Medical Imaging, 43(1):203–215, January 2024. ISSN 1558-254X. doi:10.1109/TMI.2023.3294128. URL https://ieeexplore.ieee.org/abstract/document/10178071. xConference Name: IEEE Transactions on Medical Imaging.

[27] Martin R. Pfaller, Julia M. Hörmann, Martina Weigl, Andreas Nagler, Radomir Chabiniok, Cristóbal Bertoglio, and Wolfgang A. Wall. The importance of the pericardium for cardiac biomechanics: from physiology to computational modeling. Biomechanics and Modeling in Mechanobiology, 18(2):503–529, April 2019. ISSN 1617-7940. doi:10.1007/s10237-018-1098-4. URL https://doi.org/10.1007/s10237-018-1098-4.

[28] Sheikh Mohammad Shavik, Christopher Tossas-Betancourt, C. Alberto Figueroa, Seungik Baek, and Lik Chuan Lee. Multiscale Modeling Framework of Ventricular-Arterial Bi-directional Interactions in the Cardiopulmonary Circulation. Frontiers in Physiology, 11, January 2020. ISSN 1664-042X. doi:10.3389/fphys.2020.00002. URL https://www.frontiersin.org https://www.frontiersin.org/journals/physiology/articles/10.3389/fphys.2020.00002/full. Publisher: Frontiers.

[29] F. Levrero-Florencio, F. Margara, E. Zacur, A. Bueno-Orovio, Z. J. Wang, A. Santiago, J. Aguado-Sierra, G. Houzeaux, V. Grau, D. Kay, M. Vázquez, R. Ruiz-Baier, and B. Rodriguez. Sensitivity analysis of a strongly-coupled human-based electromechanical cardiac model: Effect of mechanical parameters on physiologically relevant biomarkers. Computer Methods in Applied Mechanics and Engineering, 361:112762, April 2020. ISSN 0045-7825. doi:10.1016/j.cma.2019.112762. URL https://www.sciencedirect.com/science/article/pii/S0045782519306541.

[30] Matteo Salvador, Francesco Regazzoni, Luca Dede’, and Alfio Quarteroni. Fast and robust parameter estimation with uncertainty quantification for the cardiac function. Computer Methods and Programs in Biomedicine, 231: 107402, April 2023. ISSN 0169-2607. doi:10.1016/j.cmpb.2023.107402. URL https://www.sciencedirect.com/science/article/pii/S016926072300069X.

[31] Andrea Tonini, Francesco Regazzoni, Matteo Salvador, Luca Dede’, Roberto Scrofani, Laura Fusini, Chiara Cogliati, Gianluca Pontone, Christian Vergara, and Alfio Quarteroni. Two New Calibration Techniques of Lumped-Parameter Mathematical Models for the Cardiovascular System. International Journal for Numerical Methods in Engineering, 126(1):e7648, 2025. ISSN 1097-0207. doi:10.1002/nme.7648. URL https://onlinelibrary.wiley.com/doi/abs/10.1002/nme.7648. _eprint: https://onlinelibrary.wiley.com/doi/pdf/10.1002/nme.7648.

[32] Marina Strocchi, Christoph M Augustin, Matthias A F Gsell, Christopher A Rinaldi, Edward J Vigmond, Gernot Plank, Chris J Oates, Richard D Wilkinson, and Steven A Niederer. Integrating imaging and invasive pressure data into a multi-scale whole-heart model. Journal of Biomechanical Engineering, pages 1–52, August 2025. ISSN 0148-0731. doi:10.1115/1.4069497. URL https://doi.org/10.1115/1.4069497.

[33] Marc Hirschvogel, Marina Bassilious, Lasse Jagschies, Stephen M. Wildhirt, and Michael W. Gee. A monolithic 3D-0D coupled closed-loop model of the heart and the vascular system: Experiment-based parameter estimation for patient-specific cardiac mechanics. International Journal for Numerical Methods in Biomedical Engineering, 33(8):e2842, 2017. ISSN 2040-7947. doi:10.1002/cnm.2842. URL https://onlinelibrary.wiley.com/doi/abs/10.1002/cnm.2842. _eprint: https://onlinelibrary.wiley.com/doi/pdf/10.1002/cnm.2842.

[34] Zhinuo Jenny Wang, Maxx Holmes, Ruben Doste, Julia Camps, Francesca Margara, Mariano Vazquez, and Blanca Rodriguez. Calibration and validation strategy for electromechanical cardiac digital twins, March 2025. URL https://www.biorxiv.org/content/10.1101/2025.03.06.638897v1. xPages: 2025.03.06.638897 Section: New Results.

[35] Marina Strocchi, Stefano Longobardi, Christoph M. Augustin, Matthias A. F. Gsell, Argyrios Petras, Christopher A. Rinaldi, Edward J. Vigmond, Gernot Plank, Chris J. Oates, Richard D. Wilkinson, and Steven A. Niederer. Cell to whole organ global sensitivity analysis on a four-chamber heart electromechanics model using Gaussian processes emulators. PLOS Computational Biology, 19(6):e1011257, June 2023. ISSN 1553-7358. doi:10.1371/journal.pcbi.1011257. URL https://journals.plos.org/ploscompbiol/article?id=10.1371/journal.pcbi.1011257. Publisher: Public Library of Science.

[36] Lei Shi, Yurui Chen, and Vijay Vedula. HeartSimSage: Attention-Enhanced Graph Neural Networks for Accelerating Cardiac Mechanics Modeling, April 2025. URL http://arxiv.org/abs/2504.18968. 2504.18968 [physics].

[37] Federica Caforio, Francesco Regazzoni, Stefano Pagani, Elias Karabelas, Christoph Augustin, Gundolf Haase, Gernot Plank, and Alfio Quarteroni. Physics-informed neural network estimation of material properties in soft tissue nonlinear biomechanical models. Computational Mechanics, 75(2):487–513, February 2025. ISSN 1432-0924. doi:10.1007/s00466-024-02516-x. URL https://link.springer.com/article/10.1007/s00466-024-02516-x. Company: Springer Distributor: Springer Institution: Springer Label: Springer Number: 2 Publisher: Springer Berlin Heidelberg.

[38] Matthias Höfler, Francesco Regazzoni, Stefano Pagani, Elias Karabelas, Christoph Augustin, Gundolf Haase, Gernot Plank, and Federica Caforio. Physics-informed neural network estimation of active material properties in time-dependent cardiac biomechanical models, May 2025. URL http://arxiv.org/abs/2505.03382. 2505.03382 [cs].

[39] Shruti Motiwale, Wenbo Zhang, Reese Feldmeier, and Michael S. Sacks. A neural network finite element approach for high speed cardiac mechanics simulations. Computer Methods in Applied Mechanics and Engineering, 427:117060, July 2024. ISSN 0045-7825. doi:10.1016/j.cma.2024.117060. URL https://www.sciencedirect.com/science/article/pii/S0045782524003165.

[40] Benjamin Thomas, Christian Goodbrake, Kenneth Meyer, and Michael Sacks. CARDIAX: A JAX-based platform for Rapid Cardiac Functional Simulations, July 2025. URL https://engrxiv.org/preprint/view/4923.

[41] Thomas Fritz, Christian Wieners, Gunnar Seemann, Henning Steen, and Olaf Dössel. Simulation of the contraction of the ventricles in a human heart model including atria and pericardium. Biomechanics and Modeling in Mechanobiology, 13(3):627–641, June 2014. ISSN 1617-7940. doi:10.1007/s10237-013-0523-y. URL https://doi.org/10.1007/s10237-013-0523-y.

[42] Marina Strocchi, Matthias A. F. Gsell, Christoph M. Augustin, Orod Razeghi, Caroline H. Roney, Anton J. Prassl, Edward J. Vigmond, Jonathan M. Behar, Justin S. Gould, Christopher A. Rinaldi, Martin J. Bishop, Gernot Plank, and Steven A. Niederer. Simulating ventricular systolic motion in a four-chamber heart model with spatially varying robin boundary conditions to model the effect of the pericardium. Journal of Biomechanics, 101:109645, March 2020. ISSN 0021-9290. doi:10.1016/j.jbiomech.2020.109645. URL https://www.sciencedirect.com/science/article/pii/S002192902030052X.

[43] M. Peirlinck, F. Sahli Costabal, J. Yao, J. M. Guccione, S. Tripathy, Y. Wang, D. Ozturk, P. Segars, T. M. Morrison, S. Levine, and E. Kuhl. Precision medicine in human heart modeling. Biomechanics and Modeling in Mechanobiology, 20(3):803–831, June 2021. ISSN 1617-7940. doi:10.1007/s10237-021-01421-z. URL https://doi.org/10.1007/s10237-021-01421-z.

[44] Brian Baillargeon, Nuno Rebelo, David D. Fox, Robert L. Taylor, and Ellen Kuhl. The Living Heart Project: A robust and integrative simulator for human heart function. European journal of mechanics. A, Solids, 48:38–47, 2014. ISSN 0997-7538. doi:10.1016/j.euromechsol.2014.04.001. URL https://www.ncbi.nlm.nih.gov/pmc/articles/PMC4175454/.

[45] Tobias Gerach, Steffen Schuler, Jonathan Fröhlich, Laura Lindner, Ekaterina Kovacheva, Robin Moss, Eike Moritz Wülfers, Gunnar Seemann, Christian Wieners, and Axel Loewe. Electro-Mechanical Whole-Heart Digital Twins: A Fully Coupled Multi-Physics Approach. Mathematics, 9(11):1247, January 2021. ISSN 2227-7390. doi:10.3390/math9111247. URL https://www.mdpi.com/2227-7390/9/11/1247. Number: 11 Publisher: Multidisciplinary Digital Publishing Institute.

[46] Marco Fedele, Roberto Piersanti, Francesco Regazzoni, Matteo Salvador, Pasquale Claudio Africa, Michele Bucelli, Alberto Zingaro, Luca Dede’, and Alfio Quarteroni. A comprehensive and biophysically detailed computational model of the whole human heart electromechanics. Computer Methods in Applied Mechanics and Engineering, 410:115983, May 2023. ISSN 0045-7825. doi:10.1016/j.cma.2023.115983. URL https://www.sciencedirect.com/science/article/pii/S0045782523001068.

[47] Justina Ghebryal, Cristobal Rodero, Rosie K. Barrows, Marina Strocchi, Caroline H. Roney, Christoph M. Augustin, Gernot Plank, and Steven A. Niederer. Effect of Varying Pericardial Boundary Conditions on Whole Heart Function: A Computational Study. In Olivier Bernard, Patrick Clarysse, Nicolas Duchateau, Jacques Ohayon, and Magalie Viallon, editors, Functional Imaging and Modeling of the Heart, pages 545–554, Cham, 2023. Springer Nature Switzerland. ISBN 978-3-031-35302-4. doi:10.1007/978-3-031-35302-4_56.

[48] Liuyang Feng, Hao Gao, and Xiaoyu Luo. Whole-heart modelling with valves in a fluid–structure interaction framework. Computer Methods in Applied Mechanics and Engineering, 420:116724, February 2024. ISSN 0045-7825. doi:10.1016/j.cma.2023.116724. URL https://www.sciencedirect.com/science/article/pii/S0045782523008472.

[49] L. Asner, M. Hadjicharalambous, R. Chabiniok, D. Peressutti, E. Sammut, J. Wong, G. Carr-White, R. Razavi, A. P. King, N. Smith, J. Lee, and D. Nordsletten. Patient-specific modeling for left ventricular mechanics using data-driven boundary energies. Computer Methods in Applied Mechanics and Engineering, 314:269–295, February 2017. ISSN 0045-7825. doi:10.1016/j.cma.2016.08.002. URL https://www.sciencedirect.com/science/article/pii/S0045782516308672.

[50] Mathias Peirlinck, Kevin L. Sack, Pieter De Backer, Pedro Morais, Patrick Segers, Thomas Franz, and Matthieu De Beule. Kinematic boundary conditions substantially impact in silico ventricular function. International Journal for Numerical Methods in Biomedical Engineering, 35(1):e3151, 2019. ISSN 2040-7947. doi:10.1002/cnm.3151. URL https://onlinelibrary.wiley.com/doi/abs/10.1002/cnm.3151. _eprint: https://onlinelibrary.wiley.com/doi/pdf/10.1002/cnm.3151.

[51] Tahar Arjoune, Christian Bilas, Christian Meierhofer, Heiko Stern, Peter Ewert, and Michael W. Gee. Inverse analysis of patient-specific parameters of a 3D–0D closed-loop cardiovascular model with an exemplary application to an adult tetralogy of Fallot case. Biomechanics and Modeling in Mechanobiology, September 2025. ISSN 1617-7940. doi:10.1007/s10237-025-02006-w. URL https://doi.org/10.1007/s10237-025-02006-w.

[52] Christoph M. Augustin, Matthias A. F. Gsell, Elias Karabelas, Erik Willemen, Frits W. Prinzen, Joost Lumens, Edward J. Vigmond, and Gernot Plank. A computationally efficient physiologically comprehensive 3D–0D closedloop model of the heart and circulation. Computer Methods in Applied Mechanics and Engineering, 386:114092, December 2021. ISSN 0045-7825. doi:10.1016/j.cma.2021.114092. URL https://www.sciencedirect.com/science/article/pii/S0045782521004230.

[53] Renee Miller, Eric Kerfoot, Charlène Mauger, Tevfik F. Ismail, Alistair A. Young, and David A. Nordsletten. An Implementation of Patient-Specific Biventricular Mechanics Simulations With a Deep Learning and Computational Pipeline. Frontiers in Physiology, 12, September 2021. ISSN 1664-042X. doi:10.3389/fphys.2021.716597. URL https://www.frontiersin.org/journals/physiology/articles/10.3389/fphys.2021.716597/full. Publisher: Frontiers.

[54] Jonathan Fröhlich, Tobias Gerach, Jonathan Krauß, Axel Loewe, Laura Stengel, and Christian Wieners. Numerical evaluation of elasto-mechanical and visco-elastic electro-mechanical models of the human heart. GAMM-Mitteilungen, 46(3-4):e202370010, 2023. ISSN 1522-2608. doi:10.1002/gamm.202370010. URL https://onlinelibrary.wiley.com/doi/abs/10.1002/gamm.202370010. _eprint: https://onlinelibrary.wiley.com/doi/pdf/10.1002/gamm.202370010.

[55] Roberto Piersanti, Francesco Regazzoni, Matteo Salvador, Antonio F. Corno, Luca Dede’, Christian Vergara, and Alfio Quarteroni. 3D–0D closed-loop model for the simulation of cardiac biventricular electromechanics. Computer Methods in Applied Mechanics and Engineering, 391:114607, March 2022. ISSN 0045-7825. doi:10.1016/j.cma.2022.114607. URL https://www.sciencedirect.com/science/article/pii/S0045782522000251.

[56] Oscar O. Odeigah, Ethan D. Kwan, Kristen M. Garcia, Henrik Finsberg, Daniela Valdez-Jasso, and Joakim Sundnes. A computational study of right ventricular mechanics in a rat model of pulmonary arterial hypertension. Frontiers in Physiology, 15, March 2024. ISSN 1664-042X. doi:10.3389/fphys.2024.1360389. URL https://www.frontiersin.org/journals/physiology/articles/10.3389/fphys.2024.1360389/full. Publisher: Frontiers.

[57] Emilio A. Mendiola, Sunder Neelakantan, Qian Xiang, Samer Merchant, Ke Li, Edward W. Hsu, Richard A. F. Dixon, Peter Vanderslice, and Reza Avazmohammadi. Contractile Adaptation of the Left Ventricle Post-myocardial Infarction: Predictions by Rodent-Specific Computational Modeling. Annals of Biomedical Engineering, 51(4):846–863, April 2023. ISSN 1573-9686. doi:10.1007/s10439-022-03102-z. URL https://doi.org/10.1007/s10439-022-03102-z.

[58] Y. D. Motchon, Kevin L. Sack, M. S. Sirry, M. Kruger, E. Pauwels, D. Van Loo, A. De Muynck, L. Van Hoorebeke, Neil H. Davies, and Thomas Franz. Effect of biomaterial stiffness on cardiac mechanics in a biventricular infarcted rat heart model with microstructural representation of in situ intramyocardial injectate. International Journal for Numerical Methods in Biomedical Engineering, 39(5):e3693, 2023. ISSN 2040-7947. doi:10.1002/cnm.3693. URL https://onlinelibrary.wiley.com/doi/abs/10.1002/cnm.3693. _eprint: https://onlinelibrary.wiley.com/doi/pdf/10.1002/cnm.3693.

[59] Joy Mojumder, Lei Fan, Thuy Nguyen, Kenneth S. Campbell, Jonathan F. Wenk, Julius M. Guccione, Theodore Abraham, and Lik Chuan Lee. Computational analysis of ventricular mechanics in hypertrophic cardiomyopathy patients. Scientific Reports, 13(1):958, January 2023. ISSN 2045-2322. doi:10.1038/s41598-023-28037-w. URL https://www.nature.com/articles/s41598-023-28037-w. Publisher: Nature Publishing Group.

[60] Nicolás Laita, Ricardo M. Rosales, Ming Wu, Piet Claus, Stefan Janssens, Miguel Ángel Martínez, Manuel Doblaré, and Estefanía Peña. On modeling the in vivo ventricular passive mechanical behavior from in vitro experimental properties in porcine hearts. Computers & Structures, 292:107241, February 2024. ISSN 0045-7949. doi:10.1016/j.compstruc.2023.107241. URL https://www.sciencedirect.com/science/article/pii/S0045794923002717.

[61] Fikunwa O. Kolawole, Vicky Y. Wang, Bianca Freytag, Michael Loecher, Tyler E. Cork, Martyn P. Nash, Ellen Kuhl, and Daniel B. Ennis. Characterizing variability in passive myocardial stiffness in healthy human left ventricles using personalized MRI and finite element modeling. Scientific Reports, 15(1):5556, February 2025. ISSN 2045-2322. doi:10.1038/s41598-025-89243-2. URL https://www.nature.com/articles/s41598-025-89243-2. Publisher: Nature Publishing Group.

[62] David Holz, Denisa Martonová, Emely Schaller, Minh Tuan Duong, Muhannad Alkassar, Michael Weyand, and Sigrid Leyendecker. Transmural fibre orientations based on Laplace–Dirichlet-Rule-Based-Methods and their influence on human heart simulations. Journal of Biomechanics, 156:111643, July 2023. ISSN 0021-9290. doi:10.1016/j.jbiomech.2023.111643. URL https://www.sciencedirect.com/science/article/pii/S0021929023002129.

[63] Will Zhang, Javiera Jilberto, Gerhard Sommer, Michael S. Sacks, Gerhard A. Holzapfel, and David A. Nordsletten. Simulating hyperelasticity and fractional viscoelasticity in the human heart. Computer Methods in Applied Mechanics and Engineering, 411:116048, June 2023. ISSN 0045-7825. doi:10.1016/j.cma.2023.116048. URL https://www.sciencedirect.com/science/article/pii/S004578252300172X.

[64] Alfio Quarteroni, Toni Lassila, Simone Rossi, and Ricardo Ruiz-Baier. Integrated Heart—Coupling multiscale and multiphysics models for the simulation of the cardiac function. Computer Methods in Applied Mechanics and Engineering, 314:345–407, February 2017. ISSN 0045-7825. doi:10.1016/j.cma.2016.05.031. URL https://www.sciencedirect.com/science/article/pii/S0045782516304662.

[65] R. Verzicco. Electro-fluid-mechanics of the heart. Journal of Fluid Mechanics, 941:P1, June 2022. ISSN 0022-1120, 1469-7645. doi:10.1017/jfm.2022.272. URL https://www.cambridge.org/core/journals/journal-of-fluid-mechanics/article/electrofluidmechanics-of-the-heart/6BDA989AA6F798872F10624C6D786B09.

[66] Robert Naeije and Roberto Badagliacca. The overloaded right heart and ventricular interdependence. Cardiovascular Research, 113(12):1474–1485, October 2017. ISSN 0008-6363. doi:10.1093/cvr/cvx160. URL https://doi.org/10.1093/cvr/cvx160.

[67] Fanny Vaillant, Emma Abell, Laura R. Bear, Guido Caluori, Charly Belterman, Ruben Coronel, Sylvain Ploux, and Pierre Dos Santos. Influence of pericardium on ventricular mechanical interdependence in an isolated biventricular working pig heart model. The Journal of Physiology, 603(2):285–300, 2025. ISSN 1469-7793. doi:10.1113/JP286259. URL https://onlinelibrary.wiley.com/doi/abs/10.1113/JP286259. _eprint: https://onlinelibrary.wiley.com/doi/pdf/10.1113/JP286259.

[68] R. J. Damiano, P. La Follette, J. L. Cox, J. E. Lowe, and W. P. Santamore. Significant left ventricular contribution to right ventricular systolic function. American Journal of Physiology-Heart and Circulatory Physiology, 261 (5):H1514–H1524, November 1991. ISSN 0363-6135. doi:10.1152/ajpheart.1991.261.5.H1514. URL https://journals.physiology.org/doi/abs/10.1152/ajpheart.1991.261.5.H1514. Publisher: American Physiological Society.

[69] Matthieu Petit and Antoine Vieillard-Baron. Ventricular interdependence in critically ill patients: from physiology to bedside. Frontiers in Physiology, 14:1232340, August 2023. ISSN 1664-042X. doi:10.3389/fphys.2023.1232340. URL https://www.ncbi.nlm.nih.gov/pmc/articles/PMC10442576/.

[70] Salla M. Kim, E. Benjamin Randall, Filip Jezek, Daniel A. Beard, and Naomi C. Chesler. Computational modeling of ventricular-ventricular interactions suggest a role in clinical conditions involving heart failure. Frontiers in Physiology, 14, September 2023. ISSN 1664-042X. doi:10.3389/fphys.2023.1231688. URL https://www.frontiersin.org/journals/physiology/articles/10.3389/fphys.2023.1231688/full. Publisher: Frontiers.

[71] Kevin L. Sack, Yaghoub Dabiri, Thomas Franz, Scott D. Solomon, Daniel Burkhoff, and Julius M. Guccione. Investigating the Role of Interventricular Interdependence in Development of Right Heart Dysfunction During LVAD Support: A Patient-Specific Methods-Based Approach. Frontiers in Physiology, 9, May 2018. ISSN 1664-042X. doi:10.3389/fphys.2018.00520. URL https://www.frontiersin.org/journals/physiology/articles/10.3389/fphys.2018.00520/full. Publisher: Frontiers.

[72] R. Beyar and S. Sideman. Atrioventricular interactions: a theoretical simulation study. American Journal of Physiology-Heart and Circulatory Physiology, 252(3):H653–H665, March 1987. ISSN 0363-6135. doi:10.1152/ajpheart.1987.252.3.H653. URL https://journals.physiology.org/doi/abs/10.1152/ajpheart.1987.252.3.H653. Publisher: American Physiological Society.

[73] Gustavo G. Blume, Christopher J. Mcleod, Marion E. Barnes, James B. Seward, Patricia A. Pellikka, Paul M. Bastiansen, and Teresa S.M. Tsang. Left atrial function: physiology, assessment, and clinical implications. European Journal of Echocardiography, 12(6):421–430, June 2011. ISSN 1525-2167. doi:10.1093/ejechocard/jeq175. URL https://doi.org/10.1093/ejechocard/jeq175.

[74] S. Land and S.A. Niederer. Influence of atrial contraction dynamics on cardiac function. International Journal for Numerical Methods in Biomedical Engineering, 34(3), 2018. doi:10.1002/cnm.2931.

[75] Tobias Gerach, Steffen Schuler, Andreas Wachter, and Axel Loewe. The Impact of Standard Ablation Strategies for Atrial Fibrillation on Cardiovascular Performance in a Four-Chamber Heart Model. Cardiovascular Engineering and Technology, 14(2):296–314, April 2023. ISSN 1869-4098. doi:10.1007/s13239-022-00651-1. URL https://doi.org/10.1007/s13239-022-00651-1.

[76] Alfonso Santiago, Alfonso Santiago, Jazmín Aguado-Sierra, Jazmin Aguado-Sierra, Miguel Zavala-Aké, Miguel Zavala-Aké, Ruben Doste, Rubén Doste, Ruben Doste-Beltran, Samuel Gómez, Samuel Gómez, Ruth Arís, Ruth Arís, Juan Carlos Cajas, J.C. Cajas, Eva Casoni, Eva Casoni, Mariano Vázquez, and Mariano Vázquez. Fully coupled fluid-electro-mechanical model of the human heart for supercomputers. International Journal for Numerical Methods in Biomedical Engineering, 34(12), December 2018. doi:10.1002/cnm.3140. MAG ID: 2885942414.

[77] Cristobal Rodero, Marina Strocchi, Maciej Marciniak, Stefano Longobardi, John Whitaker, Mark D. O’Neill, Karli Gillette, Christoph Augustin, Gernot Plank, Edward J. Vigmond, Pablo Lamata, and Steven A. Niederer. Linking statistical shape models and simulated function in the healthy adult human heart. PLOS Computational Biology, 17(4):e1008851, April 2021. ISSN 1553-7358. doi:10.1371/journal.pcbi.1008851. URL https://journals.plos.org/ploscompbiol/article?id=10.1371/journal.pcbi.1008851. Publisher: Public Library of Science.

[78] Arnab Palit, Sunil K. Bhudia, Theodoros N. Arvanitis, Glen A. Turley, and Mark A. Williams. Computational modelling of left-ventricular diastolic mechanics: Effect of fibre orientation and right-ventricle topology. Journal of Biomechanics, 48(4):604–612, February 2015. ISSN 0021-9290. doi:10.1016/j.jbiomech.2014.12.054. URL https://www.sciencedirect.com/science/article/pii/S0021929015000020.

[79] Steven Niederer, Kawal Rhode, Reza Razavi, and Nic Smith. The Importance of Model Parameters and Boundary Conditions in Whole Organ Models of Cardiac Contraction. In Nicholas Ayache, Hervé Delingette, and Maxime Sermesant, editors, Functional Imaging and Modeling of the Heart, pages 348–356, Berlin, Heidelberg, 2009. Springer. ISBN 978-3-642-01932-6. doi:10.1007/978-3-642-01932-6_38.

[80] F. Dorri, P. F. Niederer, and P. P. Lunkenheimer. A finite element model of the human left ventricular systole. Computer Methods in Biomechanics and Biomedical Engineering, 9(5):319–341, October 2006. ISSN 1025-5842. doi:10.1080/10255840600960546. URL https://doi.org/10.1080/10255840600960546. Publisher: Taylor & Francis _eprint: https://doi.org/10.1080/10255840600960546.

[81] Martin Genet, Lik Chuan Lee, Liang Ge, Gabriel Acevedo-Bolton, Nick Jeung, Alastair Martin, Neil Cambronero, Andrew Boyle, Yerem Yeghiazarians, Sebastian Kozerke, and Julius M. Guccione. A Novel Method for Quantifying Smooth Regional Variations in Myocardial Contractility Within an Infarcted Human Left Ventricle Based on Delay-Enhanced Magnetic Resonance Imaging. Journal of Biomechanical Engineering, 137(8):081009, August 2015. ISSN 0148-0731, 1528-8951. doi:10.1115/1.4030667. URL https://asmedigitalcollection.asme.org/biomechanical/article/doi/10.1115/1.4030667/422020/A-Novel-Method-for-Quantifying-Smooth-Regional.

[82] Sander Land, Viatcheslav Gurev, Sander Arens, Christoph M. Augustin, Lukas Baron, Robert Blake, Chris Bradley, Sebastian Castro, Andrew Crozier, Marco Favino, Thomas E. Fastl, Thomas Fritz, Hao Gao, Alessio Gizzi, Boyce E. Griffith, Daniel E. Hurtado, Rolf Krause, Xiaoyu Luo, Martyn P. Nash, Simone Pezzuto, Gernot Plank, Simone Rossi, Daniel Ruprecht, Gunnar Seemann, Nicolas P. Smith, Joakim Sundnes, J. Jeremy Rice, Natalia Trayanova, Dafang Wang, Zhinuo Jenny Wang, and Steven A. Niederer. Verification of cardiac mechanics software: benchmark problems and solutions for testing active and passive material behaviour. Proceedings. Mathematical, Physical, and Engineering Sciences, 471(2184):20150641, December 2015. ISSN 1364-5021. doi:10.1098/rspa.2015.0641.

[83] Julius M. Guccione, Kevin D. Costa, and Andrew D. McCulloch. Finite element stress analysis of left ventricular mechanics in the beating dog heart. Journal of Biomechanics, 28(10):1167–1177, October 1995. ISSN 00219290. doi:10.1016/0021-9290(94)00174-3. URL https://linkinghub.elsevier.com/retrieve/pii/0021929094001743.

[84] Adarsh Krishnamurthy, Christopher T. Villongco, Joyce Chuang, Lawrence R. Frank, Vishal Nigam, Ernest Belezzuoli, Paul Stark, David E. Krummen, Sanjiv Narayan, Jeffrey H. Omens, Andrew D. McCulloch, and Roy C. P. Kerckhoffs. Patient-specific models of cardiac biomechanics. Journal of Computational Physics, 244:4–21, July 2013. ISSN 0021-9991. doi:10.1016/j.jcp.2012.09.015. URL https://www.sciencedirect.com/science/article/pii/S0021999112005463.

[85] Aaron L. Brown, Matteo Salvador, Lei Shi, Martin R. Pfaller, Zinan Hu, Kaitlin E. Harold, Tzung Hsiai, Vijay Vedula, and Alison L. Marsden. A modular framework for implicit 3D–0D coupling in cardiac mechanics. Computer Methods in Applied Mechanics and Engineering, 421:116764, March 2024. ISSN 0045-7825. doi:10.1016/j.cma.2024.116764. URL https://www.sciencedirect.com/science/article/pii/S0045782524000203.

[86] F. Regazzoni, L. Dedè, and A. Quarteroni. Machine learning of multiscale active force generation models for the efficient simulation of cardiac electromechanics. Computer Methods in Applied Mechanics and Engineering, 370:113268, October 2020. ISSN 0045-7825. doi:10.1016/j.cma.2020.113268. URL https://www.sciencedirect.com/science/article/pii/S0045782520304539.

[87] F. Regazzoni, M. Salvador, P. C. Africa, M. Fedele, L. Dedè, and A. Quarteroni. A cardiac electromechanical model coupled with a lumped-parameter model for closed-loop blood circulation. Journal of Computational Physics, 457:111083, May 2022. ISSN 0021-9991. doi:10.1016/j.jcp.2022.111083. URL https://www.sciencedirect.com/science/article/pii/S0021999122001450.

[88] Emilio A. Mendiola, Michael S. Sacks, and Reza Avazmohammadi. Mechanical Interaction of the Pericardium and Cardiac Function in the Normal and Hypertensive Rat Heart. Frontiers in Physiology, 13, May 2022. ISSN 1664-042X. doi:10.3389/fphys.2022.878861. URL https://www.frontiersin.org/journals/physiology/articles/10.3389/fphys.2022.878861/full. Publisher: Frontiers.

[89] Jonathan Krauß, Tobias Gerach, and Axel Loewe. Comparison of Pericardium Modeling Approaches for Mechanical Whole Heart Simulations. In 2023 Computing in Cardiology (CinC), volume 50, pages 1–4, October 2023. doi:10.22489/CinC.2023.150. URL https://ieeexplore.ieee.org/abstract/document/10363870. ISSN: 2325-887X.

[90] Myrianthi Hadjicharalambous, Christian T. Stoeck, Miriam Weisskopf, Nikola Cesarovic, Eleftherios Ioannou, Vasileios Vavourakis, and David A. Nordsletten. Investigating the reference domain influence in personalised models of cardiac mechanics. Biomechanics and Modeling in Mechanobiology, 20(4):1579–1597, August 2021. ISSN 1617-7940. doi:10.1007/s10237-021-01464-2. URL https://doi.org/10.1007/s10237-021-01464-2.

[91] Javiera Jilberto and David Nordsletten. Identification of the Unloaded Heart Configuration Including External Interactions. In Radomír Chabiniok, Qing Zou, Tarique Hussain, Hoang H. Nguyen, Vlad G. Zaha, and Maria Gusseva, editors, Functional Imaging and Modeling of the Heart, pages 331–342, Cham, 2025. Springer Nature Switzerland. ISBN 978-3-031-94559-5. doi:10.1007/978-3-031-94559-5_30.

[92] Lei Shi, Boyang Gan, Ian Y. Chen, and Vijay Vedula. Personalized multiscale modeling of left atrial mechanics and blood flow. Computer Methods in Applied Mechanics and Engineering, 448:118412, January 2026. ISSN 0045-7825. doi:10.1016/j.cma.2025.118412. URL https://www.sciencedirect.com/science/article/pii/S004578252500684X.

[93] J. D. Bayer, R. C. Blake, G. Plank, and N. A. Trayanova. A novel rule-based algorithm for assigning myocardial fiber orientation to computational heart models. Annals of Biomedical Engineering, 40(10):2243–2254, October 2012. ISSN 1573-9686. doi:10.1007/s10439-012-0593-5.

[94] Roberto Piersanti, Pasquale C. Africa, Marco Fedele, Christian Vergara, Luca Dedè, Antonio F. Corno, and Alfio Quarteroni. Modeling cardiac muscle fibers in ventricular and atrial electrophysiology simulations. Computer Methods in Applied Mechanics and Engineering, 373:113468, January 2021. ISSN 00457825. doi:10.1016/j.cma.2020.113468. URL https://linkinghub.elsevier.com/retrieve/pii/S0045782520306538.

[95] Christopher M. Kramer, Jörg Barkhausen, Chiara Bucciarelli-Ducci, Scott D. Flamm, Raymond J. Kim, and Eike Nagel. Standardized cardiovascular magnetic resonance imaging (CMR) protocols: 2020 update. Journal of Cardiovascular Magnetic Resonance, 22(1):17, February 2020. ISSN 1532-429X. doi:10.1186/s12968-020-00607-1. URL https://doi.org/10.1186/s12968-020-00607-1.

[96] David Amar, Jose A. Melendez, Hao Zhang, Catherine Dobres, Denis H. Y. Leung, and Roger E. Padilla. Correlation of peripheral venous pressure and central venous pressure in surgical patients. Journal of Cardiothoracic and Vascular Anesthesia, 15(1):40–43, February 2001. ISSN 1053-0770. doi:10.1053/jcan.2001.20271. URL https://www.sciencedirect.com/science/article/pii/S1053077001020419.

[97] Alexandre Mebazaa, Peter Karpati, Estelle Renaud, and Lars Algotsson. Acute right ventricular failure—from pathophysiology to new treatments. Intensive Care Medicine, 30(2):185–196, February 2004. ISSN 1432-1238. doi:10.1007/s00134-003-2025-3. URL https://link-springer-com.laneproxy.stanford.edu/article/10.1007/s00134-003-2025-3. Company: Springer Distributor: Springer Institution: Springer Label: Springer Number: 2 Publisher: Springer Berlin Heidelberg.

[98] Table:Normal Pressures in the Heart and Great Vessels. URL https://www.merckmanuals.com/professional/multimedia/table/normal-pressures-in-the-heart-and-great-vessels.

[99] Ian Y. Chen, Vijay Vedula, Sachin B. Malik, Tie Liang, Andrew Y. Chang, Kieran S. Chung, Nazish Sayed, Philip S. Tsao, John C. Giacomini, Alison L. Marsden, and Joseph C. Wu. Preoperative Computed Tomography Angiography Reveals Leaflet-Specific Calcification and Excursion Patterns in Aortic Stenosis. Circulation: Cardiovascular Imaging, 14(12):1122–1132, December 2021. doi:10.1161/CIRCIMAGING.121.012884. URL https://www.ahajournals.org/doi/10.1161/CIRCIMAGING.121.012884. Publisher: American Heart Association.

[100] Andriy Fedorov, Reinhard Beichel, Jayashree Kalpathy-Cramer, Julien Finet, Jean-Christophe Fillion-Robin, Sonia Pujol, Christian Bauer, Dominique Jennings, Fiona Fennessy, Milan Sonka, John Buatti, Stephen Aylward, James V. Miller, Steve Pieper, and Ron Kikinis. 3D Slicer as an image computing platform for the Quantitative Imaging Network. Magnetic Resonance Imaging, 30(9):1323–1341, November 2012. ISSN 0730-725X. doi:10.1016/j.mri.2012.05.001. URL https://www.sciencedirect.com/science/article/pii/S0730725X12001816.

[101] Adam Updegrove, Nathan M. Wilson, Jameson Merkow, Hongzhi Lan, Alison L. Marsden, and Shawn C. Shadden. SimVascular: An Open Source Pipeline for Cardiovascular Simulation. Annals of Biomedical Engineering, 45(3):525–541, March 2017. ISSN 1573-9686. doi:10.1007/s10439-016-1762-8. URL https://link.springer.com/article/10.1007/s10439-016-1762-8. Company: Springer Distributor: Springer Institution: Springer Label: Springer Number: 3 Publisher: Springer US.

[102] Hang Si. TetGen, a Delaunay-Based Quality Tetrahedral Mesh Generator. ACM Trans. Math. Softw., 41(2):11:1–11:36, February 2015. ISSN 0098-3500. doi:10.1145/2629697. URL https://dl.acm.org/doi/10.1145/2629697.

[103] Reidmen Aróstica, David Nolte, Aaron Brown, Amadeus Gebauer, Elias Karabelas, Javiera Jilberto, Matteo Salvador, Michele Bucelli, Roberto Piersanti, Kasra Osouli, Christoph Augustin, Henrik Finsberg, Lei Shi, Marc Hirschvogel, Martin Pfaller, Pasquale Claudio Africa, Matthias Gsell, Alison Marsden, David Nordsletten, Francesco Regazzoni, Gernot Plank, Joakim Sundnes, Luca Dede’, Mathias Peirlinck, Vijay Vedula, Wolfgang Wall, and Cristóbal Bertoglio. A software benchmark for cardiac elastodynamics. Computer Methods in Applied Mechanics and Engineering, 435:117485, February 2025. ISSN 0045-7825. doi:10.1016/j.cma.2024.117485. URL https://www.sciencedirect.com/science/article/pii/S0045782524007394.

[104] Gerhard A. Holzapfel and Ray W. Ogden. Constitutive Modelling of Passive Myocardium: A Structurally Based Framework for Material Characterization. Philosophical Transactions: Mathematical, Physical and Engineering Sciences, 367(1902):3445–3475, 2009. ISSN 1364-503X. URL https://www.jstor.org/stable/40485676. Publisher: Royal Society.

[105] Aaron L. Brown, Ju Liu, Daniel B. Ennis, and Alison L. Marsden. Cardiac mechanics modeling: recent developments and current challenges, September 2025. URL http://arxiv.org/abs/2509.07971. 2509.07971 [physics].

[106] W Schroeder, K Martin, and B Lorensen. The Visualization Toolkit An Object-Oriented Approach To 3D Graphics Edition 4.1, July 2018. URL https://gitlab.kitware.com/vtk/textbook/raw/master/VTKBook/VTKTextBook.pdf.

[107] Gillian M. Gunning and Bruce P. Murphy. Determination of the tensile mechanical properties of the segmented mitral valve annulus. Journal of Biomechanics, 47(2):334–340, January 2014. ISSN 0021-9290. doi:10.1016/j.jbiomech.2013.11.035. URL https://www.sciencedirect.com/science/article/pii/S0021929013005885.

[108] Michele Bucelli, Alberto Zingaro, Pasquale Claudio Africa, Ivan Fumagalli, Luca Dede’, and Alfio Quarteroni. A mathematical model that integrates cardiac electrophysiology, mechanics, and fluid dynamics: Application to the human left heart. International Journal for Numerical Methods in Biomedical Engineering, 39(3):e3678, 2023. ISSN 2040-7947. doi:10.1002/cnm.3678. URL https://onlinelibrary.wiley.com/doi/abs/10.1002/cnm.3678. _eprint: https://onlinelibrary.wiley.com/doi/pdf/10.1002/cnm.3678.

[109] Jason Bayer, Anton J. Prassl, Ali Pashaei, Juan F. Gomez, Antonio Frontera, Aurel Neic, Gernot Plank, and Edward J. Vigmond. Universal ventricular coordinates: A generic framework for describing position within the heart and transferring data. Medical Image Analysis, 45:83–93, April 2018. ISSN 13618415. doi:10.1016/j.media.2018.01.005. URL https://linkinghub.elsevier.com/retrieve/pii/S1361841518300203.

[110] Ju Liu and Alison L. Marsden. A unified continuum and variational multiscale formulation for fluids, solids, and fluid–structure interaction. Computer Methods in Applied Mechanics and Engineering, 337:549–597, August 2018. ISSN 0045-7825. doi:10.1016/j.cma.2018.03.045. URL https://www.sciencedirect.com/science/article/pii/S0045782518301701.

[111] Lei Shi, Ian Y. Chen, Hiroo Takayama, and Vijay Vedula. An optimization framework to personalize passive cardiac mechanics. Computer Methods in Applied Mechanics and Engineering, 432:117401, December 2024. ISSN 0045-7825. doi:10.1016/j.cma.2024.117401. URL https://www.sciencedirect.com/science/article/pii/S004578252400656X.

[112] Juan C. Simo and Robert L. Taylor. Quasi-incompressible finite elasticity in principal stretches. continuum basis and numerical algorithms. Computer Methods in Applied Mechanics and Engineering, 85(3):273–310, February 1991. ISSN 0045-7825. doi:10.1016/0045-7825(91)90100-K. URL https://www.sciencedirect.com/science/article/pii/004578259190100K.

[113] J. P. Mynard, M. R. Davidson, D. J. Penny, and J. J. Smolich. A simple, versatile valve model for use in lumped parameter and one-dimensional cardiovascular models. International Journal for Numerical Methods in Biomedical Engineering, 28(6-7):626–641, 2012. ISSN 2040-7947. doi:10.1002/cnm.1466. URL https://onlinelibrary.wiley.com/doi/abs/10.1002/cnm.1466. _eprint: https://onlinelibrary.wiley.com/doi/pdf/10.1002/cnm.1466.

[114] Tony F. Chan. An Approximate Newton Method for Coupled Nonlinear Systems. SIAM Journal on Numerical Analysis, 22(5):904–913, 1985. doi:10.1137/0722054. URL https://doi.org/10.1137/0722054. _eprint: https://doi.org/10.1137/0722054.

[115] Mahdi Esmaily Moghadam, Irene E. Vignon-Clementel, Richard Figliola, and Alison L. Marsden. A modular numerical method for implicit 0D/3D coupling in cardiovascular finite element simulations. Journal of Computational Physics, 244:63–79, July 2013. ISSN 0021-9991. doi:10.1016/j.jcp.2012.07.035. URL https://www.sciencedirect.com/science/article/pii/S0021999112004202.

[116] Marina Strocchi, Christoph M. Augustin, Matthias A. F. Gsell, Elias Karabelas, Aurel Neic, Karli Gillette, Orod Razeghi, Anton J. Prassl, Edward J. Vigmond, Jonathan M. Behar, Justin Gould, Baldeep Sidhu, Christopher A. Rinaldi, Martin J. Bishop, Gernot Plank, and Steven A. Niederer. A publicly available virtual cohort of four-chamber heart meshes for cardiac electro-mechanics simulations. PLOS ONE, 15(6):e0235145, June 2020. ISSN 1932-6203. doi:10.1371/journal.pone.0235145. URL https://journals.plos.org/plosone/article?id=10.1371/journal.pone.0235145. Publisher: Public Library of Science.

[117] Chi Zhu, Vijay Vedula, Dave Parker, Nathan Wilson, Shawn Shadden, and Alison Marsden. svFSI: A Multiphysics Package for Integrated Cardiac Modeling. Journal of Open Source Software, 7(78):4118, October 2022. ISSN 2475-9066. doi:10.21105/joss.04118. URL https://joss.theoj.org/papers/10.21105/joss.04118.

[118] Matteo Salvador and Alison Lesley Marsden. Branched Latent Neural Maps. Computer Methods in Applied Mechanics and Engineering, 418:116499, January 2024. ISSN 0045-7825. doi:10.1016/j.cma.2023.116499. URL https://www.sciencedirect.com/science/article/pii/S0045782523006230.

[119] Jongmin Seo, Daniele E. Schiavazzi, Andrew M. Kahn, and Alison L. Marsden. The effects of clinically-derived parametric data uncertainty in patient-specific coronary simulations with deformable walls. International Journal for Numerical Methods in Biomedical Engineering, 36(8):e3351, August 2020. ISSN 2040-7947. doi:10.1002/cnm.3351.

[120] Muhammad Owais Khan, Justin S. Tran, Han Zhu, Jack Boyd, René R. Sevag Packard, Ronald P. Karlsberg, Andrew M. Kahn, and Alison L. Marsden. Low Wall Shear Stress Is Associated with Saphenous Vein Graft Stenosis in Patients with Coronary Artery Bypass Grafting. Journal of Cardiovascular Translational Research, 14(4):770–781, August 2021. ISSN 1937-5395. doi:10.1007/s12265-020-09982-7.

[121] Vijay Vedula, Juhyun Lee, Hao Xu, C.-C. Jay Kuo, Tzung K. Hsiai, and Alison L. Marsden. A method to quantify mechanobiologic forces during zebrafish cardiac development using 4-D light sheet imaging and computational modeling. PLOS Computational Biology, 13(10):e1005828, October 2017. ISSN 1553-7358. doi:10.1371/journal.pcbi.1005828. URL https://journals.plos.org/ploscompbiol/article?id=10.1371/journal.pcbi.1005828. Publisher: Public Library of Science.

[122] Juhyun Lee, Vijay Vedula, Kyung In Baek, Junjie Chen, Jeffrey J. Hsu, Yichen Ding, Chih-Chiang Chang, Hanul Kang, Adam Small, Peng Fei, Cheng-ming Chuong, Rongsong Li, Linda Demer, René R. Sevag Packard, Alison L. Marsden, and Tzung K. Hsiai. Spatial and temporal variations in hemodynamic forces initiate cardiac trabeculation. JCI Insight, 3(13), July 2018. ISSN 0021-9738. doi:10.1172/jci.insight.96672. URL https://insight.jci.org/articles/view/96672. Publisher: American Society for Clinical Investigation.

[123] Kathrin Bäumler, Malte Rolf-Pissarczyk, Richard Schussnig, Thomas-Peter Fries, Gabriel Mistelbauer, Martin R. Pfaller, Alison L. Marsden, Dominik Fleischmann, and Gerhard A. Holzapfel. Assessment of Aortic Dissection Remodeling With Patient-Specific Fluid–Structure Interaction Models. IEEE Transactions on Biomedical Engineering, 72(3):953–964, March 2025. ISSN 1558-2531. doi:10.1109/TBME.2024.3480362. URL https://ieeexplore-ieee-org.stanford.idm.oclc.org/document/10716513.

[124] Zinan Hu, Jessica E. Herrmann, Erica L. Schwarz, Fannie M. Gerosa, Nir Emuna, Jay D. Humphrey, Adam W. Feinberg, Tain-Yen Hsia, Mark A. Skylar-Scott, and Alison L. Marsden. Multiphysics Simulations of a Bioprinted Pulsatile Fontan Conduit. Journal of Biomechanical Engineering, 147(071001), May 2025. ISSN 0148-0731. doi:10.1115/1.4068319. URL https://doi.org/10.1115/1.4068319.

[125] Erica L. Schwarz, Martin R. Pfaller, Jason M. Szafron, Marcos Latorre, Stephanie E. Lindsey, Christopher K. Breuer, Jay D. Humphrey, and Alison L. Marsden. A fluid–solid-growth solver for cardiovascular modeling. Computer Methods in Applied Mechanics and Engineering, 417:116312, December 2023. ISSN 0045-7825. doi:10.1016/j.cma.2023.116312. URL https://www.sciencedirect.com/science/article/pii/S004578252300436X.

[126] Martin R. Pfaller, Marcos Latorre, Erica L. Schwarz, Fannie M. Gerosa, Jason M. Szafron, Jay D. Humphrey, and Alison L. Marsden. FSGe: A fast and strongly-coupled 3D fluid–solid-growth interaction method. Computer Methods in Applied Mechanics and Engineering, 431:117259, November 2024. ISSN 0045-7825. doi:10.1016/j.cma.2024.117259. URL https://www.sciencedirect.com/science/article/pii/S0045782524005152.

[127] Youcef Saad and Martin H. Schultz. GMRES: A Generalized Minimal Residual Algorithm for Solving Nonsymmetric Linear Systems. SIAM Journal on Scientific and Statistical Computing, 7(3):856–869, July 1986. ISSN 0196-5204. doi:10.1137/0907058. URL https://epubs.siam.org/doi/10.1137/0907058. Publisher: Society for Industrial and Applied Mathematics.

[128] Mahdi Esmaily-Moghadam, Yuri Bazilevs, and Alison L. Marsden. A new preconditioning technique for implicitly coupled multidomain simulations with applications to hemodynamics. Computational Mechanics, 52(5):1141–1152, November 2013. ISSN 1432-0924. doi:10.1007/s00466-013-0868-1. URL https://link.springer.com/article/10.1007/s00466-013-0868-1. Company: Springer Distributor: Springer Institution: Springer Label: Springer Number: 5 Publisher: Springer Berlin Heidelberg.

[129] Pablo J. Blanco and Raúl A. Feijóo. A 3D-1D-0D Computational Model for the Entire Cardiovascular System. Mecánica Computacional, 29(59):5887–5911, 2010. URL https://cimec.org.ar/ojs/index.php/mc/article/view/3419. Number: 59.

[130] F. Regazzoni, M. Salvador, L. Dede’, and A. Quarteroni. A machine learning method for real-time numerical simulations of cardiac electromechanics. Computer Methods in Applied Mechanics and Engineering, 393: 114825, April 2022. ISSN 0045-7825. doi:10.1016/j.cma.2022.114825. URL https://www.sciencedirect.com/science/article/pii/S004578252200144X.

[131] Martin R. Pfaller, Jonathan Pham, Nathan M. Wilson, David W. Parker, and Alison L. Marsden. On the periodicity of cardiovascular fluid dynamics simulations. Annals of biomedical engineering, 49(12):3574–3592, December 2021. ISSN 0090-6964. doi:10.1007/s10439-021-02796-x. URL https://www.ncbi.nlm.nih.gov/pmc/articles/PMC9831274/.

[132] Alan Lazarus, David Dalton, Dirk Husmeier, and Hao Gao. Sensitivity analysis and inverse uncertainty quantification for the left ventricular passive mechanics. Biomechanics and Modeling in Mechanobiology, 21(3):953–982, June 2022. ISSN 1617-7940. doi:10.1007/s10237-022-01571-8. URL https://link.springer.com/article/10.1007/s10237-022-01571-8. Company: Springer Distributor: Springer Institution: Springer Label: Springer Number: 3 Publisher: Springer Berlin Heidelberg.

[133] Stefan Klotz, Ilan Hay, Marc L. Dickstein, Geng-Hua Yi, Jie Wang, Mathew S. Maurer, David A. Kass, and Daniel Burkhoff. Single-beat estimation of end-diastolic pressure-volume relationship: a novel method with potential for noninvasive application. American Journal of Physiology-Heart and Circulatory Physiology, 291 (1):H403–H412, July 2006. ISSN 0363-6135. doi:10.1152/ajpheart.01240.2005. URL https://journals.physiology.org/doi/full/10.1152/ajpheart.01240.2005. Publisher: American Physiological Society.

[134] Udo Sechtem, Barbara A. Sommerhoff, Walter Markiewicz, Richard D. White, Melvin D. Cheitlin, and Charles B. Higgins. Regional left ventricular wall thickening by magnetic resonance imaging: Evaluation in normal persons and patients with global and regional dysfunction. The American Journal of Cardiology, 59(1):145–151, January 1987. ISSN 0002-9149. doi:10.1016/S0002-9149(87)80088-7. URL https://www.sciencedirect.com/science/article/pii/S0002914987800887.

[135] Ronald M. Peshock, Roxann Rokey, Craig M. Malloy, Patrick McNamee, L. Maximilian Buja, Robert W. Parkey, and James T. Willerson. Assessment of myocardial systolic wall thickening using nuclear magnetic resonance imaging. Journal of the American College of Cardiology, 14(3):653–659, September 1989. ISSN 0735-1097. doi:10.1016/0735-1097(89)90106-X. URL https://www.sciencedirect.com/science/article/pii/073510978990106X.

[136] Sheng-Jing Dong, John H. MacGregor, Adrian P. Crawley, Elliot McVeigh, Israel Belenkie, Eldon R. Smith, John V. Tyberg, and Rafael Beyar. Left Ventricular Wall Thickness and Regional Systolic Function in Patients With Hypertrophic Cardiomyopathy. A Three-dimensional Tagged Magnetic Resonance Imaging Study. Circulation, 90(3):1200–1209, September 1994. ISSN 0009-7322. doi:10.1161/01.cir.90.3.1200. URL https://pmc.ncbi.nlm.nih.gov/articles/PMC2396316/.

[137] Charles Sillett, Orod Razeghi, Marina Strocchi, Caroline H. Roney, Hugh O’Brien, Daniel B. Ennis, Ulrike Haberland, Ronak Rajani, Christopher A. Rinaldi, and Steven A. Niederer. Optimisation of Left Atrial Feature Tracking Using Retrospective Gated Computed Tomography Images. In Daniel B. Ennis, Luigi E. Perotti, and Vicky Y. Wang, editors, Functional Imaging and Modeling of the Heart, pages 71–83, Cham, 2021. Springer International Publishing. ISBN 978-3-030-78710-3. doi:10.1007/978-3-030-78710-3_8.

[138] American Heart Association Writing Group on Myocardial Segmentation and Registration for Cardiac Imaging:, Manuel D. Cerqueira, Neil J. Weissman, Vasken Dilsizian, Alice K. Jacobs, Sanjiv Kaul, Warren K. Laskey, Dudley J. Pennell, John A. Rumberger, Thomas Ryan, and Mario S. Verani. Standardized Myocardial Segmentation and Nomenclature for Tomographic Imaging of the Heart. Circulation, 105(4):539–542, January 2002. doi:10.1161/hc0402.102975. URL https://www.ahajournals.org/doi/10.1161/hc0402.102975. Publisher: American Heart Association.

[139] Márton Tokodi, Levente Staub, Ádám Budai, Bálint Károly Lakatos, Máté Csákvári, Ferenc Imre Suhai, Liliána Szabó, Alexandra Fábián, Hajnalka Vágó, Zoltán Toşér, Béla Merkely, and Attila Kovács. Partitioning the Right Ventricle Into 15 Segments and Decomposing Its Motion Using 3D Echocardiography-Based Models: The Updated ReVISION Method. Frontiers in Cardiovascular Medicine, 8, March 2021. ISSN 2297-055X. doi:10.3389/fcvm.2021.622118. URL https://www.frontiersin.org/journals/cardiovascular-medicine/articles/10.3389/fcvm.2021.622118/full. Publisher: Frontiers.

[140] Chiara Corsini, Catriona Baker, Ethan Kung, Silvia Schievano, Gregory Arbia, Alessia Baretta, Giovanni Biglino, Francesco Migliavacca, Gabriele Dubini, Giancarlo Pennati, Alison Marsden, Irene Vignon-Clementel, Andrew Taylor, Tain-Yen Hsia, and Adam Dorfman. An integrated approach to patient-specific predictive modeling for single ventricle heart palliation. Computer Methods in Biomechanics and Biomedical Engineering, 17(14):1572–1589, October 2014. ISSN 1025-5842. doi:10.1080/10255842.2012.758254. URL https://doi.org/10.1080/10255842.2012.758254. Publisher: Taylor & Francis _eprint: https://doi.org/10.1080/10255842.2012.758254.

[141] Yunxiao Zhang, Moritz Kalhöfer-Köchling, Eberhard Bodenschatz, and Yong Wang. Physical model of enddiastolic and end-systolic pressure-volume relationships of a heart. Frontiers in Physiology, 14, August 2023. ISSN 1664-042X. doi:10.3389/fphys.2023.1195502. URL https://www.frontiersin.org/journals/physiology/articles/10.3389/fphys.2023.1195502/full. Publisher: Frontiers.

[142] Joost Lumens, Tammo Delhaas, Borut Kirn, and Theo Arts. Three-Wall Segment (TriSeg) Model Describing Mechanics and Hemodynamics of Ventricular Interaction. Annals of Biomedical Engineering, 37(11):2234–2255, November 2009. ISSN 1573-9686. doi:10.1007/s10439-009-9774-2. URL https://link.springer.com/article/10.1007/s10439-009-9774-2. Company: Springer Distributor: Springer Institution: Springer Label: Springer Number: 11 Publisher: Springer US.

[143] Marie Haghebaert, Pavlos Varsos, Roel Meiburg, and Irene Vignon-Clementel. A comparative study of lumped heart models for personalized medicine through sensitivity and identifiability analysis. The Journal of Physiology, n/a(n/a), May 2025. ISSN 1469-7793. doi:10.1113/JP287929. URL https://onlinelibrary.wiley.com/doi/abs/10.1113/JP287929. _eprint: https://physoc.onlinelibrary.wiley.com/doi/pdf/10.1113/JP287929.

[144] Socrates Dokos, Bruce H. Smaill, Alistair A. Young, and Ian J. LeGrice. Shear properties of passive ventricular myocardium. American Journal of Physiology-Heart and Circulatory Physiology, 283(6):H2650–H2659, December 2002. ISSN 0363-6135. doi:10.1152/ajpheart.00111.2002. URL https://journals.physiology.org/doi/full/10.1152/ajpheart.00111.2002. Publisher: American Physiological Society.

[145] F. C. Yin, R. K. Strumpf, P. H. Chew, and S. L. Zeger. Quantification of the mechanical properties of noncontracting canine myocardium under simultaneous biaxial loading. Journal of Biomechanics, 20(6):577–589, 1987. ISSN 0021-9290. doi:10.1016/0021-9290(87)90279-x.

[146] Debao Guan, Xiaoyu Luo, and Hao Gao. Constitutive Modelling of Soft Biological Tissue from Ex Vivo to in Vivo: Myocardium as an Example. In Takashi Suzuki, Clair Poignard, Mark Chaplain, and Vito Quaranta, editors, Methods of Mathematical Oncology, pages 3–14, Singapore, 2021. Springer. ISBN 978-981-16-4866-3. doi:10.1007/978-981-16-4866-3_1.

[147] Arnab Palit, Sunil K. Bhudia, Theodoros N. Arvanitis, Glen A. Turley, and Mark A. Williams. In vivo estimation of passive biomechanical properties of human myocardium. Medical & Biological Engineering & Computing, 56(9):1615–1631, September 2018. ISSN 1741-0444. doi:10.1007/s11517-017-1768-x. URL https://doi.org/10.1007/s11517-017-1768-x.

[148] H. Gao, W. G. Li, L. Cai, C. Berry, and X. Y. Luo. Parameter estimation in a Holzapfel–Ogden law for healthy myocardium. Journal of Engineering Mathematics, 95(1):231–248, December 2015. ISSN 1573-2703. doi:10.1007/s10665-014-9740-3. URL https://doi.org/10.1007/s10665-014-9740-3.

[149] J. M. Guccione, A. D. McCulloch, and L. K. Waldman. Passive Material Properties of Intact Ventricular Myocardium Determined From a Cylindrical Model. Journal of Biomechanical Engineering, 113(1):42–55, February 1991. ISSN 0148-0731. doi:10.1115/1.2894084. URL https://doi.org/10.1115/1.2894084.

[150] Vicky Y. Wang, H. I. Lam, Daniel B. Ennis, Brett R. Cowan, Alistair A. Young, and Martyn P. Nash. Modelling passive diastolic mechanics with quantitative MRI of cardiac structure and function. Medical Image Analysis, 13(5):773–784, October 2009. ISSN 1361-8415. doi:10.1016/j.media.2009.07.006. URL https://www.sciencedirect.com/science/article/pii/S1361841509000619.

[151] Jiahe Xi, Pablo Lamata, Steven Niederer, Sander Land, Wenzhe Shi, Xiahai Zhuang, Sebastien Ourselin, Simon G. Duckett, Anoop K. Shetty, C. Aldo Rinaldi, Daniel Rueckert, Reza Razavi, and Nic P. Smith. The estimation of patient-specific cardiac diastolic functions from clinical measurements. Medical Image Analysis, 17(2):133–146, February 2013. ISSN 1361-8415. doi:10.1016/j.media.2012.08.001. URL https://www.sciencedirect.com/science/article/pii/S1361841512001004.

[152] J van der Velden, L.J Klein, M van der Bijl, M.A.J.M Huybregts, W Stooker, J Witkop, L Eijsman, C.A Visser, F.C Visser, and G.J.M Stienen. Force production in mechanically isolated cardiac myocytes from human ventricular muscle tissue. Cardiovascular Research, 38(2):414–423, May 1998. ISSN 0008-6363. doi:10.1016/S0008-6363(98)00019-4. URL https://doi.org/10.1016/S0008-6363(98)00019-4.

[153] J van der Velden, L.J Klein, M van der Bijl, M.A.J.M Huybregts, W Stooker, J Witkop, L Eijsman, C.A Visser, F.C Visser, and G.J.M Stienen. Isometric tension development and its calcium sensitivity in skinned myocyte-sized preparations from different regions of the human heart. Cardiovascular Research, 42(3):706–719, June 1999. ISSN 0008-6363. doi:10.1016/S0008-6363(98)00337-X. URL https://doi.org/10.1016/S0008-6363(98)00337-X.

[154] Sander Land, So-Jin Park-Holohan, Nicolas P. Smith, Cristobal G. dos Remedios, Jonathan C. Kentish, and Steven A. Niederer. A model of cardiac contraction based on novel measurements of tension development in human cardiomyocytes. Journal of Molecular and Cellular Cardiology, 106:68–83, May 2017. ISSN 0022-2828. doi:10.1016/j.yjmcc.2017.03.008. URL https://www.sciencedirect.com/science/article/pii/S0022282817300639.

[155] WH Gaasch, JS Cole, A A Quinones, and JK Alexander. Dynamic determinants of letf ventricular diastolic pressure-volume relations in man. Circulation, 51(2):317–323, February 1975. doi:10.1161/01.CIR.51.2.317. URL https://www.ahajournals.org/doi/abs/10.1161/01.cir.51.2.317. Publisher: American Heart Association.

[156] Ashkan Abdollahi, Yoko Kato, Hooman Bakhshi, Vinithra Varadarajan, Omar Chehab, Ralph Zeitoun, Mohammad R. Ostovaneh, Colin O. Wu, Alain G. Bertoni, Sanjiv J. Shah, Bharath Ambale-Venkatesh, David A. Bluemke, and João A. C. Lima. Differential Stroke Volume between Left and Right Ventricles as a Predictor of Clinical Outcomes: The MESA Study. Radiology, 312(1):e232973, July 2024. ISSN 0033-8419. doi:10.1148/radiol.232973. URL https://pubs.rsna.org/doi/full/10.1148/radiol.232973. Publisher: Radiological Society of North America.

[157] Minji Kim, Seulgi You, Taeyang Ha, Tae Hee Kim, and Doo Kyoung Kang. Effect of papillary muscle and trabeculae on left ventricular function analysis via computed tomography: A cross-sectional study. Medicine, 102(46):e36106, November 2023. doi:10.1097/MD.0000000000036106. URL https://journals.lww.com/md-journal/fulltext/2023/11170/effect_of_papillary_muscle_and_trabeculae_on_left.119.aspx.

[158] E.W. Remme, A.A. Young, K.F. Augenstein, B. Cowan, and P.J. Hunter. Extraction and quantification of left ventricular deformation modes. IEEE Transactions on Biomedical Engineering, 51(11):1923–1931, November 2004. ISSN 1558-2531. doi:10.1109/TBME.2004.834283. URL https://ieeexplore.ieee.org/abstract/document/1344195.

[159] Wei Zhang, Charles S. Chung, Leonid Shmuylovich, and Sándor J. Kovács. Is left ventricular volume during diastasis the real equilibrium volume, and what is its relationship to diastolic suction? Journal of Applied Physiology, 105(3):1012–1014, September 2008. ISSN 8750-7587. doi:10.1152/japplphysiol.00799.2007. URL https://journals.physiology.org/doi/full/10.1152/japplphysiol.00799.2007. Publisher: American Physiological Society.

[160] Espen W. Remme, Anders Opdahl, and Otto A. Smiseth. Mechanics of left ventricular relaxation, early diastolic lengthening, and suction investigated in a mathematical model. American Journal of Physiology-Heart and Circulatory Physiology, 300(5):H1678–H1687, May 2011. ISSN 0363-6135. doi:10.1152/ajpheart.00165.2010. URL https://journals.physiology.org/doi/full/10.1152/ajpheart.00165.2010. Publisher: American Physiological Society.

[161] Aditya V. S. Ponnaluri, Ilya A. Verzhbinsky, Jeff D. Eldredge, Alan Garfinkel, Daniel B. Ennis, and Luigi E. Perotti. Model of Left Ventricular Contraction: Validation Criteria and Boundary Conditions. In Yves Coudière, Valéry Ozenne, Edward Vigmond, and Nejib Zemzemi, editors, Functional Imaging and Modeling of the Heart, pages 294–303, Cham, 2019. Springer International Publishing. ISBN 978-3-030-21949-9. doi:10.1007/978-3-030-21949-9_32.

[162] Kévin Moulin, Pierre Croisille, Magalie Viallon, Ilya A Verzhbinsky, Luigi E Perotti, and Daniel B. Ennis. Myofiber strain in healthy humans using DENSE and cDTI. Magnetic resonance in medicine, 86(1):277–292, July 2021. ISSN 0740-3194. doi:10.1002/mrm.28724. URL https://www.ncbi.nlm.nih.gov/pmc/articles/PMC8223515/.

[163] Eric D. Carruth, Andrew D. McCulloch, and Jeffrey H. Omens. Transmural gradients of myocardial structure and mechanics: Implications for fiber stress and strain in pressure overload. Progress in Biophysics and Molecular Biology, 122(3):215–226, December 2016. ISSN 0079-6107. doi:10.1016/j.pbiomolbio.2016.11.004. URL https://www.sciencedirect.com/science/article/pii/S0079610716301511.

[164] David J. Clancy, Anthony Mclean, Michel Slama, and Sam R. Orde. Paradoxical septal motion: A diagnostic approach and clinical relevance. Australasian Journal of Ultrasound in Medicine, 21(2):79–86, 2018. ISSN 2205-0140. doi:10.1002/ajum.12086. URL https://onlinelibrary.wiley.com/doi/abs/10.1002/ajum.12086. _eprint: https://onlinelibrary.wiley.com/doi/pdf/10.1002/ajum.12086.

[165] Cristina Méndez, Rafaela Soler, Esther Rodriguez, Marisol López, Lucia Álvarez, Noela Fernández, and Lorenzo Montserrat. Magnetic resonance imaging of abnormal ventricular septal motion in heart diseases: a pictorial review. Insights into Imaging, 2(4):483–492, April 2011. ISSN 1869-4101. doi:10.1007/s13244-011-0093-4. URL https://www.ncbi.nlm.nih.gov/pmc/articles/PMC3259355/.

[166] Shumpei Mori, Satoshi Nakatani, Hideaki Kanzaki, Kenichiro Yamagata, Yutaka Take, Yunosuke Matsuura, Shingo Kyotani, Norifumi Nakanishi, and Masafumi Kitakaze. Patterns of the interventricular septal motion can predict conditions of patients with pulmonary hypertension. Journal of the American Society of Echocardiography: Official Publication of the American Society of Echocardiography, 21(4):386–393, April 2008. ISSN 1097-6795. doi:10.1016/j.echo.2007.05.037.

[167] Myrianthi Hadjicharalambous, Liya Asner, Radomir Chabiniok, Eva Sammut, James Wong, Devis Peressutti, Eric Kerfoot, Andrew King, Jack Lee, Reza Razavi, Nicolas Smith, Gerald Carr-White, and David Nordsletten. Non-invasive Model-Based Assessment of Passive Left-Ventricular Myocardial Stiffness in Healthy Subjects and in Patients with Non-ischemic Dilated Cardiomyopathy. Annals of Biomedical Engineering, 45(3):605– 618, March 2017. ISSN 1573-9686. doi:10.1007/s10439-016-1721-4. URL https://link.springer.com/article/10.1007/s10439-016-1721-4. Company: Springer Distributor: Springer Institution: Springer Label: Springer Number: 3 Publisher: Springer US.

[168] Debao Guan, Jiang Yao, Xiaoyu Luo, and Hao Gao. Effect of myofibre architecture on ventricular pump function by using a neonatal porcine heart model: from DT-MRI to rule-based methods. Royal Society Open Science, 7 (4):191655, April 2020. doi:10.1098/rsos.191655. URL https://royalsocietypublishing.org/doi/10.1098/rsos.191655. Publisher: Royal Society.

[169] Marina Strocchi, Christoph M. Augustin, Matthias A. F. Gsell, Elias Karabelas, Aurel Neic, Karli Gillette, Caroline H. Roney, Orod Razeghi, Jonathan M. Behar, Christopher A. Rinaldi, Edward J. Vigmond, Martin J. Bishop, Gernot Plank, and Steven A. Niederer. The Effect of Ventricular Myofibre Orientation on Atrial Dynamics. In Daniel B. Ennis, Luigi E. Perotti, and Vicky Y. Wang, editors, Functional Imaging and Modeling of the Heart, pages 659–670, Cham, 2021. Springer International Publishing. ISBN 978-3-030-78710-3. doi:10.1007/978-3-030-78710-3_63.

[170] Rocío Rodríguez-Cantano, Joakim Sundnes, and Marie E. Rognes. Uncertainty in cardiac myofiber orientation and stiffnesses dominate the variability of left ventricle deformation response. International Journal for Numerical Methods in Biomedical Engineering, 35(5):e3178, 2019. ISSN 2040-7947. doi:10.1002/cnm.3178. URL https://onlinelibrary.wiley.com/doi/abs/10.1002/cnm.3178. _eprint: https://onlinelibrary.wiley.com/doi/pdf/10.1002/cnm.3178.

[171] Dennis Ogiermann, Daniel Balzani, and Luigi E. Perotti. Analyzing the Impact of Different Microstructure and Active Stress Models on Peak Systolic Kinematics. In Radomír Chabiniok, Qing Zou, Tarique Hussain, Hoang H. Nguyen, Vlad G. Zaha, and Maria Gusseva, editors, Functional Imaging and Modeling of the Heart, pages 305–318, Cham, 2025. Springer Nature Switzerland. ISBN 978-3-031-94559-5. doi:10.1007/978-3-03194559-5_28.

[172] Takumi Washio, Seiryo Sugiura, Jun-ichi Okada, and Toshiaki Hisada. Using Systolic Local Mechanical Load to Predict Fiber Orientation in Ventricles. Frontiers in Physiology, 11, June 2020. ISSN 1664-042X. doi:10.3389/fphys.2020.00467. URL https://www.frontiersin.org/journals/physiology/articles/10.3389/fphys.2020.00467/full. Publisher: Frontiers.

[173] Kasra Osouli, Francesco De Gaetano, Maria Laura Costantino, and Mathias Peirlinck. Heart in a knot: unraveling the impact of the nested tori myofiber architecture on ventricular mechanics. Biomechanics and Modeling in Mechanobiology, 24(5):1815–1835, October 2025. ISSN 1617-7940. doi:10.1007/s10237-025-01995-y. URL https://doi.org/10.1007/s10237-025-01995-y.

[174] Nicolas Toussaint, Christian T. Stoeck, Tobias Schaeffter, Sebastian Kozerke, Maxime Sermesant, and Philip G. Batchelor. In vivo human cardiac fibre architecture estimation using shape-based diffusion tensor processing. Medical Image Analysis, 17(8):1243–1255, December 2013. ISSN 1361-8415. doi:10.1016/j.media.2013.02.008. URL https://www.sciencedirect.com/science/article/pii/S1361841513000224.

[175] Vallespin Sonia Nielles, Zohya Khalique, Pedro F. Ferreira, Silva Ranil de, Andrew D. Scott, Philip Kilner, Laura-Ann McGill, Archontis Giannakidis, Peter D. Gatehouse, Daniel Ennis, Eric Aliotta, Khalil Majid Al, Peter Kellman, Dumitru Mazilu, Robert S. Balaban, David N. Firmin, Andrew E. Arai, and Dudley J. Pennell. Assessment of Myocardial Microstructural Dynamics by In Vivo Diffusion Tensor Cardiac Magnetic Resonance. Journal of the American College of Cardiology, 69(6):661–676, February 2017. doi:10.1016/j.jacc.2016.11.051. URL https://www.jacc.org/doi/full/10.1016/j.jacc.2016.11.051. Publisher: American College of Cardiology Foundation.

[176] Kévin Moulin, Ilya A. Verzhbinsky, Nyasha G. Maforo, Luigi E. Perotti, and Daniel B. Ennis. Probing cardiomyocyte mobility with multi-phase cardiac diffusion tensor MRI. PLOS ONE, 15(11):e0241996, November 2020. ISSN 1932-6203. doi:10.1371/journal.pone.0241996. URL https://journals.plos.org/plosone/article?id=10.1371/journal.pone.0241996. Publisher: Public Library of Science.

[177] Kellie Phipps, Maaike van de Boomen, Robert Eder, Sam Allen Michelhaugh, Aferdita Spahillari, Joan Kim, Shestruma Parajuli, Timothy G. Reese, Choukri Mekkaoui, Saumya Das, Denise Gee, Ravi Shah, David E. Sosnovik, and Christopher Nguyen. Accelerated in Vivo Cardiac Diffusion-Tensor MRI Using Residual Deep Learning–based Denoising in Participants with Obesity. Radiology: Cardiothoracic Imaging, 3(3):e200580, June 2021. doi:10.1148/ryct.2021200580. URL https://pubs.rsna.org/doi/full/10.1148/ryct.2021200580. Publisher: Radiological Society of North America.

[178] Alexander J. Wilson, Gregory B. Sands, Ian J. LeGrice, Alistair A. Young, and Daniel B. Ennis. Myocardial mesostructure and mesofunction. American Journal of Physiology-Heart and Circulatory Physiology, 323(2): H257–H275, August 2022. ISSN 0363-6135. doi:10.1152/ajpheart.00059.2022. URL https://journals.physiology.org/doi/full/10.1152/ajpheart.00059.2022. Publisher: American Physiological Society.

[179] Cristobal Rodero, Tiffany M. G. Baptiste, Rosie K. Barrows, Alexandre Lewalle, Steven A. Niederer, and Marina Strocchi. Advancing clinical translation of cardiac biomechanics models: a comprehensive review, applications and future pathways. Frontiers in Physics, 11, November 2023. ISSN 2296-424X. doi:10.3389/fphy.2023.1306210. URL https://www.frontiersin.org/journals/physics/articles/10.3389/fphy.2023.1306210/full. Publisher: Frontiers.

[180] Johanna Stimm, David A. Nordsletten, Javiera Jilberto, Renee Miller, Ezgi Berberoğlu, Sebastian Kozerke, and Christian T. Stoeck. Personalization of biomechanical simulations of the left ventricle by in-vivo cardiac DTI data: Impact of fiber interpolation methods. Frontiers in Physiology, 13, November 2022. ISSN 1664-042X. doi:10.3389/fphys.2022.1042537. URL https://www.frontiersin.org/journals/physiology/articles/10.3389/fphys.2022.1042537/full. Publisher: Frontiers.

[181] S. R. Ommen, R. A. Nishimura, C. P. Appleton, F. A. Miller, J. K. Oh, M. M. Redfield, and A. J. Tajik. Clinical Utility of Doppler Echocardiography and Tissue Doppler Imaging in the Estimation of Left Ventricular Filling Pressures. Circulation, 102(15):1788–1794, October 2000. doi:10.1161/01.CIR.102.15.1788. URL https://www.ahajournals.org/doi/full/10.1161/01.CIR.102.15.1788. Publisher: American Heart Association.

[182] Nandan S. Anavekar and Jae K. Oh. Doppler echocardiography: A contemporary review. Journal of Cardiology, 54(3):347–358, December 2009. ISSN 0914-5087. doi:10.1016/j.jjcc.2009.10.001. URL https://www.sciencedirect.com/science/article/pii/S0914508709002731.

[183] A Bistoquet, J Oshinski, and O Skrinjar. Myocardial deformation recovery from cine MRI using a nearly incompressible biventricular model. Medical Image Analysis, 12(1):69–85, February 2008. ISSN 13618415. doi:10.1016/j.media.2007.10.009. URL https://linkinghub.elsevier.com/retrieve/pii/S1361841507001090.

[184] Christopher C. Moore, Carlos H. Lugo-Olivieri, Elliot R. McVeigh, and Elias A. Zerhouni. Three-dimensional Systolic Strain Patterns in the Normal Human Left Ventricle: Characterization with Tagged MR Imaging. Radiology, 214(2):453–466, February 2000. ISSN 0033-8419. doi:10.1148/radiology.214.2.r00fe17453. URL https://pubs.rsna.org/doi/full/10.1148/radiology.214.2.r00fe17453. Publisher: Radiological Society of North America.

[185] Jing Liu, Ruhul Sarker, Saber Elsayed, Daryl Essam, and Nurhadi Siswanto. Large-scale evolutionary optimization: A review and comparative study. Swarm and Evolutionary Computation, 85:101466, March 2024. ISSN 2210-6502. doi:10.1016/j.swevo.2023.101466. URL https://www.sciencedirect.com/science/article/pii/S2210650223002389.

[186] N. A. Barnafi, F. Regazzoni, and D. Riccobelli. Reconstructing relaxed configurations in elastic bodies: Mathematical formulations and numerical methods for cardiac modeling. Computer Methods in Applied Mechanics and Engineering, 423:116845, April 2024. ISSN 0045-7825. doi:10.1016/j.cma.2024.116845. URL https://www.sciencedirect.com/science/article/pii/S0045782524001014.

[187] Fatemeh Fatemifar, Marc D. Feldman, Geoffrey D. Clarke, Ender A. Finol, and Hai-Chao Han. Computational Modeling of Human Left Ventricle to Assess the Effects of Trabeculae Carneae on the Diastolic and Systolic Functions. Journal of Biomechanical Engineering, 141(091014), August 2019. ISSN 0148-0731. doi:10.1115/1.4043831. URL https://doi.org/10.1115/1.4043831.

[188] Marta Serrani, Maria Laura Costantino, and Roberto Fumero. The influence of cardiac trabeculae on ventricular mechanics. In Computing in Cardiology 2013, pages 811–814, September 2013. URL https://ieeexplore.ieee.org/abstract/document/6713501. ISSN: 2325-8853.

[189] Vijay Vedula, Jung-Hee Seo, Albert C. Lardo, and Rajat Mittal. Effect of trabeculae and papillary muscles on the hemodynamics of the left ventricle. Theoretical and Computational Fluid Dynamics, 30(1):3–21, April 2016. ISSN 1432-2250. doi:10.1007/s00162-015-0349-6. URL https://doi.org/10.1007/s00162-015-0349-6.

[190] E. Rene Rodriguez and Carmela D. Tan. Structure and Anatomy of the Human Pericardium. Progress in Cardiovascular Diseases, 59(4):327–340, January 2017. ISSN 0033-0620. doi:10.1016/j.pcad.2016.12.010. URL https://www.sciencedirect.com/science/article/pii/S0033062016301463.

[191] V. Gurev, J. Constantino, J.J. Rice, and N.A. Trayanova. Distribution of Electromechanical Delay in the Heart: Insights from a Three-Dimensional Electromechanical Model. Biophysical Journal, 99(3):745–754, August 2010. ISSN 0006-3495. doi:10.1016/j.bpj.2010.05.028. URL https://www.ncbi.nlm.nih.gov/pmc/articles/PMC2913183/.

[192] J.S.S.G de Jong. Conduction - ECGpedia, September 2021. URL https://en.ecgpedia.org/index.php?title=Conduction&oldid=16965%22.

